# Granzyme-targeting quenched activity-based probes for assessing tumor response to immunotherapy

**DOI:** 10.1101/2025.03.13.643086

**Authors:** Muhammad Kazim, Arghya Ganguly, Sebastian M. Malespini, Lai Thang, Nimit L. Patel, Caleb Kim, Joseph D. Kalen, Simone Difilippantonio, Euna Yoo

## Abstract

Molecular imaging of immune activation holds tremendous potential for the development of novel immunotherapy. In particular, chemical probes capable of detecting immune responses before changes in tumor size occur can guide early therapeutic strategies. Here, we present quenched activity-based probes targeting granzymes as a biomarker of antitumor immunity. Through optimization of peptide recognition element and functional chemical warhead, we have developed an optical imaging probe Cy5-IEPCya^PhP^-QSY21, which rapidly reacts with GzmB at substoichiometric concentrations and enables efficient, selective labeling of the active enzyme in a complex proteome. With high specificity and minimal background signal, this probe produces GzmB-induced near-infrared fluorescence signals in the tumors of living mice shortly after injection. Both in vivo and ex vivo fluorescence signals correlate with GzmB expression and activity, and the population of CD8+ cells in tumor tissues. Moreover, it demonstrates the potential to track tumor response to immunotherapy. Thus, this study offers a chemical tool for assessing immune-mediated anticancer activity using noninvasive optical imaging.

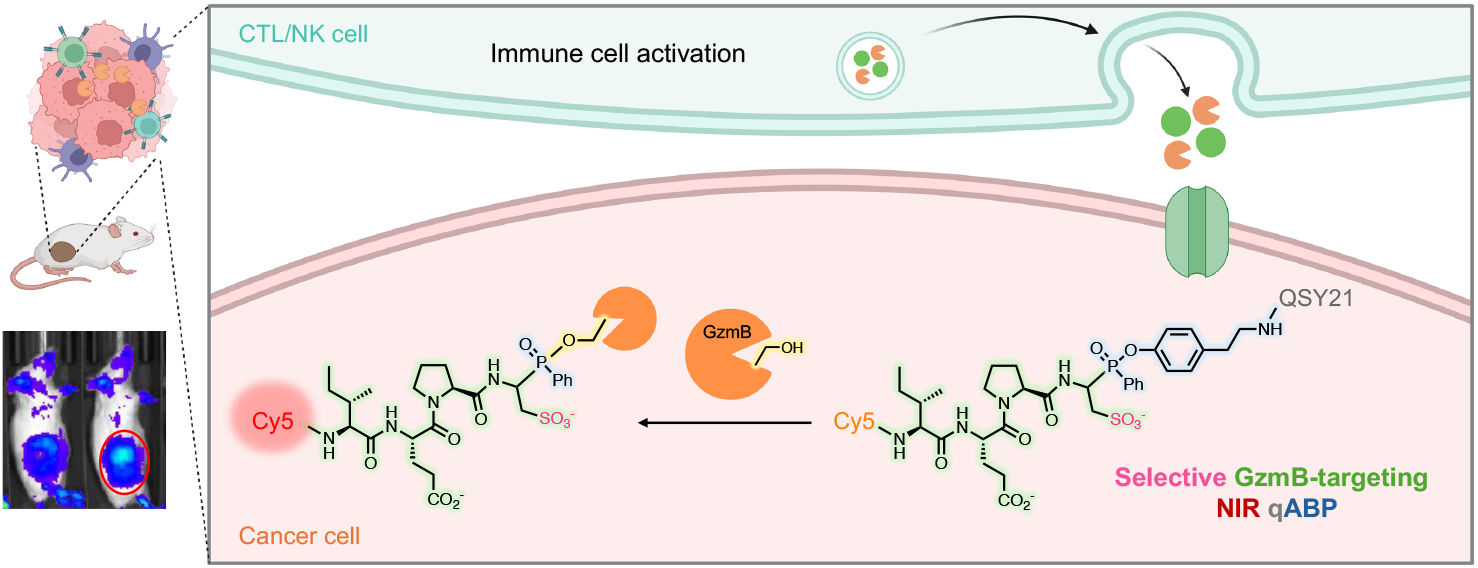

## Introduction

In recent decades, the field of tumor immunology has witnessed a major revolution, including significant clinical progress with various immunotherapeutic approaches.^1-5^ While cancer immunotherapies such as immune checkpoint blockade (ICB) and T cell-based therapies have demonstrated great clinical success, only a subpopulation of patients respond to the treatment, calling for reliable biomarkers and therapeutic strategies to maximize the benefits of immunotherapy.^6^ The critical component of successful immunotherapy is the effective infiltration and activation of immune cells, especially cytotoxic T lymphocytes (CTLs) and natural killer (NK) cells, in the tumor microenvironment (TME). Methods detecting expression of cell surface markers or secretion of cytokines of these immune cells to assess such events within the antitumor immunity cycle, which largely rely on affinity-based measurements using antibodies, require tedious steps for sample preparation and processing and are unsuitable for real-time detection.^7-10^ On the other hand, real-time functional readout of immune-mediated anticancer action has the potential to provide a more accurate assessment and identify optimal treatments at the early stage.

Molecular imaging is a rapidly growing platform for visualizing biological activities in living organisms. In the tumor immune microenvironment (TIME), imaging probes targeting functional enzymes can provide insight into immune cell dynamics, tumor characteristics, and treatment response, enabling informed decisions for therapeutic strategy. In vivo optical imaging that leverages the enzymatic turnover of substrates to products with signal amplification over time has shown variable success in detecting immune activation.^11-13^ Activity-based probes (ABPs) harnessing electrophilic warheads, on the other hand, use a covalent reaction with the catalytic residue to report the activity of enzymes.^14, 15^ The irreversible attachment to a target protein can reduce rapid diffusion, thus increasing signal retention at the site. Additionally, it allows secondary, ex vivo profiling to identify the targeted enzymes and to correlate their activity with in vivo images. Recently, there has been increased interest in developing activatable probes with near-infrared (NIR) fluorophores for various clinical applications. As the excitation and emission wavelengths of NIR fluorophores are above the wavelengths of auto-fluorescent background from living tissues, NIR fluorescence (NIRF) imaging can achieve increased signal-to-background ratios (SBRs) with deeper tissue penetration (can penetrate up to 1 cm of tissue).^16-18^ Compared to ‘always on’ probes, the ‘activatable’ nature further enhances SBRs and requires shorter times for generating high contrast at the site of interest.^19, 20^ A suite of NIR fluorophore-labeled quenched ABPs (qABPs) for tumor-specific cathepsins, which are intrinsically nonfluorescent but produce a fluorescent signal upon nucleophilic displacement of the quencher group by the active site cysteine of the proteolytically active cathepsin, has been developed for fluorescence-guided surgery of tumors.^21, 22^ However, one of the major challenges in developing optical probes targeting functional enzymes is achieving a high degree of selectivity and stability to ensure accurate activity measurement in complex biological environments such as TIME.

Granzymes (Gzms) are serine proteases stored primarily as inactive zymogens inside CTLs and NK cells.^23^ Upon recognition of cancerous cells, Gzms are secreted along with pore-forming perforin that facilitates protein entry into target cells to initiate programmed cell death, which is the critical feature of immune effector processes.^24, 25^ Therefore, measuring the activity of Gzms in the TME can provide a direct and functional readout of the immune response. Among the five human Gzms (A, B, H, K, and M), the most studied GzmB is responsible for the lion’s share of immune cell-mediated cytotoxicity.^26^ Prior studies have implemented various molecular imaging strategies, including magnetic resonance and nuclear,^27-29^ fluorescence,^30-34^ luminescence,^35, 36^ and photoacoustic imaging,^37^ to measure GzmB activity. GzmB activity has typically been assessed with proteolytic cleavage of a canonical IEPD sequence. While an important substrate for the class, it displays cross-reactivity with caspase-8 (Casp-8) and its kinetic parameters are not optimal for in vivo imaging. We hypothesized that subtle tuning of the critical P1 residue, a negatively charged carboxyl functional group, could fundamentally alter this specificity. When coupled with an optimized covalent warhead, this new sequence provides the means to precisely measure GzmB activity with high signal contrast in living organisms.

Here, we report a GzmB-targeting qABP and exploit its utility for real-time in vivo imaging of immunotherapy responses. The incorporation of cysteic acid (Cya) at the P1 position of tetrapeptide resulted in enhanced binding and specificity for GzmB compared to an existing probe, allowing for efficient and selective labeling of the enzyme in cells. Leveraging the single phenoxy leaving group of the serine reactive phenyl phosphinate ester warhead, we generated a qABP containing a pair of NIR fluorophore and quencher that produces a fluorescence signal only upon reaction with active GzmB. The optimized probe exhibits minimal background noise, excellent hydrophilicity while maintaining cell permeability, and fast reaction kinetics with GzmB. Following intravenous injection, the probe was found to quickly generate and accumulate a fluorescence signal in the tumor of living animals that is correlated with the expression level of GzmB and CD8 and therapy response at the early stage of treatment, enabling real-time detection of immune responses.

## Results and Discussion

### Design of selective GzmB-targeting probes

It is well established that GzmB hydrolyzes peptides containing aspartic acid at the P1 position almost exclusively.^38-40^ This preference, however, overlaps with cysteine-dependent aspartate-specific proteases (caspases, Casp), making the peptide substrates nonspecific. Both human and mouse GzmB cleave many of the same substrates as caspases, including PARP-1, lamin B, NuMa, DNA-PKcs, and tubulin, and exhibit substrate specificity similar to that of Casp-6, -8, and -9.^41-43^ Although the tetrapeptide sequence IEPD has been reported as optimal for GzmB and utilized in most GzmB probes, it suffers cross-reactivity with Casp-8, which limits its utility.^44^ Therefore, for the design of GzmB-selective probes, we first set out to optimize the GzmB-targeting peptide to enhance binding and selectivity. To examine the effects of isosteres, we synthesized P1/P3 derivatives of IEPD, substituting the carboxylic acid side chain with sulfonic acid in the form of fluorogenic substrates using 7-amino-4-carbamoylmethylcoumarin (ACC) fluorophore (**1**-**4** in Figure 1A, Scheme S1). Incubation with recombinant human GzmB revealed that the tetrapeptide containing cysteic acid (Cya) at the P1 position (Ac-IEPCya-ACC, **2**) is cleaved more efficiently (6-fold) by GzmB compared to Ac-IEPD-ACC (**1**) (Figure 1B). More importantly, cleavage by Casp-8 was reduced. While P3 substitution (**3**) displayed an opposite preference, showing increased cleavage by Casp-8 but reduced cleavage by GzmB, substituting both P1 and P3 positions (**4**) routed for cleavage by GzmB while concurrently decreasing cross-reactivity with Casp-8. Kinetic analysis confirmed that Ac-IEPCya-ACC (**2**) exhibits a better binding (K_M_ = 385 μM), a 10-fold increase in turnover (k_cat_ = 0.338 s^-1^), and catalytic efficiency (k_cat_/K_M_ = 879 M^-1^s^-1^) for GzmB compared to Ac-IEPD-ACC (**1**), alongside reduced cross-reactivity with Casp-8 (Figure 1C). These data indicate that the substitution of carboxylic acid with sulfonic acid at P1 significantly enhances both binding affinity and selectivity toward GzmB.

**Figure 1.**
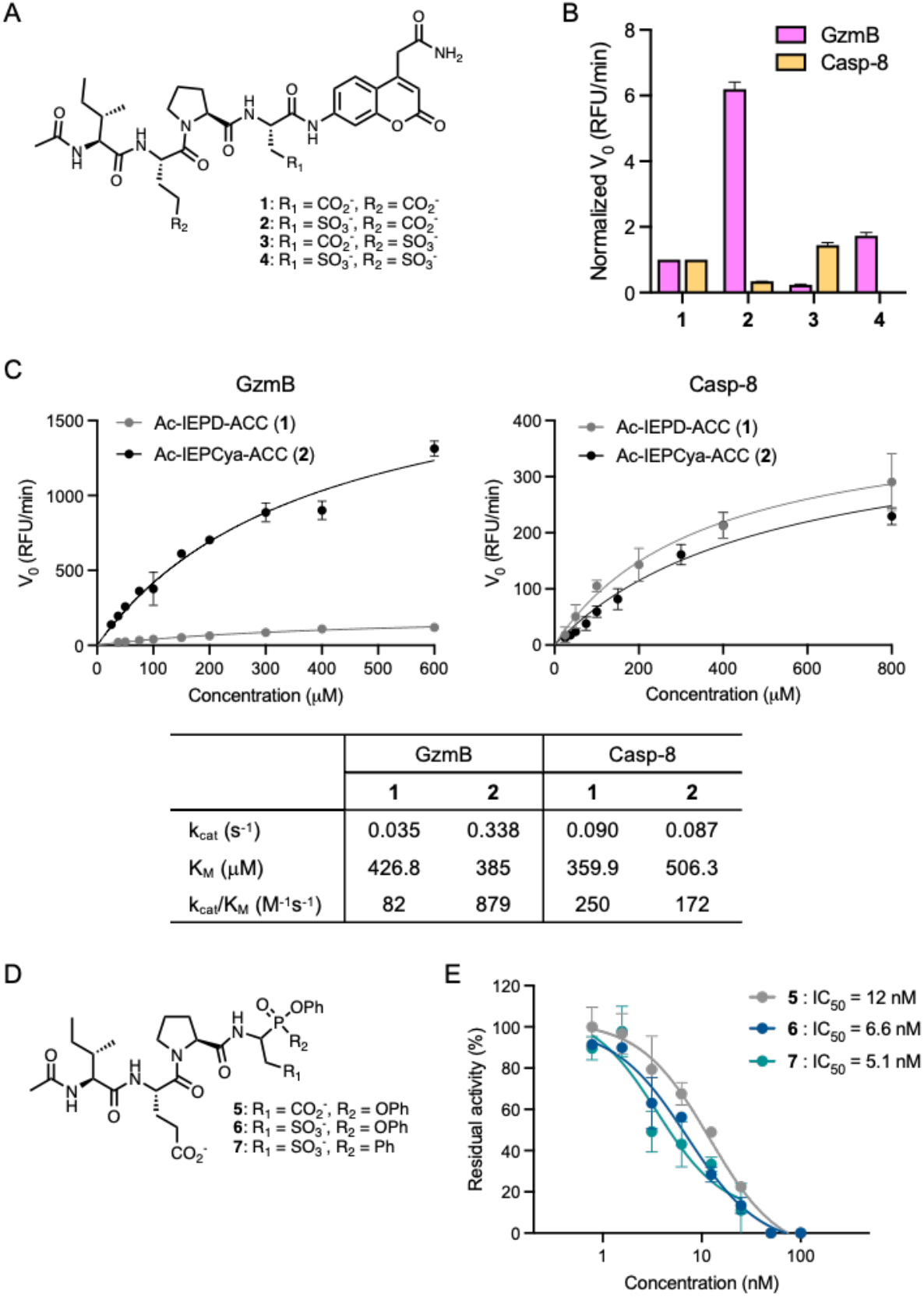
Isosteric substitution of carboxylic acid at P1 with sulfonic acid enhances selectivity for GzmB over Casp-8. (A) Chemical structures of tetrapeptide fluorogenic substrates **1**-**4**. (B) Enzyme cleavage assay. 100 μM of each substrate was incubated with either recombinant human active GzmB (10 nM) or Casp-8 (5 nM) and fluorescence was measured at Ex/Em: 380/460 nm for 1 h. The initial velocity (V_0_) of each substrate was normalized to that of the control substrate Ac-IEPD-ACC (**1**). Data points are displayed as mean ± SD (n = 3). (C) Kinetic measurement. The Michaelis-Menten plot and kinetic parameters were generated with varying concentrations of each substrate **1** and **2** incubated with recombinant human active GzmB (10 nM) and Casp-8 (5 nM). Data points are displayed as mean ± SD (n = 3). (D) Chemical structures of inhibitors **5**-**7**. (E) GzmB inhibition. Recombinant human active GzmB (2 U/μL) was incubated with varying concentrations of each inhibitor and the residual activity of GzmB was measured by the cleavage of Ac-IEPD-AMC fluorogenic substrate and normalized to DMSO control. Data points are displayed as mean ± SD (n = 3).

Using the IEPCya sequence, we converted the fluorogenic substrate into a covalent inhibitor featuring a serine-reactive diphenyl phosphonate warhead (Ac-IEPCya-PO(OPh)_2_, **6**, Figure 1D). We treated 2-(tritylthio)acetaldehyde (**I-1**, Scheme S2) with benzyl carbamate and triphenylphosphite in a Kabachnik-Fields type reaction to form the diphenyl phosphonate warhead core (**I-2**), which, after Trt-deprotection and oxidation of the free thiol, yielded the desired sulfonic acid-containing intermediate Cbz-Cya-PO(OPh)_2_ (**I-3**). Following the deprotection of the Cbz group, we coupled it with N-acetylated tripeptide Ac-IE(tBu)P-OH (**I-8**) and sub-sequently deprotected the tBu group to produce **6**. As expected, compound **6** potently inhibited GzmB activity with an IC_50_ value of 6.6 nM (Figure 1E). Given recent studies reporting phosphinate esters as viable warheads for serine proteases,^45-47^ we were intrigued to test whether phenyl phosphinate warheads would function well with GzmB. Therefore, we synthesized the phenyl phosphinate inhibitor **7** (Figure 1D) using dichlorophenylphosphine and phenol (Scheme S2). The inhibition of GzmB by Ac-IEPCya-PO(OPh)(Ph) (**7**) was comparable to that of **6**, with an IC_50_ value of 5.1 nM (Figure 1E).

Having verified that the P1 Cya performs better than the P1 Asp in GzmB substrates and inhibitors, we prepared the corresponding fluorescent ABPs **8** and **9**, which contain the NIR sulfo-Cy5 fluorophore to profile GzmB activity (Figure 2A and Scheme S3). Additionally, leveraging the single phenoxy leaving group of the phosphinate ester,^45^ we synthesized a qABP **10** featuring the sulfo-Cy5 and QSY21 (Figures 2B-C) where QSY21 quenches the fluorescence of Cy5 through Förster resonance energy transfer (FRET). 2-(tritylthio)acetaldehyde (**I-1**), benzyl carbamate, and dichlorophenylphosphine were reacted in a Mannich-type three-component reaction. The resulting intermediate was trapped with N-Alloc-protected tyramine leaving group (**I-12**) via Steglich-type esterification, followed by Trt-deprotection and oxidation to furnish the P1 Cya phenyl phosphinate building block (**I-14**). Upon selective Cbz-deprotection, a coupling reaction with the tripeptide Boc-IE(tBu)P-OH and subsequent Boc-deprotection produced the tetrapeptide core with a free amine that allowed for coupling with the sulfo-Cy5. Finally, the tyramine part was coupled with the QSY21 quencher after Alloc-deprotection to generate Cy5-IEPCya^PhP^-QSY21 **10**. When we measured the fluorescence signal of each probe at Ex/Em: 640/670 nm, our qABP **10** exhibited negligible Cy5 fluorescence, indicating efficient quenching by QSY21 (Figure S1A). We anticipated that the reaction with active GzmB would covalently conjugate the molecule to the catalytic serine within the active site, separate the quencher, and generate the fluorescent signal only from the labeled enzyme (Figure 2D). This ‘activatable’ ABP can minimize background signals and increase specificity compared to always-on probes.^48^ To assess the hydrolytic stability, we incubated qABP **10** in different media and measured the Cy5 fluorescence signal over time (Figure S1B). Although less stable in cell culture media compared to PBS, less than 50% of the hydrolyzed product was observed even after 24 h incubation. We then confirmed that our qABP **10** efficiently labels active human GzmB, but not the pro-form of the enzyme, at low nanomolar concentrations in a dose-dependent manner (Figure 2E). Compared to the always-on ABPs **8** and **9**, qABP **10** displayed slightly weaker labeling, suggesting that the incorporated quencher QSY21 may affect the probe’s initial binding. The observed labeling occurred through covalent conjugation, as the compound lacking the reactive warhead (**11**, Figure S2A) showed no labeling, even at 500 nM (Figure S2B). Furthermore, **10** was also capable of labeling mouse GzmB, although it required slightly higher concentrations, corroborating previous studies indicating that mouse and human GzmB have slightly different substrate specificity (IEFD and IEPD, respectively)^49^ (Figure S3A). Preincubation with inhibitor **6** or **7** blocked the GzmB labeling by ABPs, demonstrating that these compounds competitively bind to the active site of the enzyme (Figures S3B-C).

**Figure 2.**
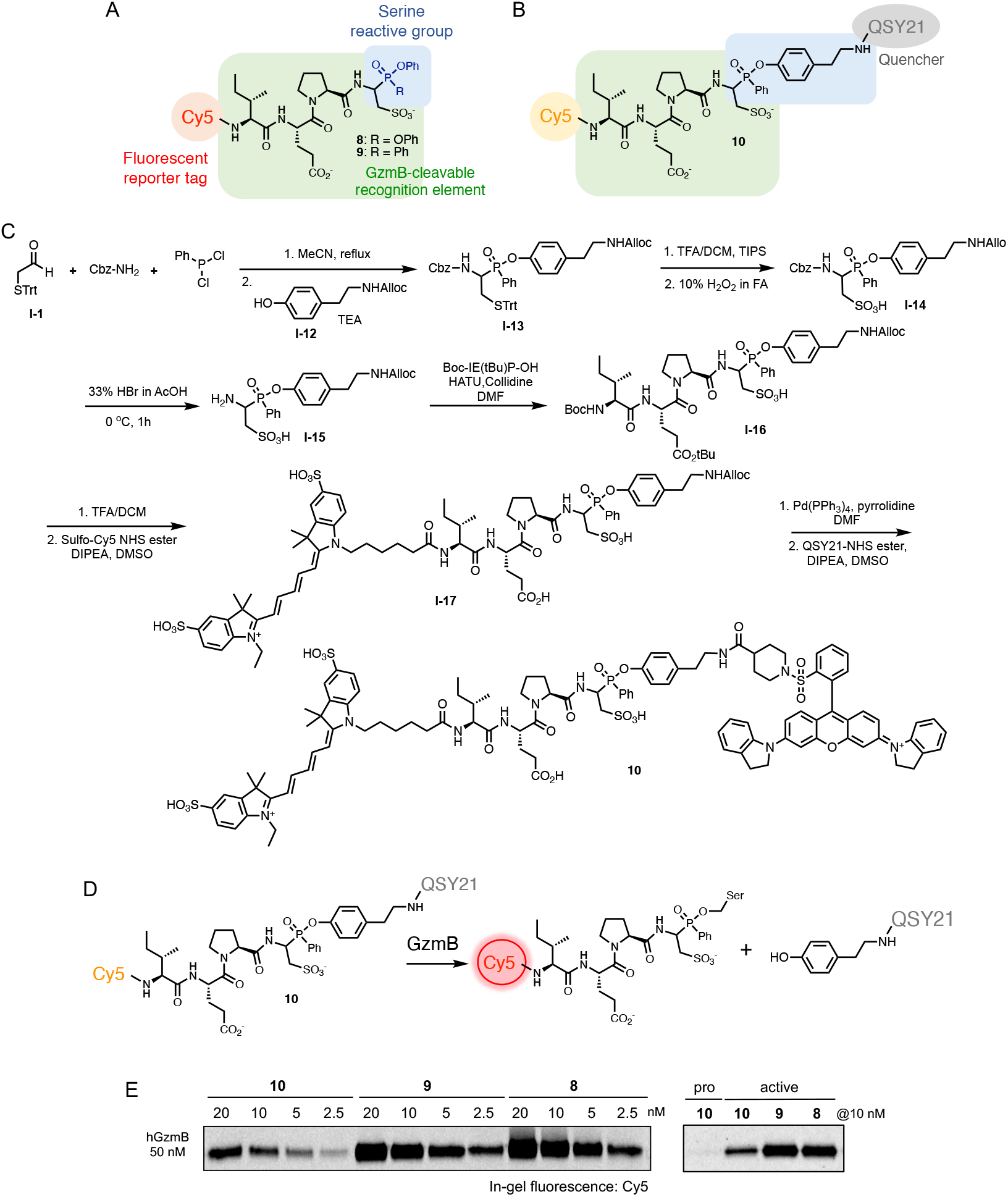
GzmB ABPs covalently label active GzmB. (A) Chemical structures of ABPs **8** and **9**. (B-C) Chemical structure and synthesis of qABP **10**. (D) Covalent reaction mechanism of qABP **10**. (E) GzmB labeling by ABPs. Recombinant human active GzmB (50 nM) was incubated with varying concentrations of each ABP for 1 h. Protein samples were analyzed by SDS-PAGE and in-gel fluorescence scanning for Cy5 signal.

### Activity-based GzmB profiling

To evaluate the efficacy and selectivity of our probes detecting GzmB activity in complex biological systems, we performed activity-based protein profiling (ABPP). Gel-based ABPP with probes **8**-**10**, performed in vitro (Figure S4A) and in situ (Figure 3A), demonstrated that these ABPs are efficiently and selectively conjugated to active GzmB. qABP **10** showed stronger labeling than **8** or **9** when intact NK-92 cells (immortalized human NK cell lines) were treated, likely due to an additional hydrophobic quencher in the molecule that enhances cell permeability. Pretreatment of NK-92 cell lysates with inhibitor **6** or **7** significantly blocked GzmB labeling by ABPs (Figure S4B). No apparent protein conjugate was observed in cells lacking GzmB expression (MDA-MB-231, Jurkat, and HEK293T cells), indicating that these ABPs are highly selective towards GzmB (Figure 3B). Similarly, when NK-92 cells treated with qABP **10** were analyzed via flow cytometry, we observed a time- and dose-dependent increase in fluorescence intensity (Figure S5A), approximately 9.1-fold greater than the signal from the GzmB non-expressing triple-negative breast cancer (TNBC) MDA-MB-231 cells at 5 μM (Figure 3C and Figure S5B). Compared to the always-on ABP **9**, which showed a signal-to-noise (S/N) value of 1.53, these results illustrate that our qABP reports GzmB activity in cells with high sensitivity and low background signal. To detect GzmB transferred to target cells from effector immune cells upon target cell recognition, we co-cultured NK-92 cells with MDA-MB-231 cells in the presence of qABP **10** (5 μM). After removing the suspended NK-92 cells, adherent MDA-MB-231 cells were harvested and subjected to either gel-based or flow cytometry analysis, where we detected the time-dependent transfer of GzmB labeled by the probe (Figure 3D-E).

**Figure 3.**
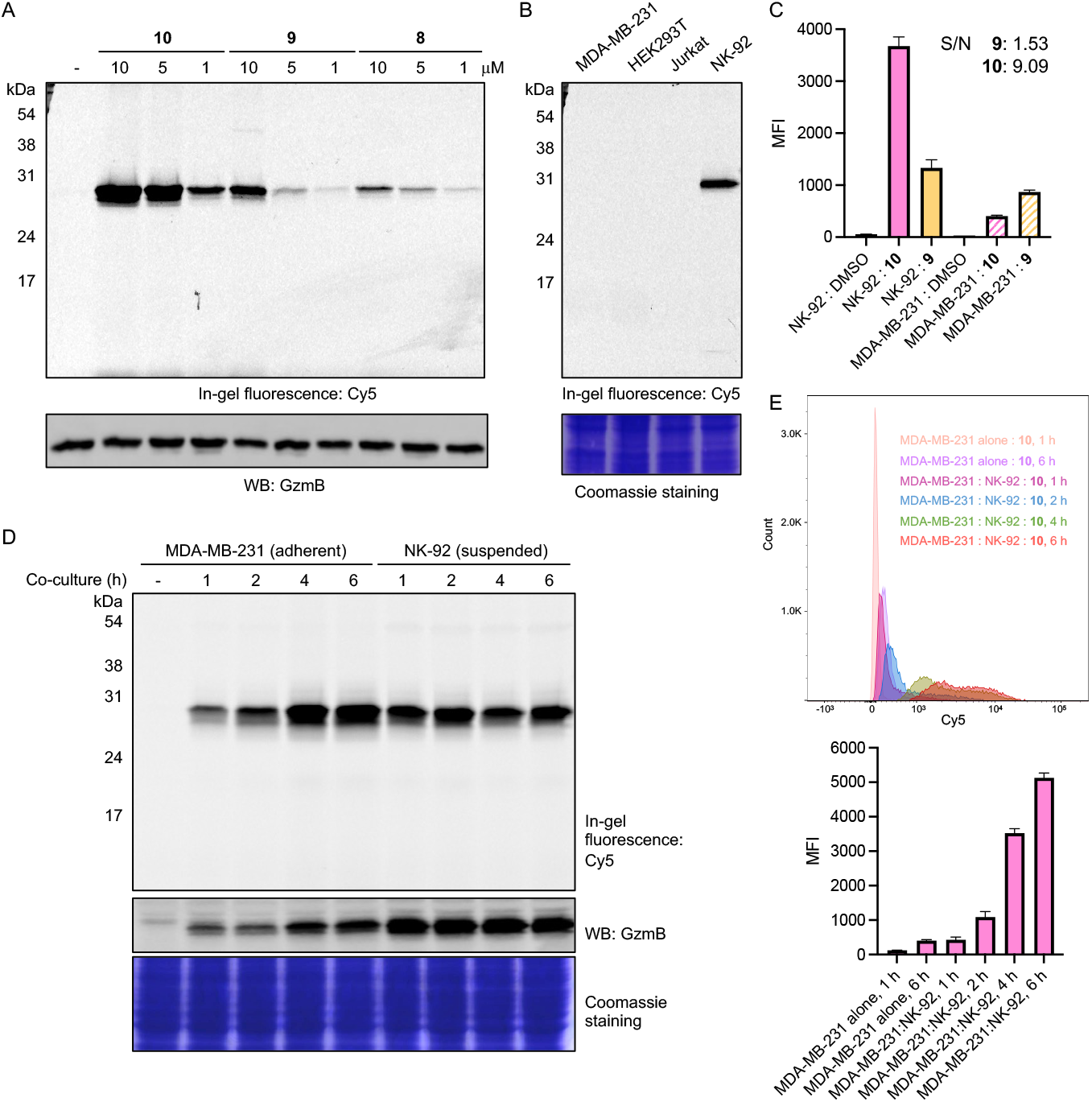
GzmB ABPs selectively label active GzmB in cells. (A) Gel-based analysis of GzmB labeling by ABPs in NK-92 cells. 5×10^5^ NK-92 cells were incubated with each ABP at varying concentrations for 2 h at 37 °C. After cell lysis, protein samples were analyzed by SDS-PAGE and in-gel fluorescence scanning for Cy5 signal. Expression of GzmB was confirmed by western blotting. (B) Gel-based analysis of GzmB labeling by qABP **10** in different cells. 2×10^6^ NK-92, MDA-MB-231, Jurkat, and HEK cells were incubated with compound **10** (5 μM) at 37 °C for 2 h. After cell lysis, protein samples were analyzed by SDS-PAGE and in-gel fluorescence scanning for Cy5 signal. Protein loading was confirmed with Coomassie Brilliant Blue (CBB) staining. (C) Flow cytometry analysis of protein labeling by ABPs **9** and **10** (5 μM) in NK-92 and MDA-MB-231 cells. Data points are displayed as mean ± SD (n= 3). (D) 5×10^5^ MDA-MB-231 cells were co-cultured with 2×10^6^ NK-92 cells in the presence of qABP **10** (5 μM) for the indicated time. NK-92 cells were removed, and MDA-MB-231 cells were washed, harvested, and lysed. Protein samples were analyzed by SDS-PAGE and in-gel fluorescence scanning for Cy5 signal. Expression of GzmB was confirmed by western blotting. Protein loading was confirmed with CBB staining (E) Flow cytometry analysis of protein labeling by qABP **10** (5 μM) in co-cultures of NK-92 and MDA-MB-231 cells. Top: Histogram; Bottom: Mean fluorescence intensity (MFI) in MDA-MB-231 cells. Data points are displayed as mean ± SD (n = 3).

Building on the results from GzmB probes, we also developed probes for GzmA, which has been reported to induce both caspase-independent apoptosis and pyroptosis in cancer cells.^50, 51^ We synthesized phosphonate and phosphinate inhibitors **12** and **13**, incorporating the GzmA substrate sequence Ile-Gly-Asn-Arg (IGNR) (Scheme S4), and compared their ability to inhibit the cleavage of a fluorogenic substrate by GzmA (Figure S6).^52-54^ With comparable inhibitory activity confirmed, we then prepared GzmA fluorescent ABPs **14, 15** (Scheme S5), and qABP **16** in a similar manner to GzmB probes (Scheme S6 and Figure S7A). As shown in Figure S7E, the emission of Cy5 fluorescence was efficiently quenched by QSY21 in **16**. These probes exhibited strong labeling of recombinant human GzmA at sub-micromolar concentrations (Figure S7B), which was blocked upon pretreatment with 5 μM of **12** or **13** (Figure S7C). Probe **16** appeared capable of labeling active hGzmA in a dose- and time-dependent manner, with comparable intensity to **14** and **15**, and no labeling of pro-enzyme was observed. Additionally, qABP **17** was designed based on the recently reported GzmA probe featuring non-natural amino acids L-octahydroindole-2-carboxylic acid (Oic) at P2 and L-1,2,3,4-tetrahydroisoquinoline-3-carboxylic acid (Tic) at P4 (Figure S7D and Scheme S7).^55^ In addition to efficient labeling of hGzmA (Figure S7F), both **16** and **17** were found to label mouse GzmA (Figure S7G). Selective, dose-dependent labeling of GzmA was observed in NK-92 cell lysates at low nanomolar concentrations, without any cross-reactivity found in the lysate of GzmA non-expressing cells (Figure S8A). Incorporating non-natural amino acids in the peptide sequence improved the stability of the probe as we observed only 20% hydrolysis of **17** after 11 h incubation in cell culture media (Figure S8B). However, the ability of qABP **17** to label GzmA in intact NK-92 cells was poor, indicating insufficient cell permeability (Figure S8C). GzmA qABP **16**, on the other hand, was shown to be cell permeable and efficient in labeling GzmA in cells. Therefore, our probe **16** can be utilized to detect GzmA activity and assess its role in the TME.

Taken together, we found that qABPs equipped with the phosphinate ester warhead and the optimal recognition sequence are capable of selectively detecting GzmA/B in complex cell environments.

### In vivo imaging of GzmB activity in a mouse model of cancer immunotherapy

After confirming the selectivity of qABP **10** both in vitro and in cells, we proceeded to evaluate its ability to detect tumor response to immunotherapy through GzmB activity in complex TME. It has been reported that 4T1 tumors are poorly immunogenic, showing minimal immune response and low sensitivity to immunotherapy.^56-58^ Using this more challenging ‘cold’ tumor model,^59^ we asked whether our GzmB-targeting probe could identify the subset of animals that exhibit a higher level of response to immunotherapy at the early stage of treatment. We implanted 4T1 tumors into the mammary fat pad of BALB/c mice and administered three doses of different immunotherapeutic agents (anti-PD1, anti-CTLA4, and BEC) every three days once tumors were established (Figure 4A). The immune checkpoint-blocking antibodies anti-PD1 and anti-CTLA4 reinvigorate immune cell killing of cancerous cells,^60-63^ while BEC, an arginase inhibitor, limits the depletion of L-arginine, a crucial amino acid for immune cell function.^64, 65^ Following the treatment, we excised tumors and performed ex vivo analysis (Figure 4A). In our 4T1 TNBC model, the effects of these immunotherapies on tumor growth were found to be weak (Figure 4B). The average tumor volume was not significantly different between the control (Δ tumor volume (TV) = 405.06 mm^3^) and treatment groups (ΔTV = 394.56 mm^3^). However, among 16 mice in our initial study, mouse #36, which received three doses of anti-CTLA4, exhibited relatively slower tumor growth (ΔTV = 148.38 mm^3^), suggesting a stronger response to therapy. A higher level of GzmB expression was detected in the tumor samples from this mouse (Figures 4C-D), accompanied by a higher number of CD8+ cells (Figure 4F). More importantly, qABP **10** reported a relatively higher level of GzmB activity in the tumor cells from mouse #36 by both gel-based (Figures 4C-D) and flow cytometry (Figure 4E) analysis, suggesting that our probe has the potential to identify individual responding tumors in the early stage of therapy.

**Figure 4.**
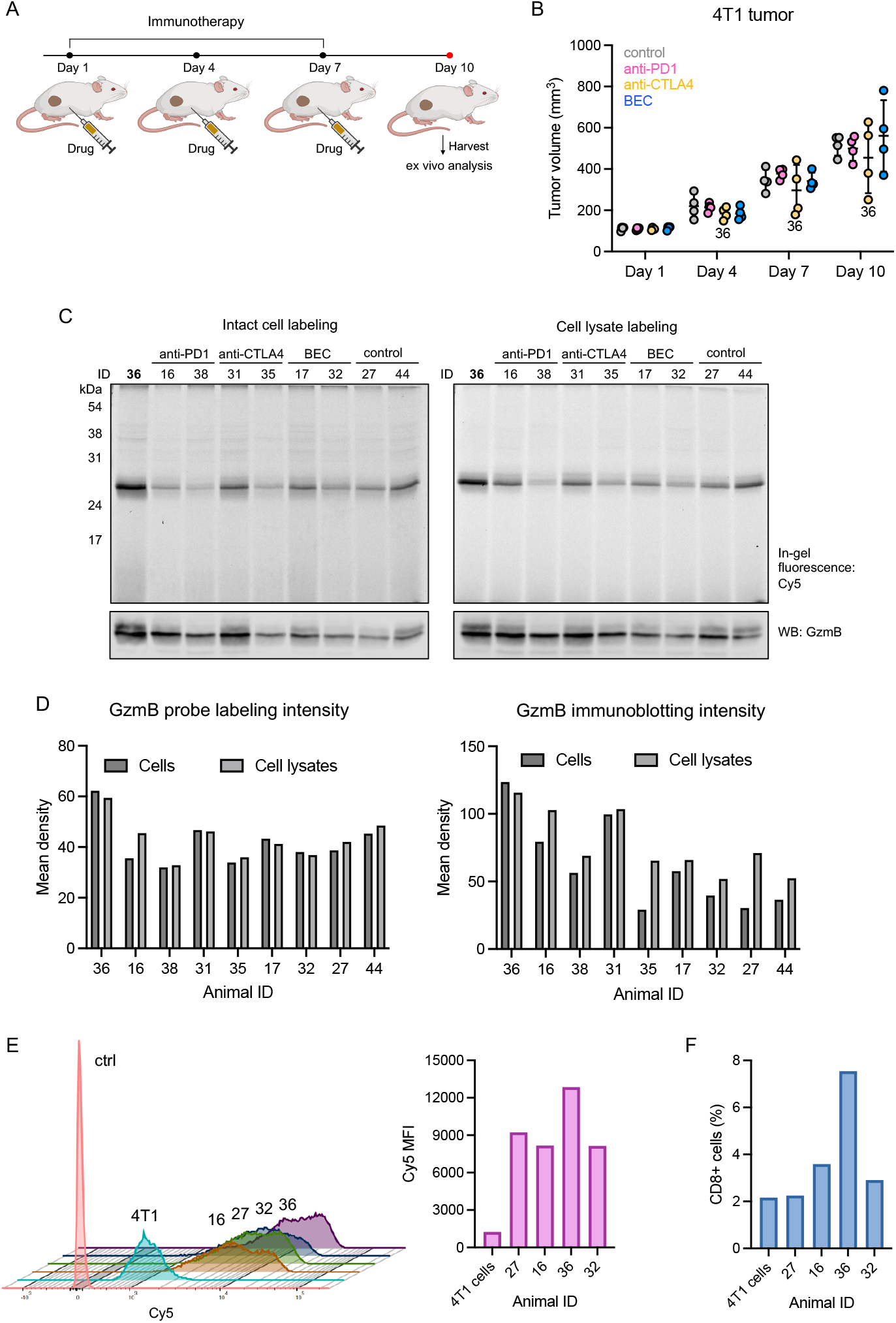
Ex vivo analysis of 4T1 tumor response to immunotherapy using GzmB qABP **10**. (A) Schematic illustration of the timeline of immunotherapy and ex vivo analysis. (B) Measurement of tumor growth. Data points are displayed as mean ± SD (n = 4). (C) In-gel Cy5 fluorescence and GzmB immunoblotting of tumor cell suspensions (left) and tumor lysates (right) treated with **10** (5 μM). (D) Quantification of protein band intensities from Cy5 signals (left) and GzmB immunoblotting (right). (E-F) Flow cytometry analysis of tumor single-cell suspensions and cultured 4T1 cells as control. Briefly, tumors were dissociated into singlecell suspensions and treated with 5 μM of **10** (or DMSO) for 2 h at 37 °C. Cells were fixed in the dark. After permeabilization, fixed cells were treated with FITC-labeled CD8α antibody for 1 h at room temperature and analyzed with flow cytometry. Separately, cultured 4T1 cells were stained under similar conditions. (E) Overlayed histograms (left) and quantification of mean fluorescence intensity (MFI) of Cy5 signals (right). (F) Population of CD8+ cells (%).

Encouraged by these results, we next investigated the in vivo effectiveness of **10** in detecting immune response through real-time fluorescence imaging (Figure 5A). We chose an ICB combination therapy, consisting of anti-PD1 and anti-CTLA4 at 10 mg/kg and 5 mg/kg, respectively (n = 5), as it has been reported to enhance the antitumoral effect compared to monotherapy.^58, 66-68^ Again, 4T1 tumor growth varied greatly among animals, although the average tumor volume was slightly lower in the treated group (ΔTV = 360.58 mm^3^) compared to the untreated group (ΔTV = 417.91 mm^3^) (Figure 5B). Three days after the final treatment, GzmB probe **10** was injected intravenously (i.v.) into the mice (50 nmol/100 μL in 10% DMSO/PBS), and we imaged the mice bearing 4T1 tumors at 2, 4, 8, and 24 h post-injection using an IVIS spectrum imaging system. When imaged prior to injection, a solution of qABP **10** in PBS showed no fluorescence, indicating minimal background signal, whereas ABP **9** exhibited intense fluorescence (Figure S9). In the animals, the NIRF signal was brightest within the tumors at 2 h post-injection and then decreased at 4, 8, and 24 h time points (Figures 5C-D). Notably, the highest in vivo fluorescence signal at 2 h was observed in mouse #57 (4.02×10^10^ p s^−1^ cm^−2^ sr^−1^), one of the mice that showed reduced tumor growth (ΔTV = 262.33 mm^3^). Ex vivo NIRF imaging of organs harvested 24 h post-injection revealed significant signal accumulation in tumors (Figures 5E and S10). When normalized to signals from the spleen (background), the highest relative signal was found in the tumor of mouse #57 (Figure 5F) with a tumor-to-background ratio (TBR) of 15.18. These results suggest that our GzmB-targeting qABP **10** can identify responding tumors with the highest stratifying in vivo signals at 2 h post-probe injection. Of note, the second highest signals were observed in vivo (2 h) in the bladder and ex vivo (24 h) in the kidneys, suggesting that excess probe is cleared through the renal pathway (Figures 5C and S10).

**Figure 5.**
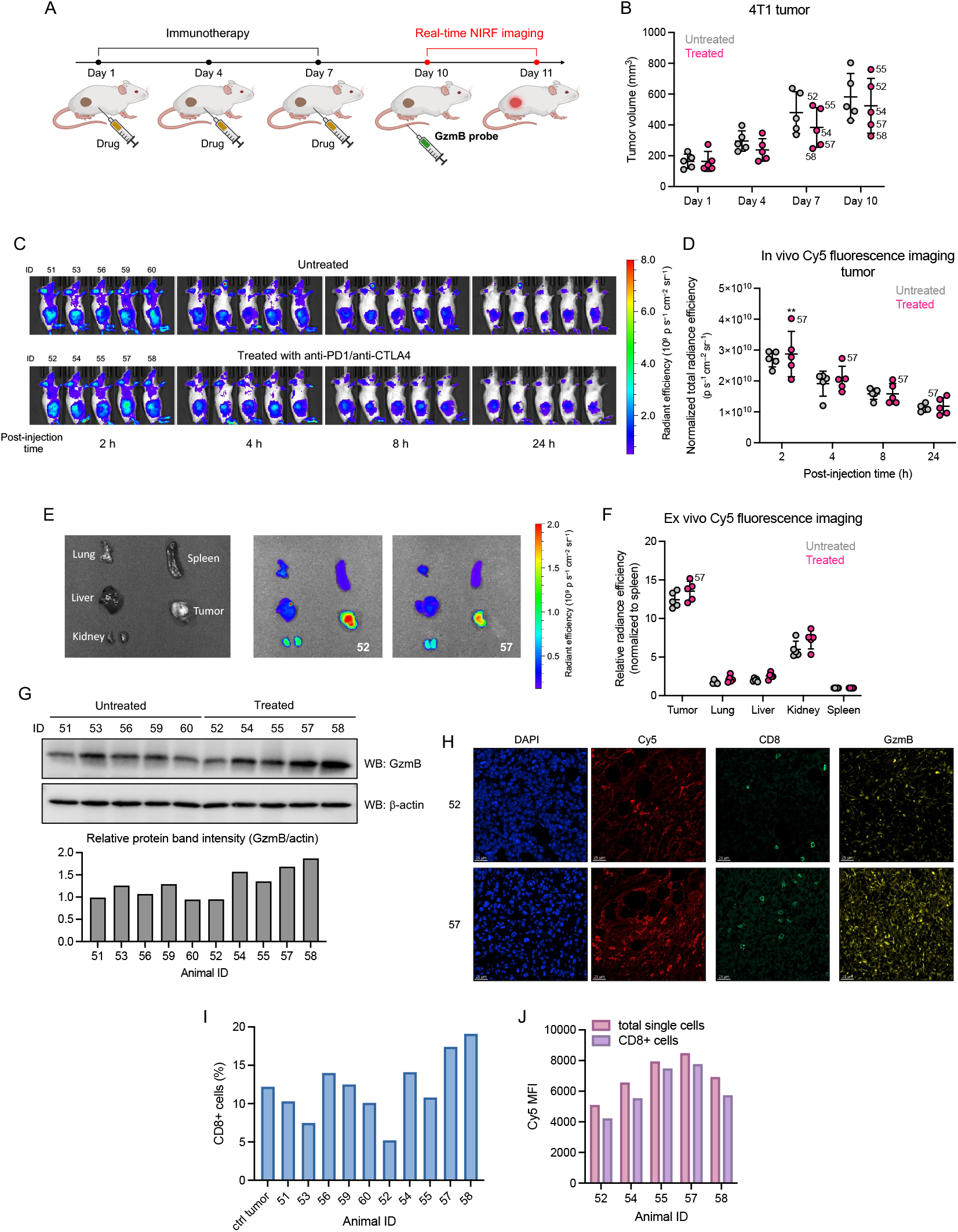
In vivo imaging of GzmB activity in mouse tumor models. (A) Schematic illustration of the timeline of immunotherapy and real-time NIRF-imaging. (B) Measurement of tumor growth comparing immunotherapy (ICB combination)-treated vs. untreated mice. Data points are shown as mean with SD. (C) Whole animal 2D fluorescence images of immunotherapy-treated mice obtained after 2, 4, 8 and 24 h post-injection at Ex/Em: 640/680 nm. (D) Quantification of in vivo NIRF signals in the 2, 4, 8 and 24 h post-injection at Ex/Em: 640/680 nm. (D) Quantification of in vivo NIRF signals in the tumors. Data points are displayed as mean ± SD, and the p-value was evaluated by the Student’s t-test (** p-value ≤ 0.01) (E) Organ layout (left) and ex vivo fluorescence images (right) of organs harvested after 24 h post-injection. (F) Quantification of ex vivo NIRF signals in the organs (normalized to spleen). Data points are displayed as mean ± SD. (G) Immunoblotting analysis of GzmB expression in tumor homogenates. Briefly, pieces of tumors were transferred to a flask containing 1 mL RIPA lysis buffer on ice and constantly grinded for 2 min. Homogenates were collected by centrifugation and 20 μg of proteins were analyzed by SDS-PAGE and immunoblotting. (H) Representative confocal microscopy images of tumor sections stained with DAPI (blue) and CD8α (green)/GzmB (yellow) antibodies and Cy5 (red) detection. (I) Fraction (%) of CD8+ cells in the tumors analyzed by flow cytometry. (J) Mean fluorescence intensity (MFI) of Cy5 signals determined by flow cytometry after dissociated tumors were treated with **10** (5 μM) for 2 h at 37 °C.

To further assess immune cell infiltrates in tumors, we analyzed the harvested tumors by immunoblotting, immunofluorescence (IFC) staining, and flow cytometry. We also attempted to directly detect the labeled GzmB by SDS-PAGE analysis of tissue homogenates collected from animals treated with the probe. Although we observed a faint fluorescently-labeled protein band at the expected molecular weight of GzmB (data not shown), the detection sensitivity of in-gel fluorescence was not sufficient to firmly determine the identity. Immunoblotting of the tumor homogenates, however, revealed overall higher levels of GzmB in the immunotherapy-treated mice (Figure 5G), while no GzmB was detected in the homogenates of other organs (Figure S11). Confocal microscopic image analysis of DAPI-stained tumor sections detected strong Cy5 fluorescence (Ex/Em: 640/680 nm; pinhole radius: 50, Figure S12), indicating the retention of the processed probe signals within tumors (Figures 5H and S13). IFC staining confirmed the presence of CD8α and GzmB in tumor sections. Additionally, flow cytometry analysis of dissociated cell suspensions showed a higher number of CD8+ cells in tumors from therapy-responsive animals (17.4% and 19.1% from #57 and #58, respectively) (Figure 5I). Intense Cy5 fluorescence signals were observed when the tumor cell suspensions were further incubated with qABP **10** (Figure 5J), while background signals in cultured 4T1 cells alone were significantly weaker (Figure S14), suggesting that the high Cy5 signals resulted from GzmB activity in tumors.

The fluorescence intensity and contrast generated within the target tissue depend on the dose and time of imaging after probe injection. Since our qABP **10** exhibited the highest in vivo NIRF signal contrast at 2 h post-injection, we performed additional ex vivo analysis comparing the 2 h vs. 24 h timepoints. 4T1-bearing mice were treated with the ICB combination therapy (n = 5) followed by i.v. injection of qABP **10** (12.5 nmol/100 μL). Two randomly selected mice were sacrificed after 2 h post-injection, while the organs of the remaining three mice were excised after 24 h for further analysis. Among the 5 immunotherapy-treated mice, the brightest in vivo NIRF signal was noted at 2 h post-injection in mouse #61 (7.26×10^9^ p s^−1^ cm^−2^ sr^−1^), one of the animals with the lowest tumor growth (ΔTV = 383.14 mm^3^) (Figures S15A-B). Although ex vivo fluorescence signals were brighter in the tumors of mice sacrificed at 2 h compared to 24 h as expected, significantly high signal accumulation was also observed in the kidneys (Figures S15C-D). Additionally, with a lower concentration of probe injection (1/4), we observed a decreased tumor-to-kidney ratio of ex vivo signals, indicating that an optimal probe concentration is required for achieving high tumor-to-background signals.

Overall, these findings suggest that our GzmB activatable probe is able to quickly assess immune responses in living systems using noninvasive optical imaging.

## Conclusion

Despite decades of research, granzyme biology remains incompletely understood in part due to our limited ability to probe their functions in different cellular and in vivo contexts. GzmB, the only known serine protease that cleaves substrates after aspartic acid (a feature shared with caspases), has a relatively broad, overlapping substrate specificity. Thus, developing chemical tools capable of selectively and efficiently detecting GzmB activity has been challenging. As there has been growing interest in developing GzmB reporters for the real-time assessment of immune-mediated anticancer action, we set out to address the limitations of currently available probes including specificity and kinetic properties.

In this study, we discovered that replacing the critical, negatively charged P1 aspartic acid with a non-natural cysteic acid in the canonical IEPD tetrapeptide sequence enhances binding and selectivity towards GzmB. Leveraging this novel, optimized sequence, we developed ABPs featuring phosphonate or phosphinate warhead and Cy5 fluorophore, which demonstrated the ability to rapidly and selectively label active GzmB in cells. In particular, our quenched ABP **10** ensued high sensitivity with minimal background signal and successfully detected the transfer of GzmB from effector immune cells to target cancer cells. Importantly, when used for in vivo imaging of immune response in tumor-bearing mice undergoing immunotherapy, our probe exhibited significantly high contrast in therapy-responding tumors at early time points. Probe **10** thus represents the first-in-class example of NIR qABPs capable of imaging cancer immunotherapy responses.

How the substitution of carboxylic acid with sulfonic acid in our probe manifests increases in binding affinity and selectivity for GzmB is still unclear. However, it suggests that the S1 pocket of the GzmB active site might be sufficiently different from that of Caps-8. Further structural investigation and analysis of molecular dynamics simulation could help understand molecular interactions of the IEPCya in the active site of GzmB or Casp-8. Nevertheless, these improvements have allowed for the detection of active GzmB even at a substoichiometric concentration of 2.5 nM. Additionally, sulfonic acids are more acidic than carboxylic acids (2– 3 pKa units more acidic) and exhibit strong hydrophilicity. The incorporation of the sulfonic acid group into the peptide sequence in combination with Cy5 and QSY21 provided high solubility while maintaining cell permeability.

To generate a qABP with efficient quenching but fast activation by GzmB, we utilized a recently reported phosphinate ester scaffold that carries only a single leaving group. Unlike phosphonate esters, phosphinate esters cannot undergo a second hydrolysis (aging) after their initial reaction with the active site serine. These features collectively also create opportunities to engage in additional interactions with the primed site of the target protease by exploring various substituents on the phenyl ring. To further illustrate the utility of the phosphinate-qABP strategy, we also generated probes for GzmA, expanding our arsenal to capture active GzmA in cells and in vivo, thereby better defining granzyme biology.

ABPs have been extensively utilized in various in vitro characterization, including gel-based and chemoproteomic platforms.^69-72^ However, their use in evaluating disease progression or treatment responses in vivo has seen limited exploration in cancer immunotherapy. This is partly due to the fact that ABPs inhibit the activity of target protein through covalent binding. Thus, maintaining a balanced control of probe concentration, post-injection timeframe, and the intensity of in vivo signals is critical to maximizing their effectiveness as imaging agents while minimizing disruption to endogenous biological environments. In contrast to the reported GzmB substrate-based fluorogenic probes (10 μmol/kg), which rely on enzymatic turnover for signal amplification and generate stratifying signals around 8-24 h post-injection,^30^ our qABP (2.5 μmol/kg) was found to rapidly react with GzmB, producing high signal contrast in therapy-responding tumors as early as 2 h post-injection, marking our first imaging timepoint. We also observed that the fluorescence signal in tumors remained bright even at 24 h post-injection of our qABP with a high tumor-to-background ratio. Our future study will involve a more comprehensive in vivo kinetic analysis of probe activation and dose optimization to determine the ideal end-point measurements with an optimal dose.

In our current study, we have chosen a poorly immunogenic 4T1 tumor model to detect individual responses. As evidenced by the tumor volume measured post-therapy, most animals in a small cohort (n = 5), however, did not respond positively to the short treatments. Thus, we plan to expand the size and time course of in vivo studies to confirm the early diagnostic power of our qABP **10** for accurately evaluating antitumor immune responses. Larger studies involving more animals would allow to sample a higher number of responders to assess the correlation with NIRF signals with statistical accuracy. Additionally, as studies have shown that different murine syngeneic tumor models exhibit variable responsiveness to immunotherapies,^73^ our GzmB qABP **10** might offer a way to measure varying degrees of immune responses and characterize tumor models for preclinical immunotherapy research. Taken together, we believe this study provides a new option for studies applying optical imaging to assess antitumor immunity and monitor the efficacy of novel immunotherapies in complex biological settings.

## ASSOCIATED CONTENT

### Supporting Information

The Supporting Information is available free of charge on the ACS Publications website.

Additional figures, detailed procedures for all experiments, specifics for reagents, instruments used, and synthetic details (PDF)

## AUTHOR INFORMATION

### Author Contributions

The manuscript was written with contributions from all authors. All authors have approved the final version of the manuscript.

### Notes

The authors declare no competing financial interest.

## ACKNOWLEDGMENT

The authors gratefully thank the staff members at the CCR-Frederick Biophysics Resource (BR), Dr. Jeff Carrell (CCR-Frederick Flow Cytometry Core Laboratory), and Dr. Stephen Lockett and Dr. Valentin Magidson (NCI-Optical Microscopy laboratory) for their assistance, technical consultation, and instrument maintenance. We acknowledge Chelsea Sanders and other staff members at the Laboratory Animal Sciences Program for assistance with the in vivo study. The authors thank Dr. Tania Lopez Silva (CBL) for help with IFC and Dr. Martin Schnermann (CBL) for critical reading of the manuscript. This work was supported by the Intramural Research Program of the National Institutes of Health, National Cancer Institute, Center for Cancer Research (ZIABC011962) and has been funded in part with Federal funds from the NCI, NIH, under Contract No. HHSN261201500003I. The content of this publication does not necessarily reflect the views or policies of the Department of Health and Human Services, nor does mention of trade names, commercial products, or organizations imply endorsement by the U.S. Government.

## ABBREVIATIONS

ICB: immune checkpoint blockad
CTL: cytotoxic T lymphocyte
NK: natural killer
TME: tumor microenvironment
TIME: tumor immune microenvironment
ABP: activity-based probe
NIR: near-infrared
NIRF: near-infrared fluorescence
SBR: signal-to-background ratio
qABP: quenched ABP
Gzm: granzyme
Casp: caspases
FRET: Förster resonance energy transfer
ABPP: activity-based protein profiling
TNBC: triple-negative breast cancer
S/N: signal-to-noise
TV: tumor volume
TBR: tumor-to-background ratio
IFC: immunofluorescence.

## Notes

### Competing Interest Statement

The authors have declared no competing interest.

