## Supplementary information for "Granzyme-targeting quenched activity-based probes for assessing tumor response to immunotherapy"

#### Supplementary Figures

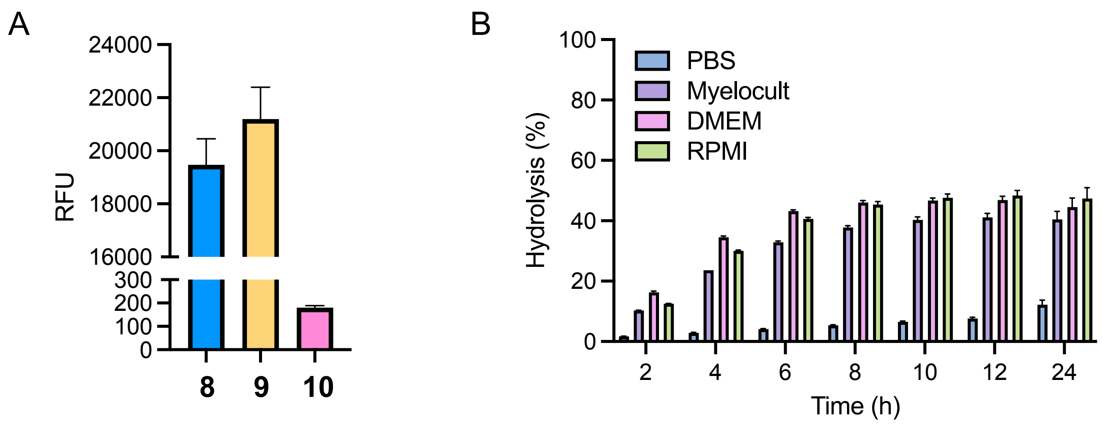


Supplementary Figure 1. Quenching effect and stability of qABP **10**. (A) The fluorescence for each probe (10 μM) was measured at Ex/Em: 640/670 nm. (B) The stability of qABP **10** in PBS and different cell culture media (supplemented with 10% FBS) was measured by monitoring the production of Cy5 fluorescence signal over time. Data points are displayed as mean ± SD (n =3).

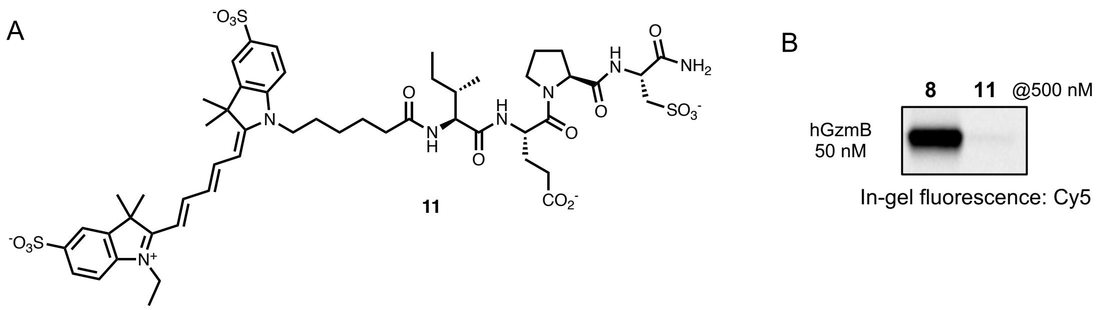


Supplementary Figure 2. (A) Chemical structure of compound **11**. (B) hGzmB labeling assay with **11**. Recombinant human active GzmB (50 nM) was incubated with **8** or **11** (500 nM) for 1 h. Protein samples were analyzed by SDS-PAGE and in-gel fluorescence scanning for Cy5 signal.

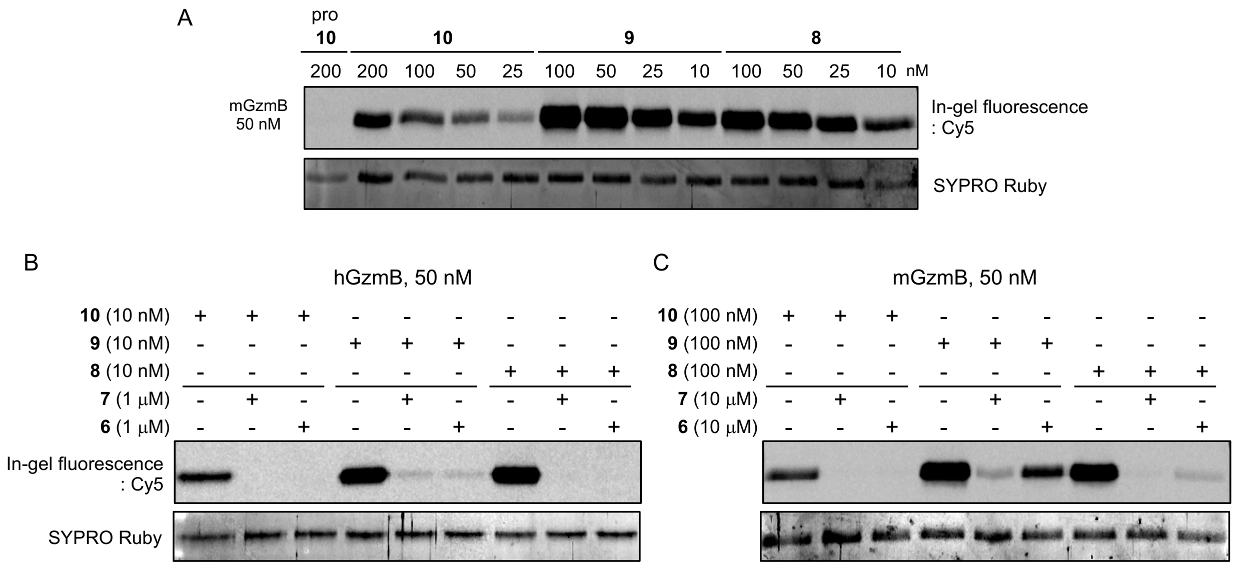


Supplementary Figure 3. (A) mGzmB labeling assay with ABPs **8**-**10**. Recombinant mouse GzmB (50 nM) was incubated with varying concentrations of each ABP for 30 min. (B-C) Competitive labeling assay. Recombinant human (B) or mouse (C) active GzmB (50 nM) was preincubated with covalent inhibitors **6** or **7** (1 μM for hGzmB or 10 μM for mGzmB) for 1 h followed by labeling with ABPs **8**-**10** (10 nM for hGzmB or 100 nM for mGzmB) for 30 min. Protein samples were analyzed by SDS-PAGE and in-gel fluorescence scanning for Cy5 signal and SYPRO Ruby staining.

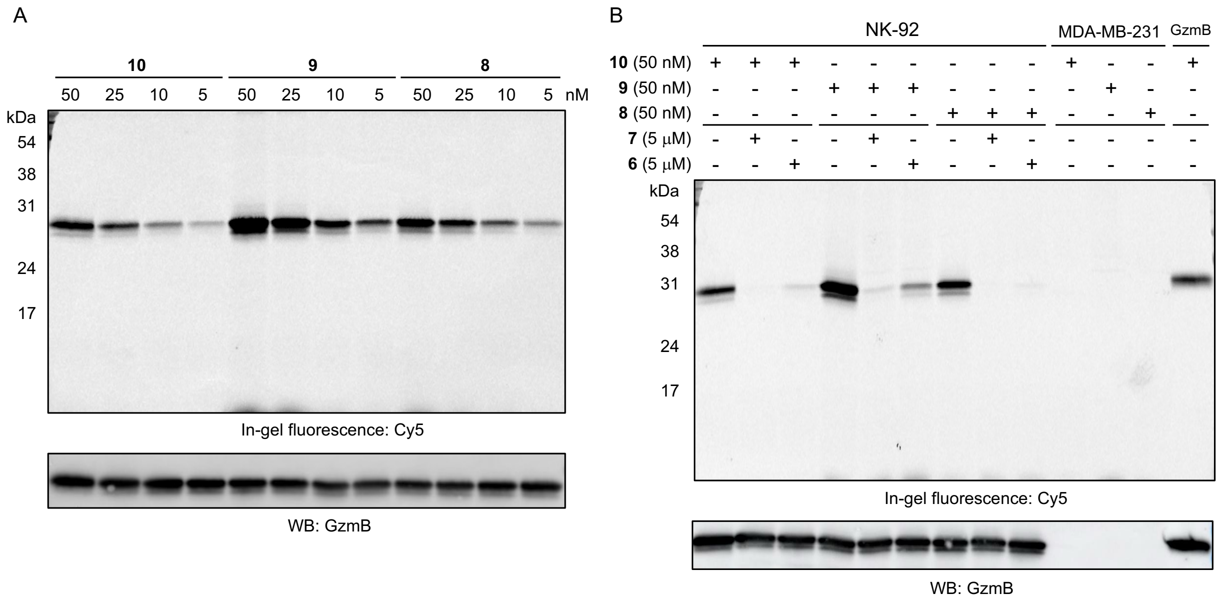


Supplementary Figure 4. In vitro cell lysate labeling. (A) 20 μg of protein samples prepared from NK-92 cell lysates were incubated with each ABP at varying concentrations for 30 min at 37 ˚C. (B) 20 μg of protein samples prepared from the lysis of NK-92 (or MDA-MB-231) cells were treated with 5 μM of compounds **6** and **7** (or DMSO) at 37 °C for 1 h while shaking at 400 rpm. Then, 50 nM of compounds **8-10** were added and samples were incubated at 37 °C for 30 min while shaking at 400 rpm. Protein samples were analyzed by SDS-PAGE and in-gel fluorescence scanning for Cy5 signal. Expression of GzmB was confirmed by western blotting.

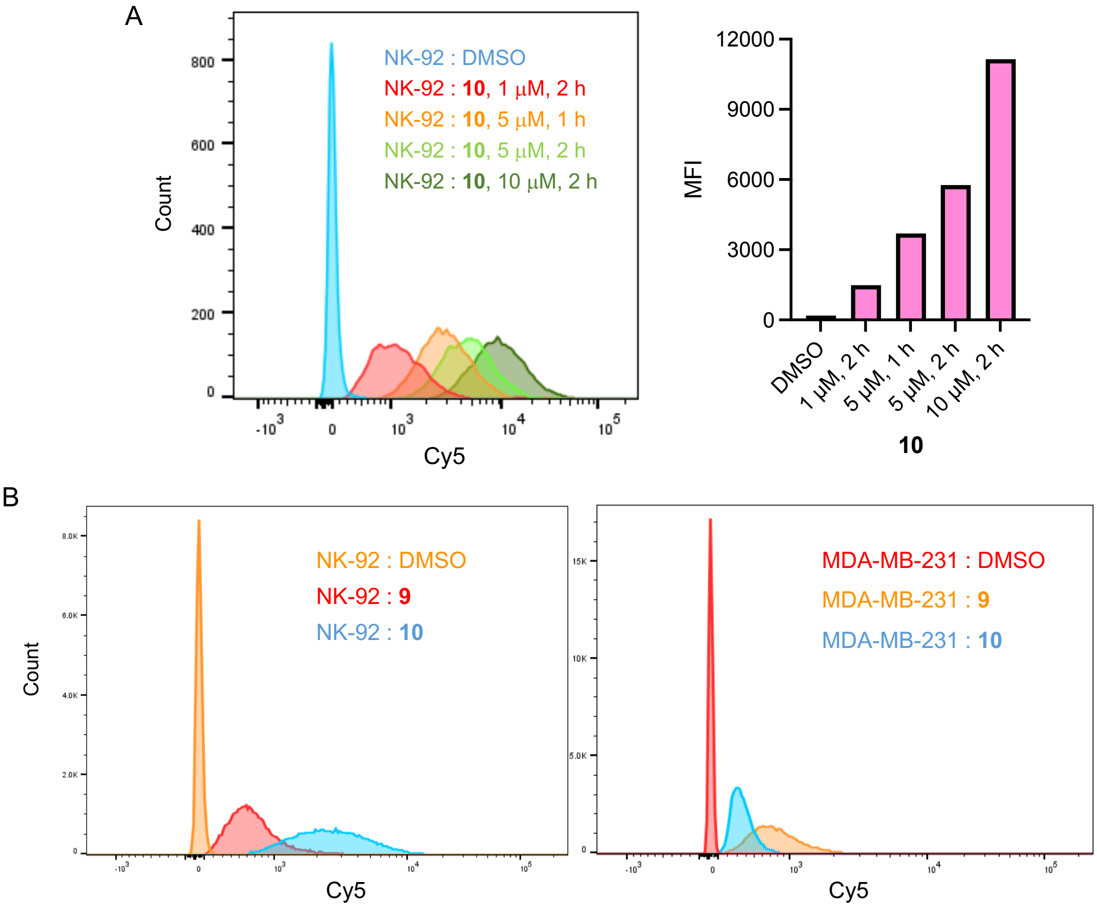


Supplementary Figure 5. Flow cytometry analysis of intact cell labeling with GzmB ABPs. (A) Left: Histogram of flow cytometry data for NK-92 cells incubated with qABP **10**. Right: Quantification of mean Cy5 fluorescence intensity. (B) Histogram of flow cytometry data for NK-92 (left) and MDA-MB-231 (right) cells incubated with ABPs **9** or **10** (5 μM, 37 °C, 2 h).

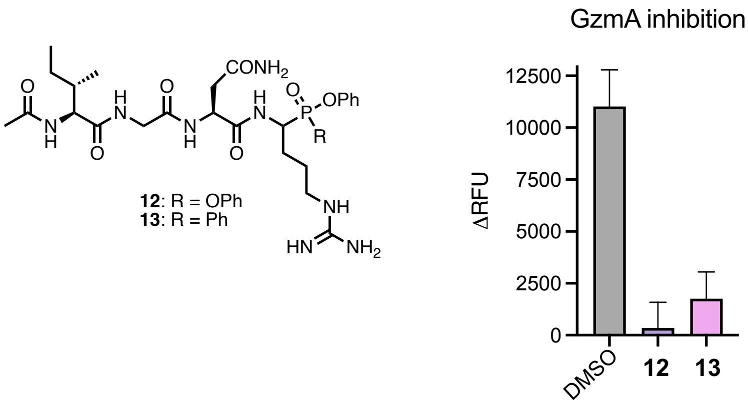


Supplementary Figure 6. Inhibition of GzmA activity. Left: Structures of inhibitors **12** and **13**. Right: Residual GzmA activity in the presence and absence of inhibitors. Briefly, hGzmA (50 nM) was incubated with inhibitor **12** or **13** (200 nM) and GzmA fluorogenic substrate (Ac-Oic-Gly-Pro-Arg-PABA-MU, 100 μM) for 15 h at 37 °C. Recorded fluorescence at Ex/Em: 380/460 nm for residual GzmA proteolytic activity. Data points are displayed as mean ± SD (n =3).

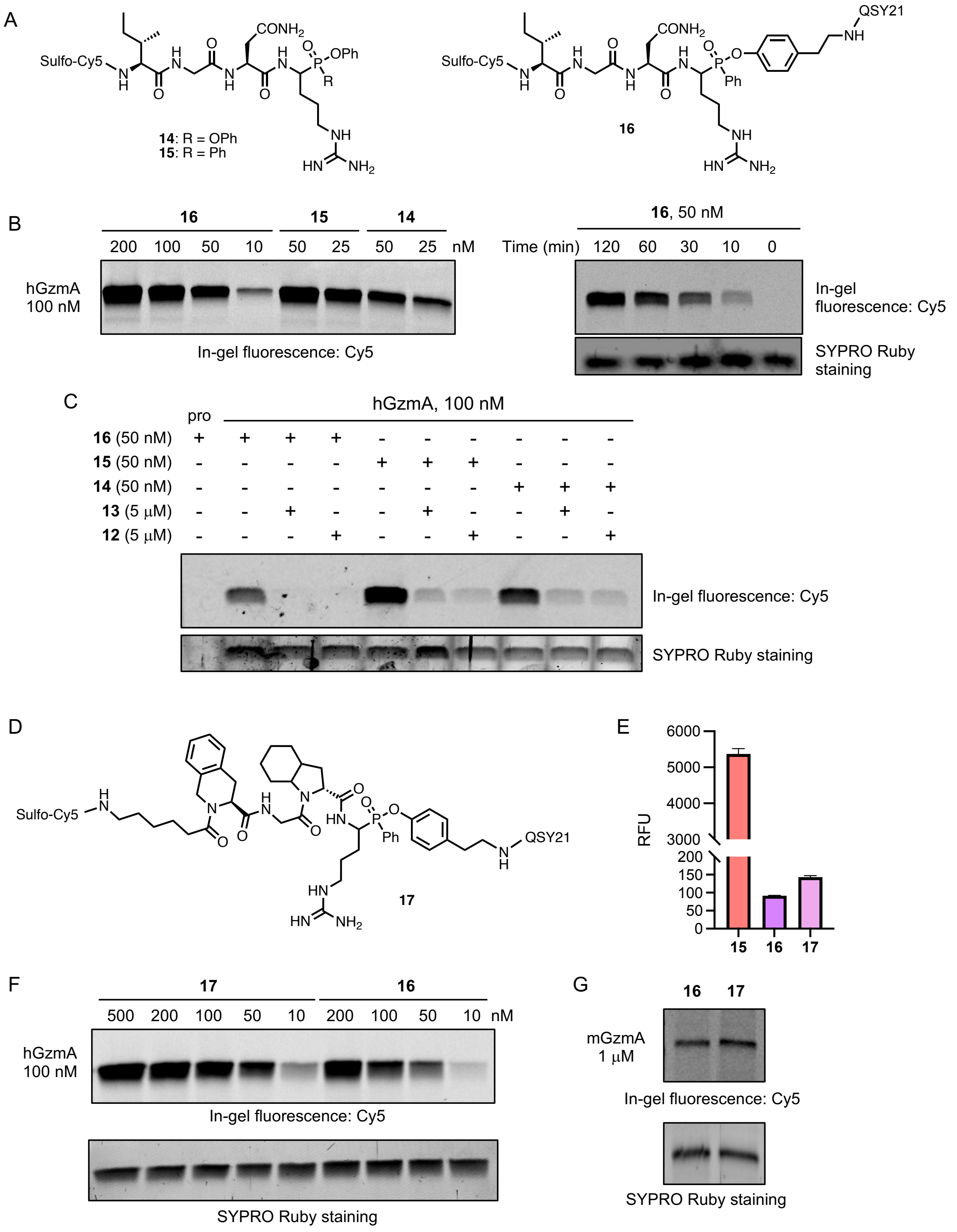


Supplementary Figure 7. GzmA qABPs **16** and **17**. (A) Structures of **14-16**. (B) Dose and time-dependent labeling of hGzmA with **14-16** (GzmA: 100 nM, 2 h, 37 °C). (C) Competitive labeling of hGzmA with **14-16** (pre-treatment of inhibitor for 2 h followed by ABP incubation for 2 h at 37 °C). (D) Structure of **17**. (E) Quenching effect of **16** and **17**. Fluorescence was recorded at Ex/Em: 640/670 nm for each probe (10 µM). Data points are displayed as mean ± SD (n =3). (F) Dose-dependent labeling of hGzmA with **16** and **17** (2 h, 37 °C). (G) Labeling of mGzmA with **16** and **17** (1 µM, 24 h, 37 °C, Buffer: 50 mM Tris, pH 8).

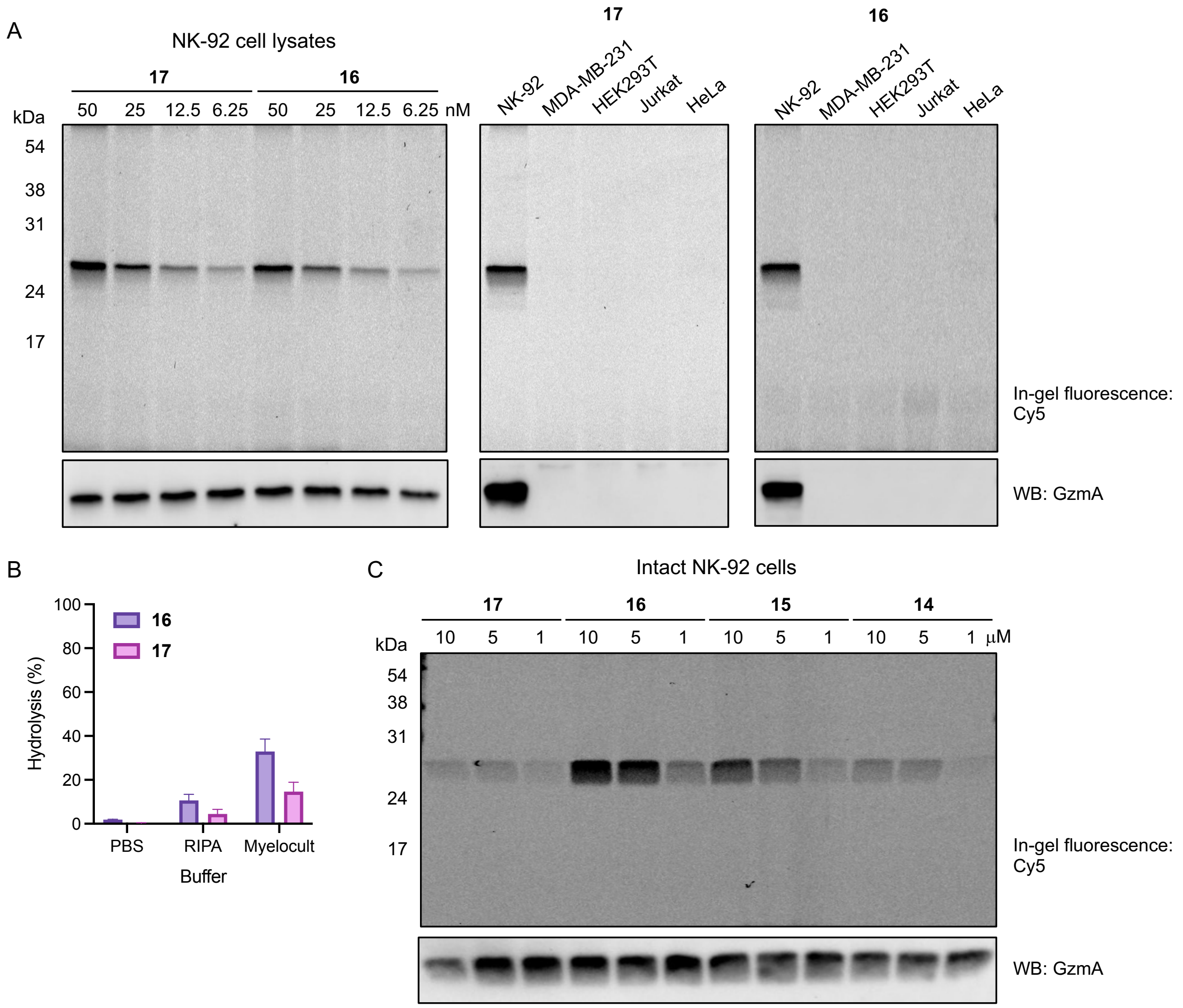


Supplementary Figure 8. Activity-based GzmA profiling. (A) Left: Dose-dependent labeling of GzmA with **16** and **17** in NK-92 cell lysates. Middle, Right: GzmA selective labeling in different cell lysates with **17** (Middle) and **16** (Right) (50 nM, 1 h, 37 °C). Briefly, cell lysates were prepared with RIPA lysis buffer, and 20 μg of proteins were incubated with 50 nM of **16** or **17** for 1 h at 37 °C and analyzed by SDS-PAGE. (B) Relative hydrolysis (%) of **16** and **17** (1 μM) when incubated at 37 °C for 11 h in different media. Data points are displayed as mean ± SD (n =3). (C) Dose-dependent in situ labeling of GzmA in NK-92 cells with **14**-**17** (2 h, 37 °C). Briefly, $1x10^6$ NK-92 cells were incubated with different concentrations of ABPs for 2 h at 37 °C. After washing with PBS (x2), cells were lysed with RIPA lysis buffer at 0 °C for 30 min and lysates were collected by centrifugation. After denaturing with 4X SDS-reducing buffer, 20 μg of proteins were analyzed by SDS-PAGE and in-gel fluorescence scanning for Cy5 signal.

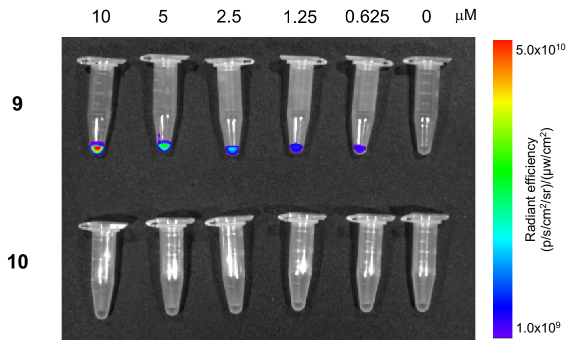


Supplementary Figure 9. Measurement of fluorescence of probes with the IVIS Spectrum imaging system. 2D fluorescence images of microcentrifuge tubes containing varying concentrations of **9** and **10** were obtained at Ex/Em: 640/680 nm.

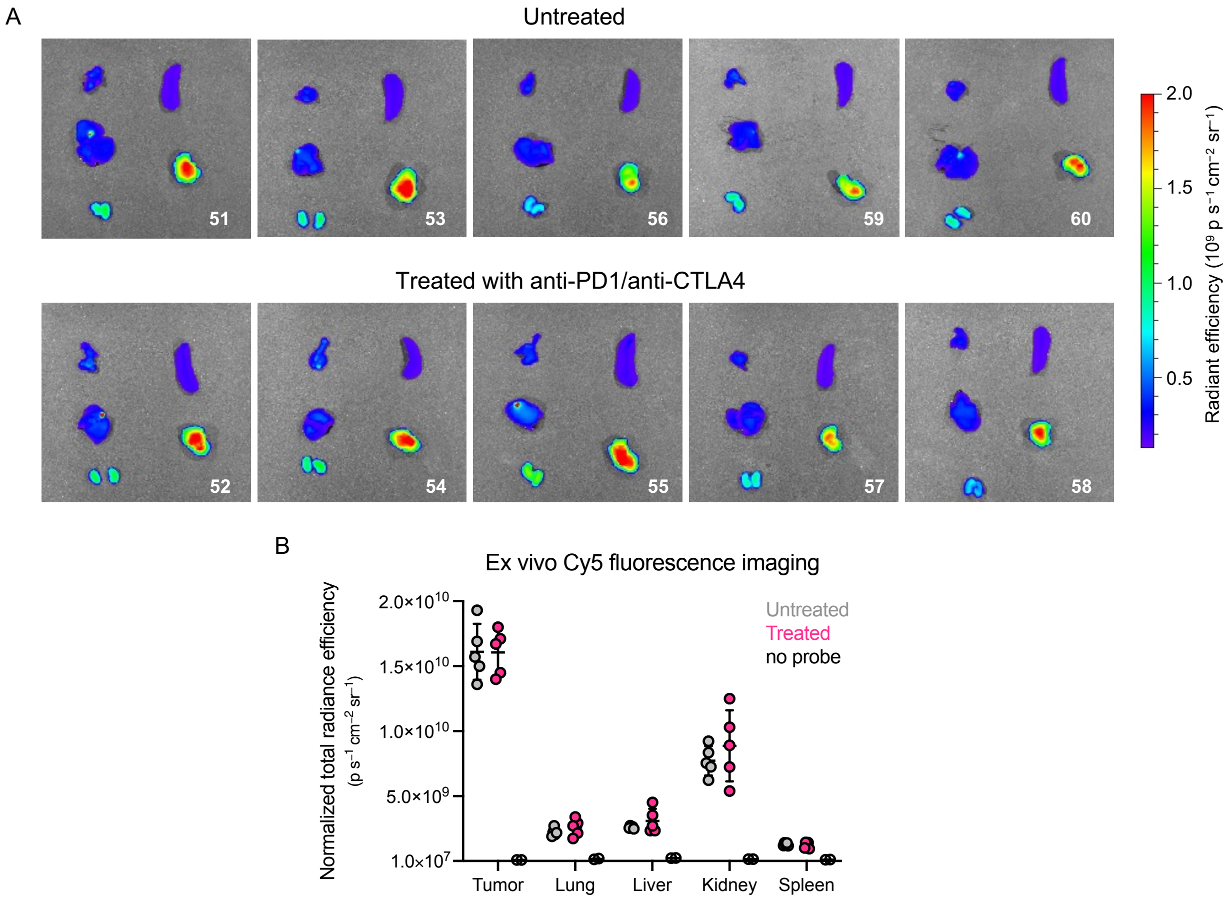


Supplementary Figure 10. Ex vivo fluorescence images of organs harvested 24 h after probe injection. (A) 2D fluorescence images of organs (lung, liver, kidneys, spleen, tumor) obtained with IVIS Spectrum at Ex/Em: 640/680 nm. (B) Quantification of Cy5 fluorescence signals in different organs. Data points are displayed as mean ± SD.

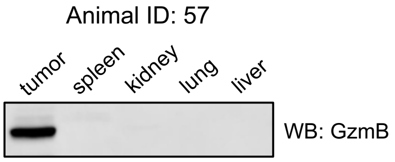


Supplementary Figure 11. Immunoblotting for GzmB in homogenates of different organs harvested 24 h after probe injection. Small pieces of organs were transferred to a flask containing 1 mL RIPA lysis buffer on ice. The mixture was constantly ground for 2 min (roughly 50 strokes), homogenates were transferred to microcentrifuge tubes, and the supernatant was collected by centrifugation (15000g, 4 °C, 30 min). Protein concentration was determined with a pierce BCA protein assay and 30 μg of proteins were analyzed by SDS-PAGE. Expression of GzmB was confirmed by western blotting.

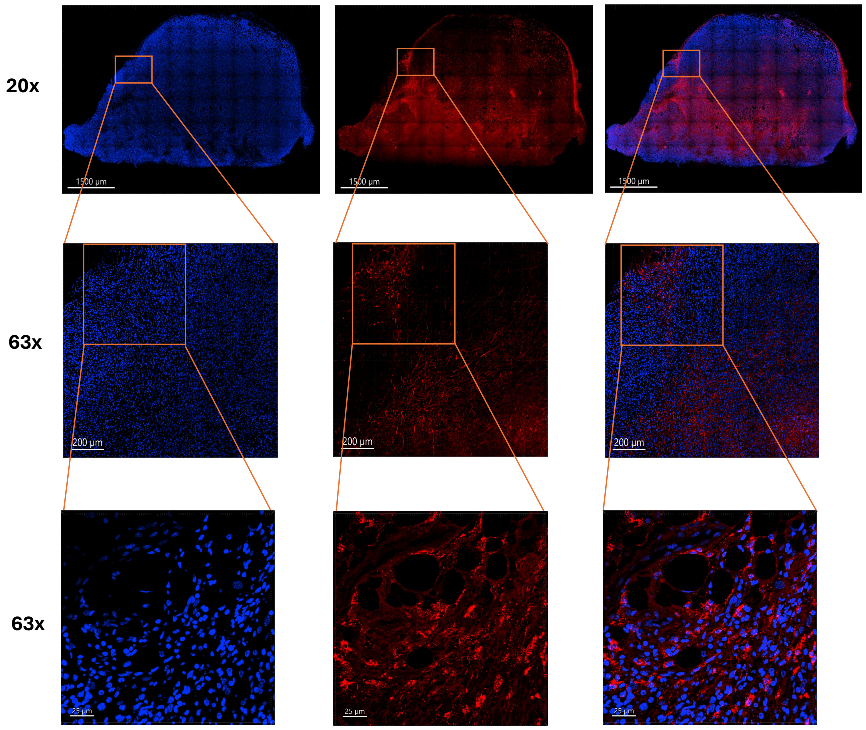


Supplementary Figure 12. Schematic overview of confocal microscopic image analysis of tumor sections. Briefly, paraffin-embedded tissues were processed according to the standard protocol outlined in the experimental section. Images of the entire tissues were first acquired under a 20x dry lens. Regions of interest were then imaged under a magnified 63x (oil immersion) lens to achieve higher resolution. Representative fields from the stitched file were selected as final images.

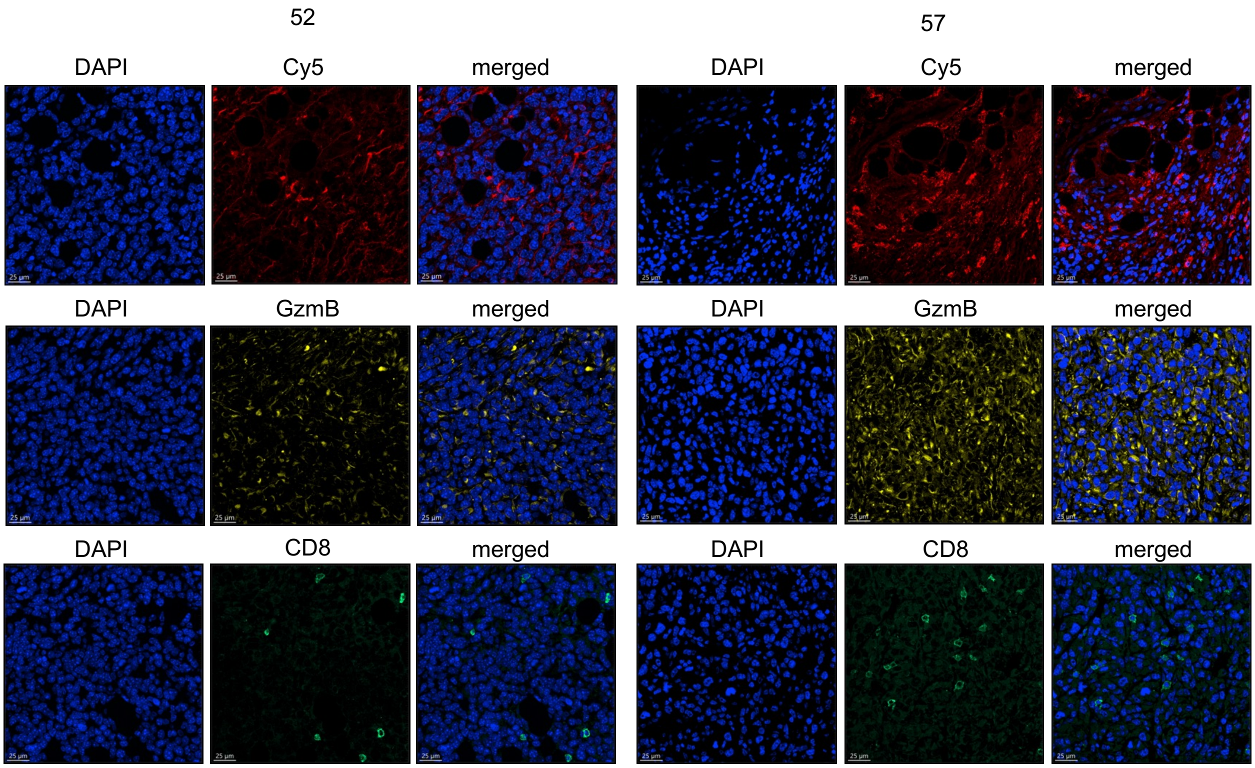


Supplementary Figure 13. Representative confocal images of immunofluorescence (IFC) stained tumor sections for animals #52 and #57. Paraffin-embedded tumor sections were processed according to the standard protocol outlined in the experimental section. Top Lane: Cy5 (red) detected and DAPI (blue) stained; Middle Lane: DAPI and GzmB antibody (yellow) stained; Bottom Lane: DAPI and CD8 antibody (green) stained.

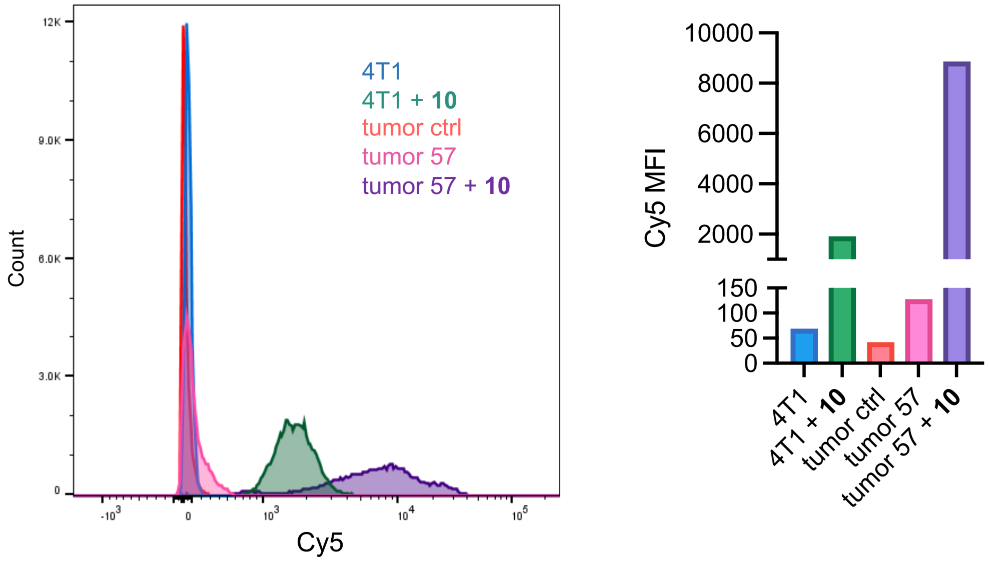


Supplementary Figure 14. Flow cytometry analysis of intact cell labeling for tumor cell suspension and 4T1 cells. Left: Histogram of flow cytometry data for 4T1 cells and tumor cells of mouse #57 treated with **10** (5 μM, 2 h, 37 °C). Right: Quantification of Cy5 signals (mean fluorescence intensity) from flow cytometry data of cells labeled with **10** (5 μM, 2 h, 37 °C).

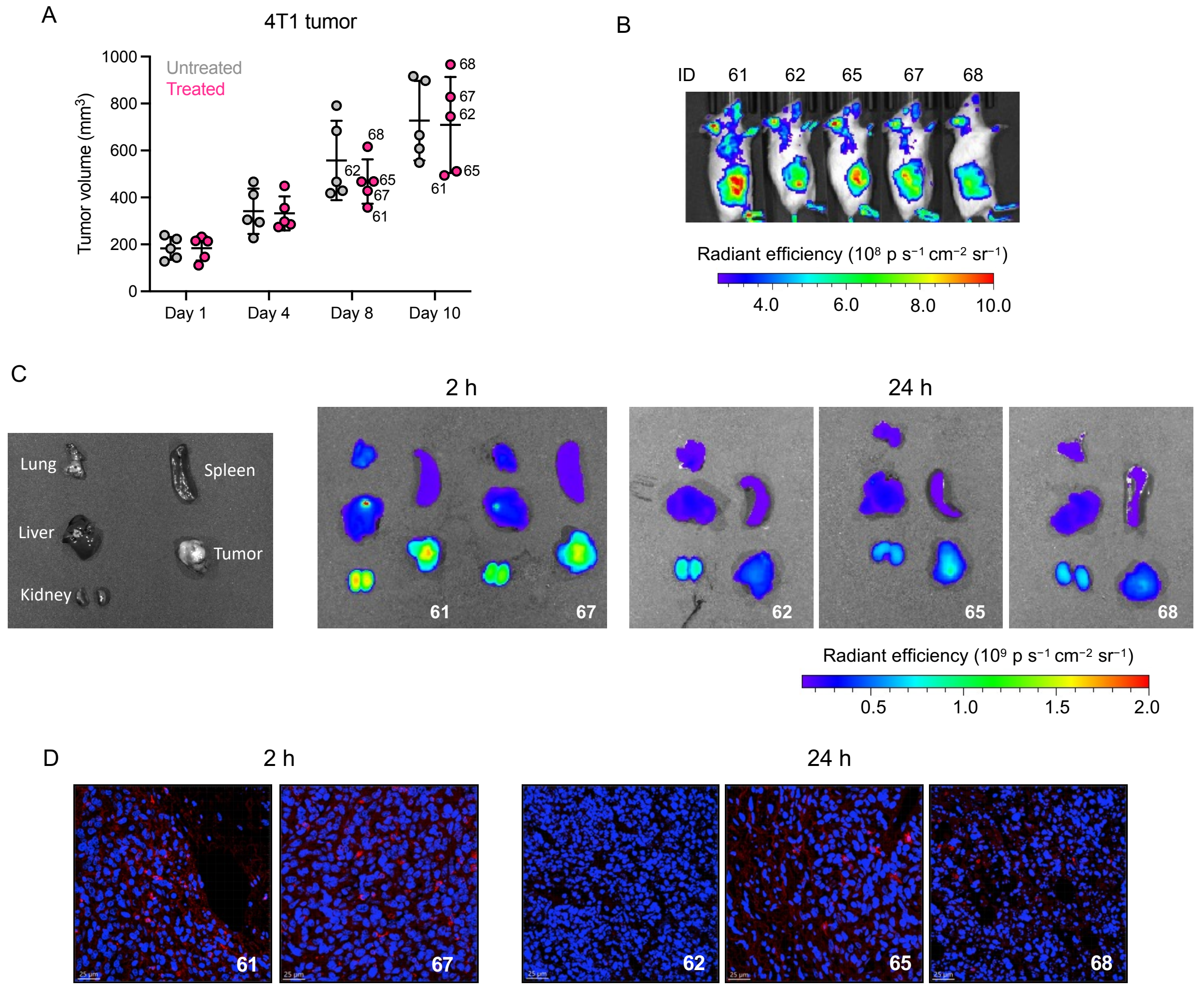


Supplementary Figure 15. Ex vivo analyses at 2 h vs 24 h post probe injection. (A) Measurement of tumor volume over the course of immunotherapy. Data points are displayed as mean ± SD. (B) Whole animal 2D fluorescence images of immunotherapy-treated mice obtained with IVIS Spectrum at Ex/Em: 640/680 nm after 2 h post-injection. (C) Organ layout for ex vivo imaging (left), ex vivo fluorescence images of organs harvested after 2 h (middle) and 24 h (right) of probe injection. (D) Confocal images of Cy5 (red) -detected and DAPI (blue)-stained tumor sections.

### Methods

#### A. Enzyme kinetic assays

All kinetic experiments were performed on a spectrofluorometer (Biotek cytation5 or Synergy|Mx microplate reader) and analyzed using Microsoft Excel and GraphPad Prism. Unless otherwise stated, granzyme B assay buffer consisted of 50 mM HEPES, 50 mM NaCl, 0.01% Tween 20, pH 7.4, whereas the assay buffer for caspases contained 25 mM HEPES, 10 mM DTT, 0.1% CHAPS, pH 7.4. Granzyme A assay buffer consisted of 50 mM Tris, 100 mM NaCl, 5 mM CaCl2, 0.01% (v/v) Tween-20, pH 7.5. Assay buffers were prepared at room temperature while the kinetic experiments were performed at 37 °C. Data are presented as the mean with standard deviation (at least two independent experiments). For kinetic experiments, pro-GzmB (0.22 mg/mL) was activated with mouse cathepsin C (22 μg/mL in 50 mM MES, 50 mM NaCl, pH 5.5) and diluted in assay buffer. Alternatively, active GzmB at 100 U/μL was purchased from Enzo Life Sciences (#BML-SE238-5000) and diluted in assay buffer. Conditions for individual experiments are described below:

**Figure 1B-C:** 10 nM of active GzmB or 5 nM of Casp-8 were treated with different substrates (compounds **1-4**) and recorded fluorescence (Ex/Em: 380/460 nm) for 1 h at 37 °C. To determine Michaelis-Menten constants, cleavage assays with 10 nM of GzmB or 5 nM of Casp-8 were performed with different concentrations of substrates **1** and **2**.

**Figure 1E:** Inhibition assays were performed with inhibitors **5-7** (in 50 mM Tris, 100 mM NaCl, pH 7.4) to determine $IC_{50}$ values. In 384-well black microplates, 100 μM of Ac-IEPD-AMC (Cayman Chemicals, #27140) and different concentrations of inhibitors were plated, then treated with 2 U/μL of GzmB and recorded fluorescence (Ex/Em: 380/460 nm) for 1 h.

#### B. Cell culture and preparation of cell lysates

All cell lines were cultured at 37 °C under 5% $CO_2$ in a humidified atmosphere. Jurkat and 4T1 cells were cultured in RPMI supplemented with 10% fetal bovine serum (FBS), 1% penicillin/streptomycin (PS), and 2 mM L-glutamine, while HEK293T and MDA-MB-231 cells were cultured in DMEM supplemented likewise as described above. NK-92 cells were cultured in Myelocult™ H5100 media supplemented with Gibco™ Horse Serum (63 mL/500 mL media; New Zealand Origin), penicillin/streptomycin (1% of culture media), and IL-2 (1 mL/500 mL media). To prepare the lysates, cells were harvested, pelleted (125g, 6 min), washed with ice-cold PBS, resuspended in ice-cold RIPA (40 μL per $1\times10^6$ cells) on ice for 30 min and pelleted by ultracentrifugation (15000g, 30 min, 4 °C). The supernatant was collected, and protein concentration was measured using a Pierce BCA protein assay. Protein concentrations were adjusted to 1 mg/mL and stored at -80 °C (or used for further analysis).

#### C. In-gel fluorescence imaging

For recombinant enzyme labeling, pro-GzmB (0.22 mg/mL) was activated with mouse cathepsin C (22 μg/mL in 50 mM MES, 50 mM NaCl, pH 5.5) for 4 h at 37 °C, then diluted to 400 nM in assay buffer that contained 50 mM HEPES, 50 mM NaCl, 0.01% Tween 20, pH 7.4 and aliquots were stored at -80 °C. Working concentrations were prepared by further diluting in the assay buffer before use. Mouse pro-GzmB (0.474 mg/mL) was also activated with mCatC (52.14 μg/mL in 50 mM MES, 50 mM NaCl, pH 5.5), diluted in assay buffer and stored as 500 nM aliquots at -80 °C, which were further diluted in assay buffer to make final working concentrations. In-gel fluorescence images were acquired on a Typhoon FLA9500 instrument while western blot and Coomassie staining images were acquired on an AMHERST ImageQuant 800 instrument.

**Conditions for probe/inhibitor treatment for in-gel fluorescence analysis:** Inhibitors and probes were incubated with active (or pro-) GzmB followed by treatment with 4X SDS reducing buffer. Samples were then heated to 100 °C for 5 min, brought to room temperature, and 20 μL of each sample was loaded to 15% polyacrylamide gel, which was then subjected to electrophoresis followed by the acquisition of in-gel fluorescence images. GzmB labeling was confirmed by either western blotting or SYPRO Ruby staining, while protein loading was confirmed with Coomassie Brilliant Blue (CBB) staining of the gel. Incubation conditions for each figure/experiment are described below:

**Figure 2E:** In microcentrifuge tubes, 10 nM of compounds **8-10** were treated with 50 nM active GzmB (or pro-GzmB) for 1 h at 37 °C with a total reaction volume of 20 μL. For dose-dependent labeling, different concentrations of compounds **8-10** were treated with 50 nM of active GzmB (total reaction volume of 20 μL) and incubated at 37 °C for 1 h while shaking at 400 rpm. The same protocol was repeated with mouse GzmB wherein different concentrations of compounds **8-10** were treated with 50 nM of active mouse GzmB (or pro-mGzmB) in microcentrifuge tubes and incubated at 37 °C for 30 min while shaking at 400 rpm. In the case of inhibited GzmB assays, active human (or mouse) GzmB was first incubated with compounds **6-7** (or DMSO) for 1 h at 37 °C, then treated with compounds **8-10** for 30 min while shaking at 400 rpm. Protein samples were analyzed by SDS-PAGE and in-gel fluorescence scanning for Cy5 signal.

**Figure 3A-B**: (A) In a 12-well cell culture plate, $1\times10^6$ NK-92 cells were treated with different concentrations of compounds **8-10** (or DMSO) and incubated at 37 °C (5% $CO_2$) for 2 h. (B) In falcon tubes, $2\times10^6$ of NK-92, MDA-MB-231, Jurkat, and HEK cells were collected, pelleted, and resuspended in 300 μL of their respective fresh cell culture media. Then, 0.6 μL of 2.5 mM compound **10** was added (final working concentration of 5 μM), mixed well, and incubated at 37 °C (5% $CO_2$) for 2 h. Cells were then collected, washed with PBS (x3), and pelleted in microcentrifuge tubes. Cell lysates were then prepared according to the method described above, treated with 4x SDS reducing buffer, and subjected to electrophoresis.

**Figure 3D-E**: $5\times10^5$ of MDA-MB-231 cells were seeded in 6-well culture plates overnight. After confirming cell attachment under a microscope, media were removed, and fresh media (containing $2\times10^6$ of NK92 cells) containing **10** (5 μM) were added. Cells were co-incubated for different times at 37 °C (5% $CO_2$). Media containing NK-92 cells were removed, collected in microcentrifuge tubes, and washed with PBS 3 times. Attached MDA-MB-231 cells were washed with PBS 3 times and detached by trypsinization. (D) All cells were pelleted by centrifugation, lysed according to the method described above, 20 μg of proteins were treated with 4x SDS reducing buffer and subjected to electrophoresis. (E) All cells were pelleted by centrifugation, fixed in the dark at room temperature for 20 min, permeabilized at room temperature, and analyzed with flow cytometry.

#### D. Animal tumor model

NCI-Frederick and NCI-Bethesda are accredited by AAALAC International and follow the Public Health Service Policy for the Care and Use of Laboratory Animals. Animal care was provided in accordance with the procedures outlined in the “Guide for Care and Use of Laboratory Animals.” All animal experiments were performed following the regulation of the Animal Care and Use Committee (ACUC) of the National Cancer Institute (NCI). Five- to seven-week-old female Balb/c mice from Charles River Laboratories were injected with $5\times10^4$ 4T1 cells into the mammary fat pad #4 in 0.05 mL HBSS. Tumors were measured twice/week with calipers. Tumor volumes were calculated with $V(mm^3) = (D \times d^2)/6 \times 3.14$, where D and d represent the longest and shortest tumor axis, respectively. When tumors were about 100 $mm^3$, mice were randomized into treatment groups based on tumor volume and body weight using the StudyLog randomization software with

3-5 mice/group. For all studies, mice were monitored by daily observation and serial tumor measurements and body weights.
Mice were treated with 20 mg/kg BEC i.v., 10 mg/kg anti-PD1 i.p., 5 mg/kg anti-CTLA4 i.p. or a combination of 10 mg/kg anti-PD-1 and 5 mg/kg anti-CTLA4. All the treatments were administered into living mice bearing 4T1 tumor every three days for three times (q3dx3). Dosing was administered at 0.01 ml/g body weight. On day 9 post-treatment, the mice were injected with 2 nmol/g GzmB qABP i.v., followed by fluorescence imaging at 2, 4, 8, and 24 h. Following the last imaging time point, mice were euthanized, and tissues were collected for ex vivo imaging.

##### E. In vivo and ex vivo fluorescence imaging (FLI)

All fluorescence imaging (FLI) were performed using Xenogen IVIS Spectrum (Revvity Health Sciences, Inc., Hopkinton, Massachusetts), which was previously owned by PerkinElmer. Prior to in vivo or ex vivo FLI, quality control of the IVIS system was performed with XFM-2X fluorescent phantom mouse (Revvity Health Sciences, Inc., Hopkinton, Massachusetts). The total radiant efficiency of AF680 phantom was $1.44 \times 10^9 \pm 5.7 \times 10^7$ at 40% threshold for the entire study. Both in vivo and ex vivo FLI of mice including dye phantom was performed with Ex/Em: 640/680 nm. The wavelength difference between excitation and emission was large enough to prevent signal contamination by mixing. The maximum exposure time was fixed at 120 seconds to avoid background noise. In vivo imaging used 8 binning, 2 f-stop, and 1.5 cm sample height, whereas ex vivo imaging used 8 binning, 2–4 f-stop, and 0.5 cm sample height. The quantified data are presented as radiant efficiency or normalized radiant efficiency by area.

##### F. Flow cytometry analysis

All flow cytometry data were collected on BD LSRFortessa and FACSymphony A5 Cell Analyzers and analyzed with FlowJo software. Briefly, cells were processed according to the intended protocols for each experiment. Thereafter, cells were pelleted, fixed in the dark for 20 min at room temperature (Fixation buffer: BioLegend #420801), and pelleted again by centrifugation. Fixed cells were then resuspended in 1x intracellular staining perm wash buffer (BioLegend #421002), incubated for 10 min at room temperature, and pelleted by centrifugation. Fixed/Permeabilized cells were resuspended 100 μL 1x intracellular staining perm wash buffer and incubated with FITC-conjugated CD8 antibody (BioLegend #100706) at 0 °C for 1 h. After centrifugation, cell pellets were washed with 1x intracellular staining perm wash buffer (twice) and resuspended in the appropriate amount of cell staining buffer (BioLegend # 420201) and analyzed with flow cytometry.

##### G. Immunofluorescence (IFC) staining of tumor sections

Pieces of harvested tumors were fixed in 4% formaldehyde solution for 48h at room temperature, washed several times with PBS and stored in 70% ethanol before being paraffin-embedded. Tumor sections were then transferred to slides and stored before further analysis. Paraffin-embedded tumor sections were deparaffinated and rehydrated according to standard protocols. Briefly, slides were sequentially submerged in xylenes (5 min, x2), 100% ethanol (3 min, x2), 95% ethanol, 80% ethanol, 70% ethanol and 50% ethanol (1 min each) and finally, water (1 min). Slides were then submerged in a sodium citrate buffer pH 6.0 and incubated in boiling water (95-100 °C) for 20 min for antigen retrieval. After cooling down to room temperature, slides were submerged in PBST (2 min, x2) and blocked with 1% BSA in PBST for 30 min. Tissues were then incubated with primary antibody solution (diluted to manufacturer recommended concentrations) overnight at 4 °C. After washing with PBST (2 min, x4), tissues were incubated with secondary antibody (diluted to manufacturer recommended concentrations) for 1 h at room temperature and washed again with PBST (2 min). Tissues were then stained with DAPI for 30 min, washed with PBST (2 min, x3) and mounted with ProLong™ Diamond Antifade Mountant (Life Technologies #P36965). Slide was allowed to dry overnight at 4 °C and then imaged with confocal microscopy. For Cy5 imaging, tissues were directly stained with DAPI after deparaffination and rehydration stage.

#### H. Synthesis of compounds

$^{1}H$, $^{13}C$ and $^{31}P$ spectra were acquired on a 500 MHz NMR in $CDCl_3$ or $CD_3OD$ at 25 °C. The $^{1}H$ chemical shifts are given in parts per million (δ) with respect to an internal tetramethylsilane (TMS, δ 0.00 ppm) standard. NMR data are reported in the following format: chemical shifts (multiplicity (s = singlet, d = doublet, t = triplet, m = multiplet), coupling constants [Hz], integration). LCMS data were acquired on Agilent 1200 series LC/MS or Shimadzu Prominence-i LC-2030s instrument whereas HRMS data were obtained on an Agilent Q-TOF LC/MS/MS instrument.

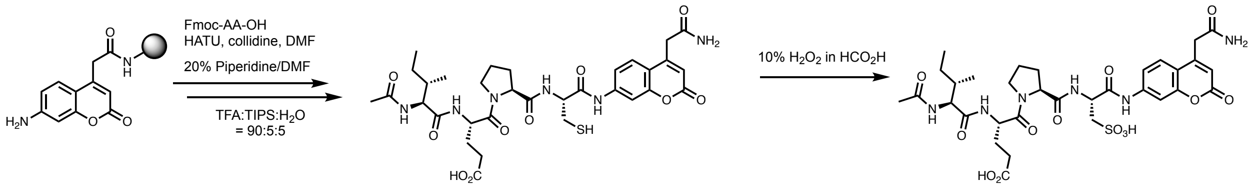


Scheme S1. Synthesis of fluorogenic substrates.

##### Solid-phase peptide synthesis of fluorogenic substrates (Compounds 1-4)

An appropriate amount of rink amide (60 to 80 mg) resin was weighed out in the SPPS cartridge. To the resin was added DMF and swelled for 1 h. Then 20% piperidine/DMF solution was added to the resin and rocked for 20 min to deprotect the Fmoc group on rink amide. This Fmoc deprotection sequence was repeated twice more for 20 min each and then the resin was washed three times with DMF.

In a falcon tube, 5 eq of Fmoc-ACC-OH was added to DMF followed by 5 eq HATU and 5 eq 2,4,6-collidine. The solution was mixed very well to ensure no solid was left and then, left at room temperature for 5 minutes. This reaction mixture was then added to resin in the SPPS cartridge and rocked at room temperature overnight. Next day, the same sequence was repeated by adding a solution containing 5 eq Fmoc-ACC-OH, HATU and 2,4,6-collidine to the resin and rocked for overnight again. Next day, the resin was washed three times with DMF. Then, a solution consisting of 3:2 Acetic anhydride/pyridine was added to the resin and rocked for an hour to cap the unreactive amine of the rink amide resin. Resin was thoroughly washed with DMF and the loading of Fmoc-ACC on the rink amide resin was calculated. Fmoc-ACC coupled rink amide resin was treated with 20% piperidine/DMF two times for 20 minutes each. Now, the resin was ready to undergo sequential amino acid coupling followed by Fmoc-deprotection. For each amino acid coupling, 3 eq of amino acid (with respect to the calculated loading efficiency), 3 eq of HATU and 3 eq of 2,4,6-collidine were premixed in DMF in falcon tubes and the resulting solution mixture was added to the resin and rocked for 1.5-3hrs depending on the nature of the amino acid. Acetylated isoleucine was mostly used as the last amino acid to complete the sequence.

After the solid phase peptide synthesis was complete, the peptide was washed with DMF (two times) followed by DCM (two times). Then, the peptide was cleaved from the resin using TFA/TIPS/$H_2O$ (18:1:1) for peptides containing a single cysteine and TFA/phenol/water/TIPS (88:5:5:2) for peptides containing two or more cysteines. In each case, the resin was rocked with the cleavage solution twice for 45 min to an hour each. The cleaved solution was drained to a flask and concentrated under reduced pressure to obtain the crude residue, which was further subjected to oxidation without further purification. The peptides with two or more free cysteines are prone to disulfide bond formation and needs to be protected from too much exposure to air or light and subjected to the oxidation as soon as possible.

**Oxidation of thiols to corresponding sulfonic acid:**

10% solution of hydrogen peroxide (32%) in formic acid was prepared and left in dark for an hour for the formation of performic acid. The round bottom flask containing the crude peptide was cooled to 0 °C using an ice-water bath and 10% $H_2O_2$ in $HCO_3H$ solution (6 mL per mmol of peptide) was carefully added to the flask. The reaction mixture was stirred at 0 °C for two hours and then concentrated under air or using genevac. The concentrated residue was purified by reverse phase HPLC.

**Compound 1:** HRMS (Q-TOF) m/z: $[M+H]^+$ Calcd for $C_{33}H_{43}N_6O_{12}^+$ 715.2939; Found 715.2993.
**Compound 2:** HRMS (Q-TOF) m/z: $[M+H]^+$ Calcd for $C_{32}H_{43}N_6O_{13}S^+$ 751.2609; Found 751.2658.
**Compound 3:** HRMS (Q-TOF) m/z: $[M+H]^+$ Calcd for $C_{32}H_{43}N_6O_{13}S^+$ 751.2609; Found 751.2611.
**Compound 4:** HRMS (Q-TOF) m/z: $[M+H]^+$ Calcd for $C_{31}H_{43}N_6O_{14}S_2^+$ 787.2279; Found 787.0964.

**Synthesis of intermediate peptides:**

Intermediate peptides were synthesized through solid-phase peptide synthesis (SPPS) following subsequent Fmoc-deprotection strategy. Briefly, 2-chlorotrityl resin was first swelled in DCM for 30 minutes in a SPPS vessel. After flushing the DCM, solution of P1 amino acid (3 eq) and collidine (5 eq) in DCM was added and the vessel was rocked overnight on a shaking platform at room temperature (Note: small amounts of DMF were added if the P1 amino acid showed solubility issues). After flushing the solution, resin was washed with DCM (x2), 6 mL of blocking agent (DCM:MeOH:DIPEA = 17:2:1) was added and the vessel was rocked at room temperature for 30 minutes. After flushing the solution, resin was washed sequentially with DCM (x2) and DMF (x2). Thereafter, subsequent Fmoc deprotection and amino acid coupling reactions were performed to build the intermediate peptide (coupling conditions: Amino Acid = 3 eq; HCTU = 5 eq; 2,4,6-collidine = 5 eq). After solid-phase synthesis was completed, peptide was cleaved with 20% hexafluoroisopropanol (HFIP) in DCM for 30 minutes (x2). Collected mixture was dried under reduced pressure, purified with reverse phase HPLC, fractions containing intermediate peptides were lyophilized and characterized with LCMS.

**Synthesis of 5:** Compound **5** was synthesized following the previously reported protocol.[1] Analytical data matched with those previously reported. HRMS (Q-TOF) m/z: $[M+H]^+$ Calcd for $C_{33}H_{44}N_4O_{11}P^+$ 703.2744; Found 703.1578.

[1] Mahrus, S.; Craik, C. S. *Chem. Biol.* **2005**, *12*, 567-577.

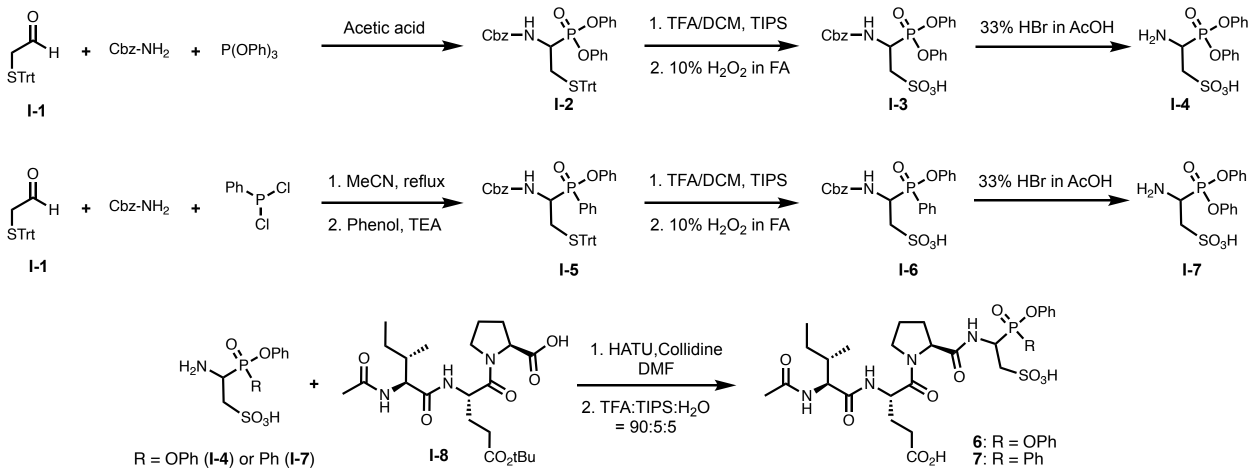


Scheme S2. Synthesis of phosphonate and phosphinate GzmB inhibitors **6** and **7**.

**Compound I-1.** Compound **I-1** was synthesized following previously reported protocols and its analytical data matched with previous reports.[2] $^1$H NMR (500 MHz, $CDCl_3$) δ 8.89 (t, *J* = 2.8 Hz, 1H), 7.52 – 7.46 (m, 4H), 7.39 – 7.24 (m, 12H), 3.13 (dd, *J* = 2.8, 1.3 Hz, 2H).

**Compound I-2.** A round bottom flask, equipped with a magnetic bead, was charged with **6a** (1g, 6.57 mmol), benzyl carbamate (780 mg, 5.26 mmol), triphenyl phosphite (1.4 mL, 5.26 mmol) and glacial acetic acid 15 mL. The resulting reaction mixture was stirred at room temperature for 30 minutes, then heated to 80 ˚C and stirred at that temperature for 2 h. Excess acetic acid was neutralized with saturated $NaHCO_3$, mixture was extracted with 200 mL EtOAc and washed with brine (200 mL, x1). Organic layer was dried over anhydrous $Na_2SO_4$, concentrated under reduced pressure, and purified by combi-flash silica gel chromatography to obtain **I-2** as a thick yellow oil (2g, 55%). $^1$H NMR (500 MHz, $CDCl_3$) δ 7.56 – 7.42 (m, 7H), 7.42 – 7.20 (m, 23H), 7.13 – 6.97 (m, 4H), 5.38 (d, *J* = 10.3 Hz, 1H), 5.15 (d, *J* = 12.1 Hz, 3H), 4.42 (dtd, *J* = 17.2, 10.4, 3.8 Hz, 1H), 2.95 – 2.75 (m, 2H).

**Compound I-3.** A round bottom flask, equipped with a magnetic bead was charged with **I-2** (2 g, 2.92 mmol) and the compound was dissolved in 40 mL DCM. Then 5 ml TFA was added dropwise followed by triisopropylsilane (2.5 mL). The resulting reaction mixture was stirred at room temperature for 1 hour. Then, the reaction was concentrated to dryness under reduced pressure and obtained residue was used for the next step without further purification. 10% solution of hydrogen peroxide (32%) in formic acid was prepared and left in dark for an hour for the formation of performic acid. The round bottom flask containing the crude thiol was cooled to 0 °C using an ice-water bath and 10% $H_2O_2$ in $HCO_3H$ solution (6 mL per mmol of crude thiol) was carefully added to the flask. The reaction mixture was stirred at 0 °C for two hours and then concentrated under air or using genevac. The concentrated residue was purified by reverse phase HPLC to obtain **I-3** as a thick yellow oil (590 mg, 41%). $^1$H NMR (500 MHz, MeOD) δ 7.44 – 7.24 (m, 9H), 7.24 – 7.09 (m, 6H), 5.18 (d, *J* = 12.6 Hz, 1H), 5.07 (td, *J* = 14.5, 8.2 Hz, 2H), 3.41 (ddd, *J* = 14.2, 8.7, 2.2 Hz, 1H), 3.37 – 3.33 (m, 1H). LCMS m/z $[M+H]^+$ Calcd for $C_{22}H_{23}NO_8PS^+$ 492.09; Found 492.1.

**Compound I-4.** To a round bottom flask, equipped with a magnetic bead were added **I-3** (145 mg, 0.295 mmol) and 33% HBr in AcOH. The reaction was stirred for 1 hour, then concentrated

[2] Asahina, Y.; Hojo, H. J. Org. Chem. 2020, 85, 1458-1465.

under air and purified by reverse phase HPLC to afford **I-4** as thick yellow oil (76 mg, 72%). $^1$H NMR (500 MHz, MeOD) δ 7.41 (td, *J* = 8.0, 2.4 Hz, 4H), 7.34 – 7.14 (m, 6H), 4.43 (ddt, *J* = 16.3, 11.9, 2.0 Hz, 1H), 3.59 – 3.50 (m, 1H), 3.39 – 3.32 (m, 1H). LCMS m/z $[M+H]^+$ Calcd for $C_{14}H_{17}NO_6PS^+$ 358.05; Found 358.1.

**Compound I-5.** To a solution of **I-1** (2g, 6.29 mmol, 1 eq) in 30 mL MeCN were added benzyl carbamate (950 mg, 6.29 mmol, 1 eq) and dichlorophenyl phosphine (1.28 mL, 9.43 mmol, 1.5 eq). The mixture was refluxed for 45 minutes and then brought to room temperature. A solution of phenol (886 mg, 9.43 mmol, 1.5 eq) in 10 mL MeCN was added to the solution followed by dropwise addition of triethylamine (2.63 mL, 18.86 mmol, 3 eq) and the mixture was stirred at room temperature for 2h. Excess solvent was removed under reduced pressure and the mixture was dissolved in 100 mL EtOAc, washed with saturated $NaHCO_3$ (100 mL, x1) and brine (100 mL, x1). Organic layer was dried over anhydrous $Na_2SO_4$, evaporated under reduced pressure, and subjected to purification on a combi-flash silica gel chromatography. Fractions containing **I-5** were collected and taken to the next step without further purification (Note: purification of **I-5** was attempted by crystallization of the fractions collected from chromatography with limited success).

**Compound I-6.** To a solution of intermediate **I-5** in 30 mL DCM were added 1 mL TFA and 0.5 mL TIPS. Solution was stirred at room temperature for 1 h and excess solvents were removed under reduced pressure. The flask was put on ice-water bath, 6 mL mixture of 10% $H_2O_2$ in formic acid (previously kept in dark for 1 h at room temperature) was added and the mixture was stirred at 0 °C for 2 h. Solution was concentrated under air and purified by reverse phase HPLC to obtain compound **I-5** as a yellow oil (200 mg, 35%). $^1$H NMR (400 MHz, MeOD) δ 7.82 – 7.72 (m, 2H), 7.55 – 7.46 (m, 1H), 7.42 – 7.31 (m, 2H), 7.28 – 7.09 (m, 6H), 7.09 – 6.88 (m, 4H), 5.04 – 4.87 (m, 2H), 4.84 (d, *J* = 5.4 Hz, 1H), 4.73 (d, *J* = 12.6 Hz, 1H), 3.33 (ddd, *J* = 14.4, 7.1, 1.9 Hz, 1H), 3.20 – 3.06 (m, 1H). LCMS m/z: $[M+H]^+$ Calcd for $C_{22}H_{23}NO_7PS^+$ 476.09; Found 476.1.

**Compound I-7.** Compound **I-6** (200 mg, 0.42 mmol) was added to a vial, 1 mL of 33% HBr in AcOH was added, and the mixture was stirred at room temperature. Upon completion, monitored by LCMS, mixture was concentrated under air and purified by reverse phase HPLC to obtain compound **I-7** as a white solid (50 mg, 34.7%). $^1$H NMR ($CD_3OD$) δ 7.82–7.92 (m, 2H), 7.61–7.70 (m, 1H), 7.49–7.59 (m, 2H), 7.17–7.26 (m, 2H), 7.03–7.17 (m, 3H), 4.15–4.35 (m, 1H), 3.34–3.44 (m, 1H), 2.88–2.95 (m, 1H). HRMS (Q-TOF) m/z: $[M + H]^+$ Calcd for $C_{14}H_{17}NO_5PS^+$ 342.0565; Found 342.0579.

**Ac-Ile-Glu(tBu)-Pro-OH (Compound I-8).** Tripeptide **I-8** was synthesized through solid-phase peptide synthesis following general procedure outlined above. LCMS m/z: $[M + H]^+$ Calcd for $C_{22}H_{38}N3O_7^+$ 456.6; Found 456.3.

**Compound 6.** To a round bottom flask, equipped with a magnetic stir bar were added **I-4** (40 mg, 0.112 mmol), Ac-Ile-Glu(tBu)-Pro-OH (**I-8**, 17 mg, 0.037 mmol) and HATU (71 mg, 0.187 mmol) and dissolved in 3 mL DMF. Then collidine (25 μL, 0.187 mmol) was added and the solution was stirred at room temperature for 2h (monitored for completion with HPLC). Then the solution was concentrated on genevac and purified with reverse phase HPLC. Fractions containing intermediate peptide were lyophilized (6 mg, m/z $[M+H]^+$ Calcd for $C_{36}H_{51}N_4O_{12}PS^+$ 795.3; Found 795.2), dissolved in 5 mL (DCM:TFA = 4:1) and 0.25 mL TIPS and stirred at room temperature for 1h. After completion (monitored by LCMS), solvents were removed under air and product was purified by reverse phase HPLC to obtain compound **6** as a white solid (1.5 mg, 5.6%). $^1$H NMR (500 MHz, MeOD) δ 7.34 (qd, *J* = 7.4, 3.9 Hz, 4H), 7.24 – 7.18 (m, 5H), 7.16 (m, 1H), 5.42 – 5.22 (m, 1H), 4.75 (dd, *J* = 9.2, 4.7 Hz, 1H), 4.63 – 4.53 (m, 1H), 4.19 (dd, *J* = 7.9, 3.8 Hz, 1H), 3.79 (dddd, *J* = 17.3, 11.9, 8.6, 6.3 Hz, 2H), 3.42 (ddt, *J* = 11.4, 8.3, 3.8 Hz, 1H), 2.50 – 2.36 (m, 2H),

2.27 – 2.03 (m, 2H), 1.96 – 1.86 (m, 1H), 1.52 (tdd, *J* = 14.9, 7.4, 3.7 Hz, 1H), 1.24 – 1.10 (m, 1H), 0.90 (td, *J* = 7.1, 4.5 Hz, 5H). HRMS (Q-TOF) m/z $[M+H]^+$ Calcd for $C_{32}H_{44}N_4O_{12}PS^+$ 739.2414; found 739.2508.

**Compound 7.** To a solution of intermediate **I-7** (30 mg, 0.09 mmol) in 1 mL DMF were added Ac-IE(OtBu)P-OH (**I-8**, 18 mg, 0.04 mmol), HATU (70 mg, 0.18 mmol) and Collidine (25 μL, 0.19 mmol). Reaction mixture was stirred at room temperature for 1 h, concentrated on genevac and purified by reverse phase HPLC. Fractions containing intermediate peptide were combined (LCMS m/z: $[M+H]^+$ Calcd for $C_{36}H_{52}N_4O_{11}PS^+$ 779.31; Found 779.3), lyophilized and dissolved in 5 mL of DCM:TFA (4:1) containing 5% TIPS. Mixture was stirred at room temperature for 1 h, completion monitored by LCMS. Mixture was concentrated under air and purified by reverse phase HPLC to obtain compound **7** as a white solid (1.35 mg, 4.7%). $^1$H NMR (500 MHz, MeOD) δ 7.93 – 7.76 (m, 2H), 7.57 – 7.33 (m, 3H), 7.23 – 6.93 (m, 5H), 5.22 (t, *J* = 12.8 Hz, 1H), 5.04 (d, *J* = 9.1 Hz, 1H), 4.70 – 4.49 (m, 1H), 4.47 – 4.17 (m, 1H), 4.20 – 3.95 (m, 1H), 3.66 (ddd, *J* = 39.1, 20.1, 11.3 Hz, 2H), 3.41 – 3.28 (m, 1H), 2.44 – 2.14 (m, 3H), 2.12 – 1.84 (m, 6H), 1.84 – 1.63 (m, 3H), 1.47 – 1.40 (m, 1H), 1.17 – 0.99 (m, 1H), 0.95 – 0.75 (m, 6H). HRMS (Q-TOF) m/z: $[M]^+$ Calcd for $C_{32}H_{44}N_4O_{11}PS^+$ 723.2465; Found 723.2471.

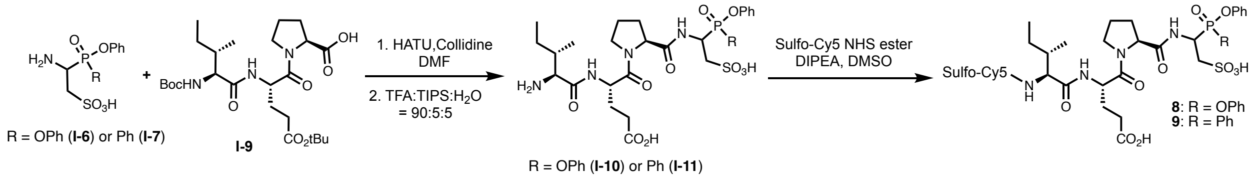


Scheme S3. Synthesis of phosphonate and phosphinate GzmB ABPs **8** and **9**.

**Boc-Ile-Glu(tBu)-Pro-OH (Compound I-9):** Tripeptide **I-9** was synthesized through solid-phase peptide synthesis following general procedure outlined above. LCMS m/z: $[M + H]^+$ Calcd for $C_{25}H_{44}N_3O_8^+$ 514.6; Found 514.3.

**Compound I-10.** To a stirred solution of tripeptide Boc-Ile-Glu(tBu)-Pro-OH in dry DMF under $N_2$ was added **I-6** followed by HATU and 2,4,6-collidine at 0 °C. The reaction was raised to room temperature and stirred at room temperature for 18 hrs. Then the reaction was concentrated under reduced pressure and the crude compound was used for the next step without further purification. The crude intermediate was suspended in dry DCM and cooled to 0 °C. TFA was added dropwise to the reaction mixture, followed by TIPS. The reaction was stirred at rt for 1 h and then concentrated under reduced pressure. The obtained residue was purified by reverse phase HPLC. $^1$H NMR (500 MHz, MeOD) δ 7.43 – 7.31 (m, 5H), 7.28 – 7.10 (m, 6H), 6.77 (dd, *J* = 10.8, 7.8 Hz, 1H), 5.43 (t, *J* = 14.1 Hz, 1H), 5.37 – 5.22 (m, 1H), 5.22 – 5.10 (m, 1H), 4.83 – 4.72 (m, 3H), 4.70 – 4.46 (m, 2H), 4.09 (dd, *J* = 8.4, 4.6 Hz, 1H), 3.95 – 3.64 (m, 3H), 3.65 – 3.35 (m, 4H), 3.27 – 3.06 (m, 2H), 2.63 – 2.33 (m, 4H), 2.35 – 2.05 (m, 4H), 2.06 – 1.73 (m, 6H), 1.73 – 1.37 (m, 3H), 1.39 – 1.16 (m, 2H), 1.16 – 0.87 (m, 6H). LCMS m/z $[M+H]^+$ Calcd for $C_{30}H_{42}N_4O_{11}PS^+$ 697.23; Found 697.2.

**ABP 8.** In an Eppendorf tube, **I-10** was dissolved in 50 uL DMSO and to that was added Sulfo-Cy5-NHS ester solution in DMSO followed by DIPEA. The Eppendorf tube was immediately wrapped in Al-foil and shaken overnight at room temperature. Next day, the reaction mixture was diluted with 1 ml of $H_2O$, purified by reverse phase HPLC and product lyophilized to obtain ABP **8** as a blue solid. HRMS (Q-TOF) m/z $[M]^+$ Calcd for $C_{63}H_{78}N_6O_{18}PS_3^+$ 1333.4272; Found 1335.4418 (Deconvoluted: 1334.44).

**ABP 9.** To a solution of intermediate **I-7** (30 mg, 0.09 mmol) in 1 mL DMF were added Boc-IE(tBu)P-OH (**I-9**, 40 mg, 0.08 mmol), HATU (150 mg, 0.39 mmol) and Collidine (50 μL, 0.38 mmol). Reaction mixture was stirred at room temperature for 1h, concentrated on genevac and purified by reverse phase HPLC. Fractions containing intermediate peptide were combined, lyophilized ($^1$H NMR attached; LCMS m/z: $[M+H]^+$ Calcd for $C_{39}H_{58}N_4O_{12}PS^+$ 837.35; Found 837.3) and dissolved in 5 mL of DCM:TFA (4:1) containing 5% TIPS. The mixture was stirred at room temperature for 1 h, complete deprotection observed by LCMS. The mixture (**I-11**) was concentrated under air, redissolved in 500 μL DMSO, and added to a vial containing 7 mg sulfo-Cy5-NHS (9.3 μmol) and the vial was covered with aluminum foil to protect from light. DIPEA (50 μL) was then added to the solution and stirred at room temperature for 2 h. The mixture was then purified by reverse phase HPLC to obtain compound **9** as a blue solid (3.95 mg, 32%). HRMS (Q-TOF) m/z: $[M]^+$ Calcd for $C_{63}H_{78}N_6O_{17}PS_3^+$ 1317.4323; Found 1318.4296 (Deconvoluted: 1318.43).

**Compound I-12.** Tyramine (2g, 14.58 mmol) was dissolved in 30 mL of 1:1 Water:1,4-Dioxane and $NaHCO_3$ (3.67 g, 43.74 mmol) was added to the solution. Then allyl chloroformate (1.86 mL, 17.5 mmol) was added dropwise and the solution was stirred at room temperature overnight. Mixture was then extracted with 200 mL EtOAc, washed with 1M HCl (x1, 200 mL) and brine (x1, 200 mL), dried over anhydrous $Na_2SO_4$ and concentrated under reduced pressure. Product 10a was isolated with combi-flash silica gel chromatography as a dense liquid which solidified upon standing (2.5 g, 77.4%). $^1$H NMR ($CDCl_3$) δ 7.00–7.06 (m, 2H), 6.74–6.81 (m, 2H), 5.79–6.00 (m, 1H), 5.63 (s, 1H), 5.12–5.35 (m, 2H), 4.77 (s, 1H), 4.48–4.64 (m, 2H), 3.30–3.47 (q, 2H), 2.67–2.78 (t, 2H). LCMS m/z: $[M+H]^+$ Calcd for $C_{12}H_{16}NO_3^+$ 222.1; Found 222.0.

**Compound I-13.** To a solution of **I-1** (2g, 6.29 mmol) in 30 mL MeCN were added benzyl carbamate (950 mg, 6.29 mmol) and dichlorophenyl phosphine (1.28 mL, 9.43 mmol, 1.5 eq). The mixture was refluxed for 45 minutes and then brought to room temperature. A solution of **I-12** (3 g, 13.57 mmol) in 10 mL MeCN was added to the mixture, stirred for 15 minutes, then 4 mL triethylamine was added dropwise, and the mixture was stirred at room temperature overnight. Excess solvents were evaporated under reduced pressure and mixture was dissolved in 200 mL EtOAc, washed with $NaHCO_3$ (200 mL, x1) and brine (200 mL, x1). Organic layer was dried over $Na_2SO_4$, evaporated under reduced pressure, and subjected to purification on a combi-flash silica gel chromatography. Fractions containing compound **I-13** were collected and taken to the next step without further purification.

**Compound I-14.** To a solution of intermediate **I-13** (~1.5 g with impurities) in 20 mL DCM were added 2 mL TFA and 1 mL TIPS. Solution was stirred at room temperature for 2h and excess solvents were removed under reduced pressure. The flask was put on ice-water bath, 6 mL mixture of 10% $H_2O_2$ in formic acid (previously kept in dark for 1h at room temperature) was added. Flask was brought to room temperature and stirred for 1h. Solution was concentrated under air and purified by reverse phase HPLC and fractions containing compound **I-14** were lyophilized to obtain product as a white solid (300 mg, 26%). $^1$H NMR (500 MHz, MeOD) δ 7.76 (t, *J* = 9.6 Hz, 2H), 7.55 – 7.46 (m, 1H), 7.41 – 7.22 (m, 2H), 7.18 (m, 4H), 7.08 – 6.85 (m, 5H), 5.85 – 5.71 (m, 1H), 5.15 (dtd, *J* = 17.1, 2.0, 1.3 Hz, 1H), 5.09 – 5.03 (m, 1H), 4.93 (dd, *J* = 21.2, 12.4 Hz, 1H), 4.85 – 4.77 (m, 1H), 4.72 (d, *J* = 12.6 Hz, 1H), 4.37 (d, *J* = 5.1 Hz, 2H), 3.15 (t, *J* = 7.3 Hz, 2H), 2.60 (t, *J* = 7.3 Hz, 2H). LCMS m/z: $[M+H]^+$ Calcd for $C_{28}H_{32}N_2O_9PS^+$ 603.16; Found 603.2.

**Compound I-15.** Compound **I-14** (300 mg, 0.03 mmol) was added to a vial and the vial was brought to 0 °C in an ice-water bath. Then 1 mL of 33% HBr in AcOH was added and the mixture was stirred at 0 °C for 1 h (Note: If the reaction is carried out at room temperature, both Cbz- and Alloc-groups are deprotected). Mixture was then concentrated under air and purified by reverse phase HPLC. Lyophilization of fractions containing intended product yielded compound **I-15** as a white solid (140 mg, 60%). $^1$H NMR (400 MHz, MeOD) δ 7.92 – 7.82 (m, 2H), 7.71 – 7.61 (m, 1H), 7.54 (tt, *J* = 7.7, 4.0 Hz, 2H), 7.11 – 7.02 (m, 4H), 5.78 (ddt, *J* = 16.2, 10.6, 5.4 Hz, 1H), 5.22 – 4.92 (m, 2H), 4.37 (d, *J* = 5.3 Hz, 1H), 4.33 – 4.12 (m, 2H), 3.43 – 3.33 (m, 1H), 3.16 (td, *J* = 7.1, 3.0 Hz, 2H), 2.97 – 2.85 (m, 1H), 2.62 (t, *J* = 7.2 Hz, 2H). LCMS m/z: $[M+H]^+$ Calcd for $C_{20}H_{26}N_2O_7PS^+$ 469.12; Found 469.1.

**Compound I-16.** Compound **I-15** (60 mg, 0.13 mmol) was dissolved in 1 mL DMF. Boc-IE(OtBu)P-OH (70 mg, 0.14 mmol), HATU (260 mg, 0.684 mmol) and Collidine (90 μL, 0.681 mmol) were added and the mixture was stirred at room temperature for 1h. Mixture was concentrated on genevac and purified by reverse-phase HPLC. Lyophilization of fractions containing intended product yielded compound **I-16** as a white solid (45 mg, 36.3%). LCMS m/z: $[M+H]^+$ Calcd for $C_{45}H_{67}N_5O_{14}PS^+$ 964.41; Found 964.55.

**Compound I-17.** Compound **I-16** (45 mg, 0.096 mmol) was dissolved in 5 mL DCM:TFA (4:1) and stirred at room temperature for 1 h (monitored for completion with LCMS). Solvents were removed under air and mixture was purified with reverse-phase HPLC to obtain the deprotected intermediate a white solid (18 mg, 47.7% yield, LCMS m/z: [M+H]+ Calcd for $C_{36}H_{51}N_5O_{12}PS+$ 808.3; Found 808.5). Intermediate was transferred to a vial, dissolved in 1 mL DMF and Sulfo-Cy5-NHS (15 mg, 20 μmol) and 35 μL DIPEA were added to the solution. The vial was covered with aluminum foil and solution was stirred at room temperature for 1h (monitored for completion with LCMS). Solution was then concentrated on genevac and purified with reverse-phase HPLC to yield **I-17** as blue solid (13 mg, 40%). LCMS m/z: $[M]^+$ Calcd for $C_{69}H_{88}N_7O_{19}PS_3^+$ 1445.5; Found 1446.8.

**qABP 10.** A vial containing compound **I-17** (13 mg, 9 μmol) dissolved in 1 mL DMF was covered with aluminum foil. The mixture was set under $N_2$ atmosphere, $Pd(PPh_3)_4$ (20 mg, 17 umol) and 40 μL pyrrolidine were added and the mixture was stirred at room temperature under $N_2$ atmosphere for 30 minutes (completion confirmed with LCMS). Thereafter, QSY-21-NHS (7 mg, 8.6 μmol) and 30 μL DIPEA were added, and the mixture was stirred at room temperature under $N_2$ atmosphere for another 1h (monitored with LCMS). Solution was then concentrated on genevac and purified with reverse-phase HPLC to yield compound **10** as blue solid (2.5 mg, 14.4%, also isolated 2 mg of unreacted intermediate). HRMS (Q-TOF) m/z: $[M+H]^+$ Calcd for $C_{106}H_{118}N_{10}O_{21}PS_4^+$ 2025.7094; Found 2026.6998 (Deconvoluted: 2025.70).

**Compound 11:** Tetrapeptide Boc-Ile-Glu(tBu)-Pro-Cys(Trt)-$CONH_2$ was synthesized through SPPS with rink amide resin. Global deprotection with TFA followed by oxidation with 10% $H_2O_2$ in Formic Acid as described above yielded $H_2N$-Ile-Glu-Pro-Cya-$CONH_2$, which was dissolved in 100 uL DMSO and transferred to an Eppendorf tube containing sulfo-Cy5 NHS ester. Tube was covered with Al-foil, DIPEA was added and shaken overnight at room temperature. Mixture was diluted with 1:1 ACN:$H_2O$ and purified with reverse-phase HPLC to yield product **11** as a blue solid. HRMS (Q-TOF) m/z: $[M+H]^+$ Calcd for $C_{52}H_{71}N_7O_{16}S_3^+$ 1146.3580; Found 1146.4209.

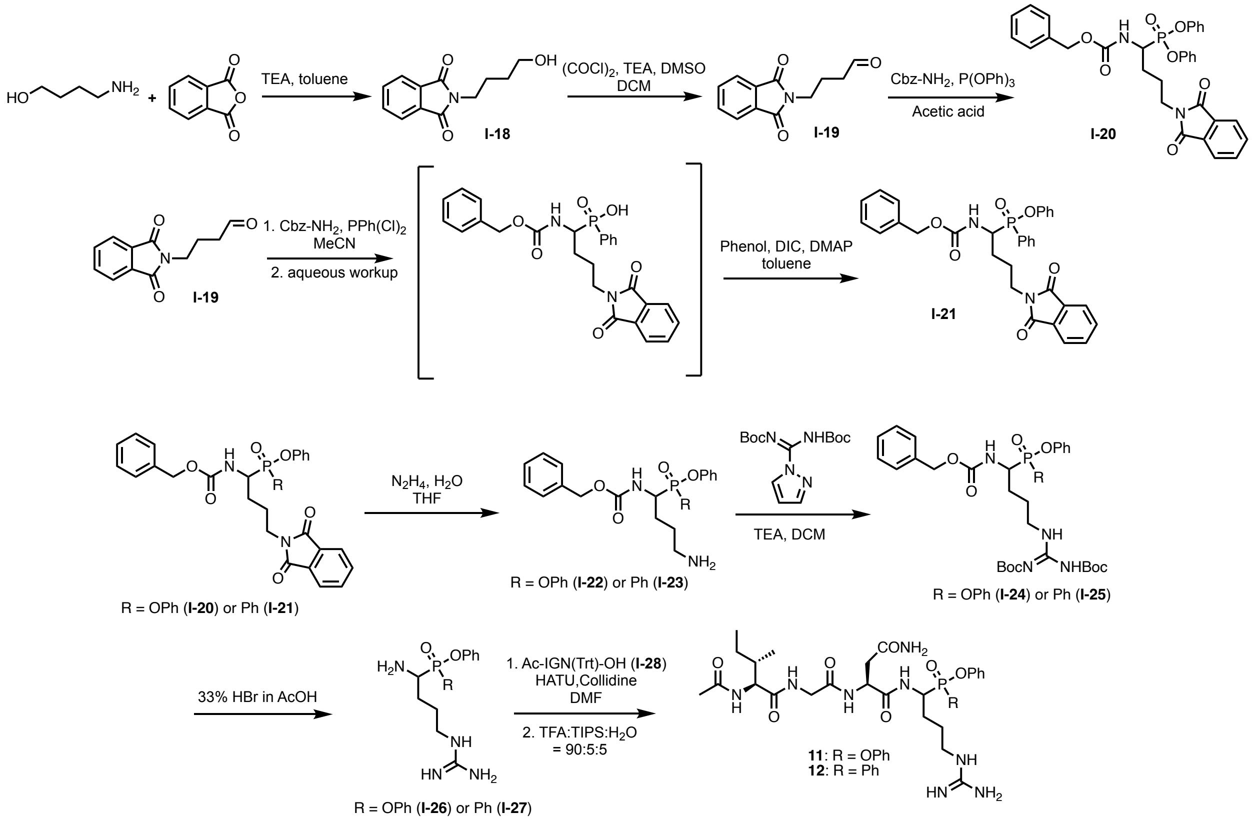


Scheme S4. Synthesis of phosphonate and phosphinate GzmA inhibitors **12** and **13**.

**Compounds I-24 and I-25:** The phosphonate and phosphinate cores **I-24** and **I-25** were synthesized following previously reported protocols.[3,4] Briefly, intermediate **I-18** obtained by the protection of phthalic anhydride with 4-aminobutanol was oxidized to the aldehyde **I-19** by Swern oxidation followed by a 3-component reaction with benzyl carbamate and triphenyl phosphite or dichlorophenyl phosphine to form the corresponding phosphonate (**I-20**) or phosphinate (**I-21**) intermediates. Selective deprotection of the phtalimide moiety generated the intermediate amines **I-22** and **I-23** before treatment Bis-Boc-pyrazolocarboxamidine to form **I-24** and **I-25**.

**Compound I-26.** Compound **I-26** was synthesized following previously reported protocols and its analytical data matched with previous reports.[4] LCMS m/z $[M+H]^+$ Calcd for $C_{17}H_{24}N_4O_3P^+$ 363.2; Found 363.2.

**Compound I-27.** 470 mg (0.69 mmol, 1 eq) of **I-25** was dissolved in 3.5 mL of 33% HBr in AcOH, and allowed to stir at room temperature for 2 h. Excess reagents were removed under air, and the crude was suspended in cold ether. After decanting the mother liquor, obtained precipitates were subjected to separation by reverse-phase HPLC. Purified fractions were lyophilized, yielding 102 mg (43%) of a white solid. $^1$H NMR (500 MHz, MeOD) δ 7.99 (ddd, *J* = 12.4, 8.1, 1.4 Hz, 2H), 7.75 (tdd, *J* = 5.0, 3.3, 1.7 Hz, 1H), 7.64 (td, *J* = 7.7, 3.9 Hz, 2H), 7.30 (qd, *J* = 7.2, 1.9 Hz, 2H), 7.26 – 7.10 (m, 3H), 4.17 (ddd, *J* = 10.3, 8.0, 5.6 Hz, 1H), 3.32 (s, 2H), 2.21 (dtt, *J* = 14.9, 11.8, 5.8 Hz, 1H), 2.08 – 1.92 (m, 2H), 1.86 (dtt, *J* = 18.9, 13.2, 6.2 Hz, 1H).

---

[3] Ji, S. et al *Org. Biomol. Chem.* **2023**, *21*, 6498-6502.
[4] Zhang, L. et al *JACS*, **2022**, *144*, 22493-22504.

**Ac-Ile-Gly-Asn(Trt)-OH (Compound I-28):** Tripeptide **I-28** was synthesized through solid-phase peptide synthesis following general procedure outlined above. LCMS m/z: $[M + H]^+$ Calcd for $C_{33}H_{39}N_4O_6^+$ 587.3; Found 587.3.

**Compound 11.** 35 mg (0.060 mmol, 1 eq) of **I-28**, 33.2 mg (0.066 mmol, 1.1 eq) of HATU, and 53 μL (0.30 mmol, 5 eq) of 2,4,6-collidine were added to a 20 mL scintillation vial containing 26 mg (0.072 mmol, 1.2 eq) of **I-26** dissolved in 2 mL of DMF. The reaction was allowed to stir at room temperature for 2 h, after which solvent was removed under vacuum. The crude was taken forward to the next step, in which 3 mL of a 2.5% DCM, 2.5% TIPS, 95% TFA solution was added and stirred for an hour. Excess reagents were removed under air and mixture was purified by reverse-phase HPLC. Desired fractions were collected and lyophilized, yielding 3.63 mg of a white solid (8.8%). HRMS (Q-TOF) m/z: $[M+H]^+$ Calcd for $C_{31}H_{46}N_8O_8P^+$ 689.3171; Found 681.3183.

**Compound 12.** 53 mg (0.09 mmol, 1 eq) of **I-28**, 37.6 mg (0.099 mmol, 1.1 eq) of HATU, and 59.5 μL (0.45 mmol, 5 eq) of 2,4,6,-trimethylpyridine were added to a 20 mL scintillation vial containing 52.9 mg (0.09 mmol, 1.2 eq) of **I-27** dissolved in 2 mL of DMF. The reaction was allowed to stir for 2 h, after which solvent was removed under vacuum. The crude was taken forward to the next step, in which 3 mL of a 2.5% DCM, 2.5% TIPS, 95% TFA solution was added and stirred for an hour. Excess reagents were removed under air and mixture was purified by reverse-phase HPLC. Desired fractions were collected and lyophilized, yielding 3.6 mg of a white solid (6.6%). HRMS (Q-TOF) m/z: $[M+H]^+$ Calcd for $C_{31}H_{46}N_8O_7P^+$ 673.3222; Found 673.3223.

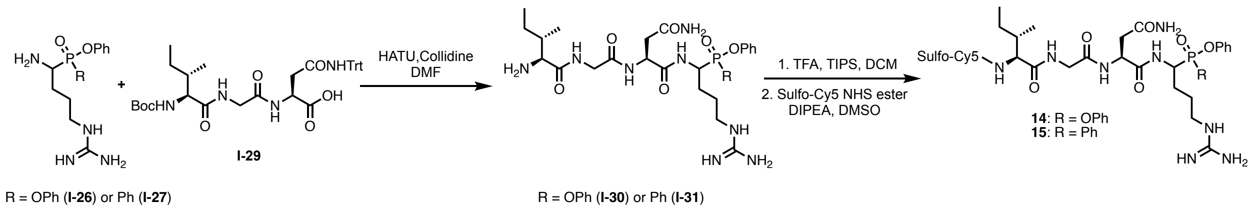


Scheme S5. Synthesis of phosphonate and phosphinate GzmA ABPs **14** and **15**.

**Boc-Ile-Gly-Asn(Trt)-OH (Compound I-29):** Tripeptide **I-29** was synthesized through solid-phase peptide synthesis following general procedure outlined above. LCMS m/z: $[M + H]^+$ Calcd for $C_{36}H_{45}N_4O_7^+$ 645.3 Found 645.3.

**Compound I-30.** 31.8 mg of **I-26** (0.084 mmol, 1.1 eq) was added to a solution of 48.9 mg of **I-29** (0.076 mmol, 1 eq) in 2 mL of anhydrous DMF. After addition of 32 mg HATU (0.084 mmol, 1.1 eq) and 50.8 μL of 2,4,6-trimethylpyridine (0.39 mmol, 5 eq), the reaction was allowed to run for 4 h, and progress monitored by LCMS. Solvent was removed by genevac and the crude was purified by reverse-phase HPLC. Desired fractions were collected and lyophilized, yielding 58 mg of a white solid (77%). LCMS m/z $[M+H]^+$ Calcd for $C_{53}H_{66}N_8O_9P^+$ 989.5; Found 990.3.

**Compound I-31.** 32 mg of **I-27** (0.0935 mmol, 1.1 eq) was added to a solution of 54.5 mg of **I-29** (0.085 mmol, 1 eq) and 35.5 mg of HATU (0.0935 mmol, 1.1 eq) in 2 mL of anhydrous DMF. After addition of 56.2 μL of 2,4,6-trimethylpyridine (0.43 mmol, 5 eq.), the reaction was allowed to run for 4 h, and progress monitored by LCMS. Solvent was removed by genevac and the crude was purified by reverse-phase HPLC. Desired fractions were collected and lyophilized, yielding 33 mg of a white solid (40%). LCMS m/z $[M+H]^+$ Calcd for $C_{53}H_{66}N_8O_8P^+$ 973.4; Found 973.4.

**ABP 14.** 22.5 mg of **I-30** (0.023 mmol, 1 eq) was dissolved in a solution of 10:2:1 DCM: TFA: TIPS and allowed to stir for 5 h with reaction progress monitored by LCMS. Excess reagents were removed under vacuum and the crude was taken forwards to the next step without purification. After dissolving the crude in 1 mL of anhydrous DMSO, 4.3 mg (0.0057 mmol, 0.25 eq) of Sulfo-Cy5-NHS ester and 40 μL (0.23 mmol, 10 eq) of DIPEA were added. The reaction vessel was covered in aluminum foil and allowed to stir overnight at room temperature under $N_2$. Purification was carried out by reverse-phase HPLC, yielding 1.95 mg (27%) of a blue solid. HRMS (Q-TOF) m/z: $[M+H]^+$ Calcd for $C_{62}H_{83}N_{10}O_{14}PS_2^{2+}$ 643.2629; Found 643.2651 (Deconvoluted 1284.51).

**ABP 15.** 16.5 mg of **I-31** (0.017 mmol, 1 eq) was dissolved in a solution of 10:2:1 DCM: TFA: TIPS and allowed to stir for 3 h with reaction progress monitored by LCMS. Excess reagents were removed under vacuum, the crude was dissolved in 1 mL of anhydrous DMSO and 9 mg (0.012 mmol, 0.70 eq) of Sulfo-Cy5-NHS ester and 30 μL (0.17 mmol, 10 eq) of DIPEA were added. The reaction vessel was covered in aluminum foil and allowed to stir overnight at room temperature under $N_2$. Purification was carried out by reverse-phase HPLC, yielding 1.42 mg (6.8%) of a blue solid. HRMS (Q-TOF) m/z: $[M+H]^+$ Calcd for $C_{62}H_{83}N_{10}O_{13}PS_2^+$ 635.2655; Found$^+$ 635.2674 (Deconvoluted 1268.52).

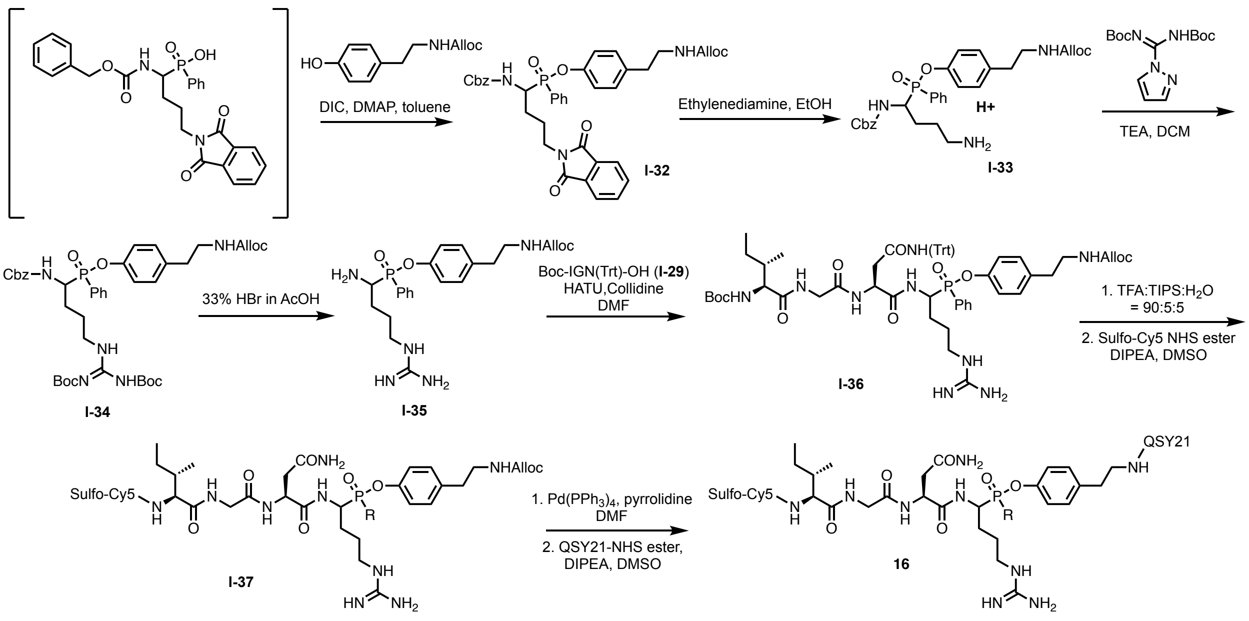


Scheme S6. Synthesis of phosphinate GzmA qABP **16**

**Compound I-32.** 3.34 g (6.8 mmol, 1 eq) of the intermediate phosphinic acid (Scheme S6) was dissolved in 15 mL of toluene. 1.65 g (7.48 mmol, 1.1 eq) of alloc-tyramine, 4.23 mL DIC (27.2 mmol, 4 eq), and 83.1 mg (0.68 mmol, 0.1 eq) of DMAP were added, a condenser was attached, and the reaction was stirred for 6h at 80 °C. Solvent was then removed via genevac, and the crude was suspended in DCM and subsequently washed x3 with each of the following: 1M $KHSO_4$, $H_2O$, $NaHCO_3$, and brine. Organic layer was dried over anhydrous $Na_2SO_4$, evaporated under reduced pressure, and subjected to purification with combi-flash silica gel chromatography, yielding 2.61 grams (45%) of yellow oil. $^1H$ NMR (400 MHz, MeOD) δ 7.88 – 7.77 (m, 6H), 7.61 – 7.53 (m, 1H), 7.45 (tt, $J$ = 8.0, 3.9 Hz, 2H), 7.32 – 7.19 (m, 4H), 7.17 – 7.12 (m, 1H), 7.09 – 6.93 (m, 4H), 5.88 (ddt, $J$ = 16.2, 10.7, 5.4 Hz, 1H), 5.28 – 5.21 (m, 1H), 5.15 (dd, $J$ = 10.5, 1.6 Hz, 1H), 5.06 – 4.89 (m, 2H), 4.64 – 4.52 (m, 1H), 4.47 (d, $J$ = 5.4 Hz, 2H), 3.70 (dt, $J$ = 13.4, 5.4 Hz,

2H), 3.23 (t, *J* = 7.3 Hz, 2H), 2.68 (t, *J* = 7.3 Hz, 2H), 1.99 – 1.82 (m, 2H), 1.77 (q, *J* = 8.6 Hz, 2H).LCMS m/z $[M+H]^+$ Calcd for $C_{38}H_{39}N_3O_8P^+$ 696.2; Found 696.5.

**Compound I-33.** 1.2 g (1.73 mmol, 1 eq) of **I-32** was dissolved in 17 mL of EtOH, 347 μL (5.19 mmol, 3 eq) of ethylenediamine was added, and the reaction was heated for 78 °C for 2 h. Upon complete consumption of the starting material, monitored by LCMS, the reaction was allowed to cool to room temperature and solvent was removed under reduced pressure. After filtering precipitates, the filtrate was dried under reduced pressure, yielding a yellow oil that was taken to the next step without purification. The presence of product was confirmed by LCMS (m/z $[M+H]^+$ Calcd for $C_{30}H_{37}N_3O_6P^+$ 566.2; Found 566.4).

**Compound I-34.** Roughly 980 mg (1.73 mmol, 1 eq) of **I-33** was dissolved in 25 mL DCM. 2.15 grams (6.92 mmol, 4 eq) of N,N'-Di-Boc-1H-pyrazole-1-carboxamidine and 965 µL (6.92 mmol, 4 eq) of TEA were added to the solution and the reaction was allowed to stir overnight at room temperature. Upon consumption of the starting material (LCMS), solvent was removed under reduced pressure and the mixture was resuspended in EtOAc. Organic layer was subsequently washed x3 with each of the following: 1N HCl, $NaHCO_3$, and brine. It was then dried over anhydrous $Na_2SO_4$, filtered, evaporated under reduced pressure, and subjected to purification with combi-flash silica gel chromatography, yielding 585 milligrams (42%) of yellow oil. $^1H$ NMR (500 MHz, MeOD) δ 7.85 (ddt, *J* = 11.7, 8.3, 1.9 Hz, 2H), 7.61 (tt, *J* = 7.4, 1.4 Hz, 1H), 7.48 (tt, *J* = 6.4, 3.1 Hz, 2H), 7.32 – 6.98 (m, 9H), 5.89 (ddt, *J* = 16.2, 10.7, 5.4 Hz, 1H), 5.25 (dq, *J* = 17.3, 1.7 Hz, 1H), 5.15 (dd, *J* = 10.5, 1.6 Hz, 1H), 5.04 – 4.90 (m, 2H), 4.48 (t, *J* = 1.6 Hz, 1H), 4.47 – 4.40 (m, 2H), 3.46 – 3.34 (m, 2H), 3.24 (t, *J* = 7.3 Hz, 2H), 2.70 (td, *J* = 7.2, 2.6 Hz, 2H), 2.12 – 2.03 (m, 1H), 1.85 – 1.73 (m, 2H), 1.70 – 1.54 (m, 1H), 1.49 (d, *J* = 27.9 Hz, 18H). LCMS m/z $[M+H]^+$ Calcd for $C_{41}H_{55}N_5O_{10}P^+$ 808.4; Found 809.3.

**Compound I-35.** 250 mg (0.31 mmol) of **I-34** was cooled to 0 °C and then dissolved in approximately 2 mL of 33% HBr in AcOH. The reaction was allowed to run for 30 min, after which the solvent was removed under air while the vial was kept at 0 °C. After diluting the crude with 3 mL of 1:1 ACN:$H_2O$, it was subject to separation by reverse-phase HPLC. Desired fractions were lyophilized, yielding 127 mg (86%) of a white solid. $^1H$ NMR (500 MHz, MeOD) δ 7.95 (ddd, *J* = 12.3, 8.2, 1.4 Hz, 2H), 7.79 – 7.71 (m, 1H), 7.63 (td, *J* = 7.7, 4.4 Hz, 2H), 7.19 – 7.08 (m, 4H), 5.88 (ddt, *J* = 16.2, 10.6, 5.3 Hz, 1H), 5.24 (dt, *J* = 17.1, 1.7 Hz, 1H), 5.15 (dt, *J* = 10.5, 1.5 Hz, 1H), 4.46 (dt, *J* = 5.4, 1.6 Hz, 2H), 4.18 – 4.02 (m, 1H), 3.27 – 3.24 (m, 4H), 2.71 (td, *J* = 7.2, 3.4 Hz, 2H), 2.16 (dtd, *J* = 14.7, 11.1, 5.6 Hz, 1H), 2.01 – 1.88 (m, 2H), 1.87 – 1.78 (m, 1H). LCMS m/z $[M+H]^+$ Calcd for $C_{23}H_{32}N_5O_4P^+$ 474.2; Found 474.2.

**Compound I-36.** 127 mg of **I-35** (0.27 mmol, 1.1 eq) was dissolved in 4 mL DMF and 158 mg (0.245 mmol, 1 eq) of **I-29,** 103 mg (0.27 mmol, 1.1 eq) of HATU, and 163 µL (1.23 mmol, 5 eq) of 2,4,6-collidine were added to the solution and stirred for 2.5 h. Solvent was removed under $N_2$ and the mixture was purified by reverse-phase HPLC. Desired fractions were lyophilized, yielding 166 mg (61%) of a white solid. LCMS m/z $[M+H]^+$ Calcd for $C_{59}H_{75}N_9O_{10}P^+$ 1100.5; Found 1100.5.

**Compound I-37.** 20 mg (0.018 mmol, 1.5 eq) of **I-36** was dissolved in a solution of 90:5:5 TFA:DCM:TIPS and stirred at room temperature. After complete deprotection was observed (45 minutes), the crude was pipetted into a 15 mL Falcon tube. Precipitation was carried out with 12 mL of cold ether, the pellets were dissolved in 500 μL anhydrous DMSO and transferred to a 1.5 mL Eppendorf tube containing 9.0 mg (0.012 mmol, 1 eq) of sulfo-Cy5-NHS. Then, roughly 21 μL (0.12 mmol, 10 eq) of DIPEA was added, reaction vessel covered in Al-foil and shaken in an incubator overnight (Note: additional 1 eq of TSTU and 10 eq of DIPEA were added over the course of 24 h to reactivate any hydrolyzed sulfo-Cy5 in situ). The reaction was stopped after approximately 40 h and purified by reverse-phase HPLC. Desired fractions were collected and lyophilized, yielding 6.82 mg (41%) of a dark blue solid. LCMS m/z $[M+H]^+$ Calcd for $C_{68}H_{92}N_{11}O_{15}PS_2^+$ 698.8; Found 699.1.

**qABP 16.** 1.6 mg (0.0014 mmol, 0.5 eq) of $Pd(PPh_3)_4$ was quickly added to a 4 mL scintillation vial containing 4 mg (0.028 mmol, 1 eq) of **I-37** in dissolved in 1 mL anhydrous DMF. 10 μL of pyrrolidine was added, and the reaction was stirred for 30 min under $N_2$ atmosphere. Upon complete consumption of the starting material, the crude was purified by reverse-phase HPLC, desired fractions were lyophilized, redissolved in 1 mL of DMSO and added to a 1.5 mL Eppendorf tube containing 1.48 mg (0.0019 mmol, 0.7 eq) of QSY-21 NHS ester. Then, 4.8 μL (0.28 mmol, 10 eq) of DIPEA was added, and the reaction was allowed to stir for 1 hour, after which it was subjected to reverse-phase HPLC. Lyophilization of desired fractions yielded 1.65 mg (44%) of a dark blue solid. HRMS (Q-TOF) m/z: $[M]^{2+}$ Calcd for $C_{105}H_{121}N_{14}O_{17}PS_3^{2+}$ 988.8978; Found 988.8993 (Deconvoluted 1975.78).

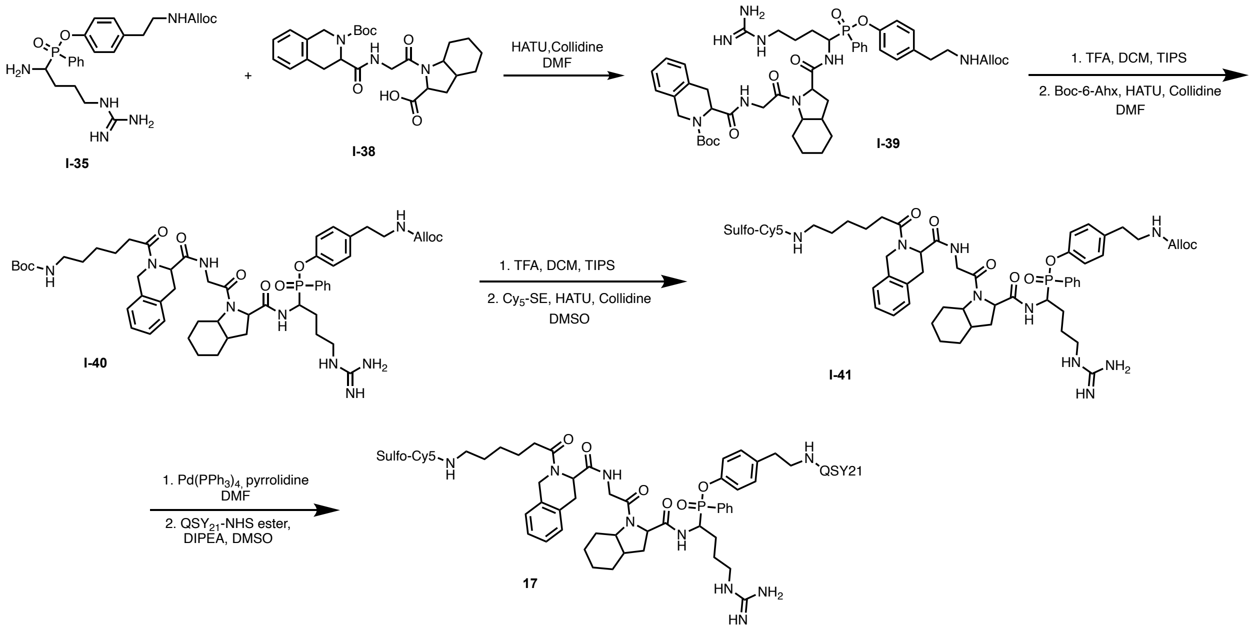


Scheme S7. Synthesis of phosphinate GzmA qABP **17**

**Boc-Tic-Gly-Oic-COOH (Compound I-38):** Tripeptide **I-38** was synthesized through solid-phase peptide synthesis following general procedure outlined above. LCMS m/z: $[M + H]^+$ Calcd for $C_{26}H_{36}N_3O_6^+$ 486.3; Found 486.0.

**Compound I-39.** 43.7 mg (0.092 mmol, 1.2 eq) of **I-35** was added to a 20 mL scintillation vial containing 37.4 mg (0.077 mmol, 1 eq) of **I-38** and 35 mg of HATU (0.092 mmol, 1.2 eq). Then 50.9 μL (0.385 mmol, 5 eq) of 2,4,6-collidine was added, and mixture was stirred for 4 h. Solvent was removed under air and the crude was subjected to purification by reverse-phase HPLC. Desired fractions were collected and lyophilized, yielding 42.2 mg (58%) of a white solid. LCMS m/z $[M+H]^+$ Calcd for $C_{49}H_{66}N_8O_9P^+$ 941.5; Found 941.2.

**Compound I-40.** 15.4 mg (0.016 mmol, 1 eq) of **I-40** was dissolved in 1 mL of a 9:1:1 TFA:DCM:TIPS solution and stirred at room temperature for 2 h. Excess reagents were removed under reduced pressure and mixture was resuspended in 1 mL of anhydrous DMF. 3.70 mg (0.016 mmol, 1 eq) of Boc-6-aminohexanoic acid, 6.1 mg (0.016 mmol, 1 eq) of HATU, and 10.6 μL (0.08 mmol, 5 eq) of 2,4,6-collidine were added, and the reaction was stirred at room temperature. At the 1 h mark, an additional 3 mg (0.008 mmol, 0.5 eq) HATU, 1.85 mg boc-6-aminohexanoic acid (0.008 mmol, 0.5 eq) and 5.3 μL (0.04 eq) 2,4,6-collidine were added and the mixture was stirred for another hour. The crude was concentrated under air, and diluted in 3 mL of 1:1 ACN:$H_2O$ solution before being purified by reverse-phase HPLC. Desired fractions were collected and lyophilized, yielding 10.6 (63%) mg of a white solid. LCMS m/z $[M+H]^+$ Calcd for $C_{55}H_{77}N_9O_{10}P^+$ 1054.6; Found 1054.8.

**Compound I-41.** 10.6 mg (0.01 mmol, 1 eq) of **I-40** was dissolved in 1.5 mL of a 9:1:1 solution of TFA:DCM:TIPS and allowed to stir for 1 hour. After removing solvents under vacuum, the crude was resuspended in 1.5 mL of anhydrous DMSO and transferred to an Eppendorf tube containing 7.54 mg (0.01 mmol, 1 eq) of Sulfo-Cy5 NHS ester. Then, 17.4 μL (0.1 mmol, 10 eq) of DIPEA was added, the reaction vessel was covered with Al-foil and allowed to stir at room temperature overnight. The crude was purified by reverse-phase HPLC, and desired fractions were lyophilized, yielding 3.41 mg (17%) of a blue solid. LCMS m/z $[M+H]^+$ Calcd for $C_{83}H_{108}N_{11}O_{15}PS_2^+$ 796.9; Found 797.4.

**qABP 17.** 2.74 mg (0.0017 mmol, 1 eq) of **I-40** was dissolved in 0.5 mL of anhydrous DMF and transferred to a 4 mL scintillation vial containing 0.98 mg (0.00085 mmol, 0.5 eq) of $Pd(PPh_3)_4$. Then, 5 μL of pyrrolidine was quickly added, and the vial was sealed and put under $N_2$ atmosphere. After 30 min, reaction was stopped, and the solvent was removed under reduced pressure. The crude was purified by reverse-phase HPLC, yielding 1.82 mg (71% yield, 0.0012 mmol, 1 eq) of a blue solid that was subsequently dissolved in 500 μL of anhydrous DMSO and transferred to an Eppendorf tube containing 0.94 mg (0.0012 mmol, 1 eq) of QSY21 NHS-ester. After adding 1.05 uL (0.006 mmol, 5 eq) of DIPEA, the reaction was stirred for 1 h at room temperature, after which the crude was diluted in 500 μL of a 1:1 ACN:$H_2O$ and purified by reverse-phase HPLC. Desired fractions were lyophilized, yielding 1.13 mg (43%) of a blue solid. HRMS (Q-TOF) m/z: $[M]^{2+}$ Calcd for $C_{120}H_{138}N_{14}O_{17}PS_3^{2+}$ 1086.9604; Found 1086.9632 (Deconvoluted 2171.91).

#### NMR Spectra

$^{1}$H NMR of Compound **1**

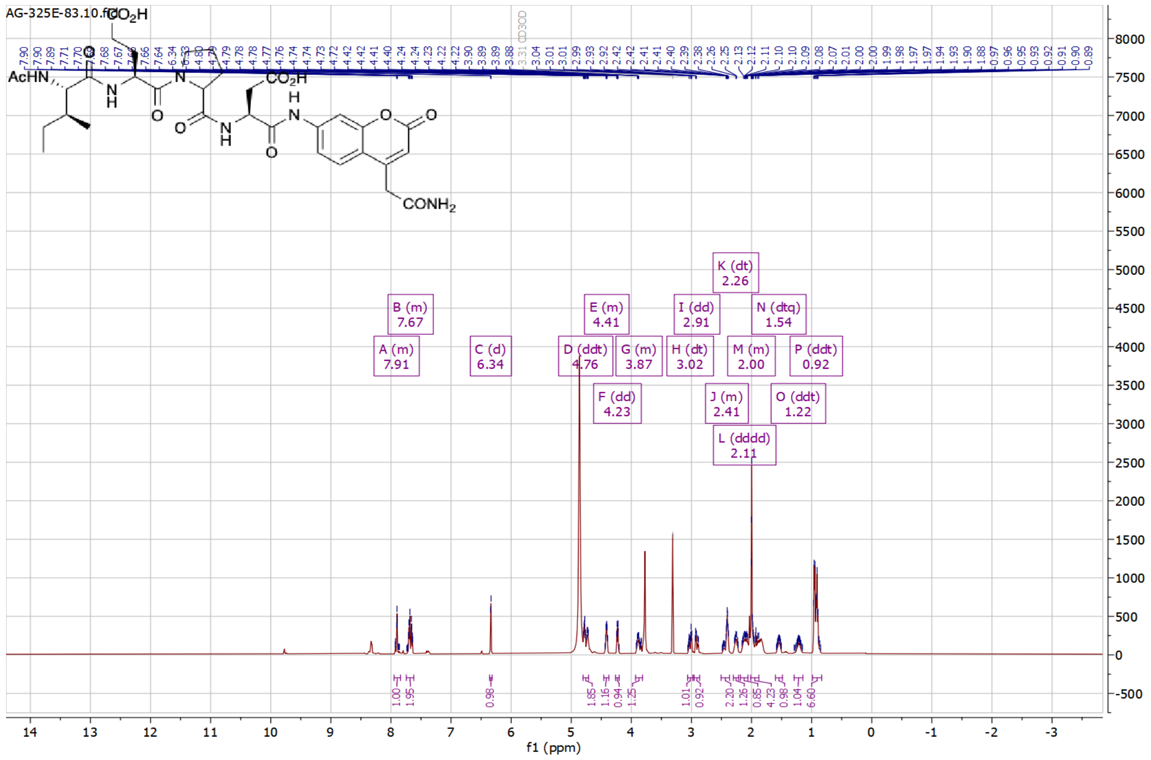

$^{1}$H NMR of Compound **2**

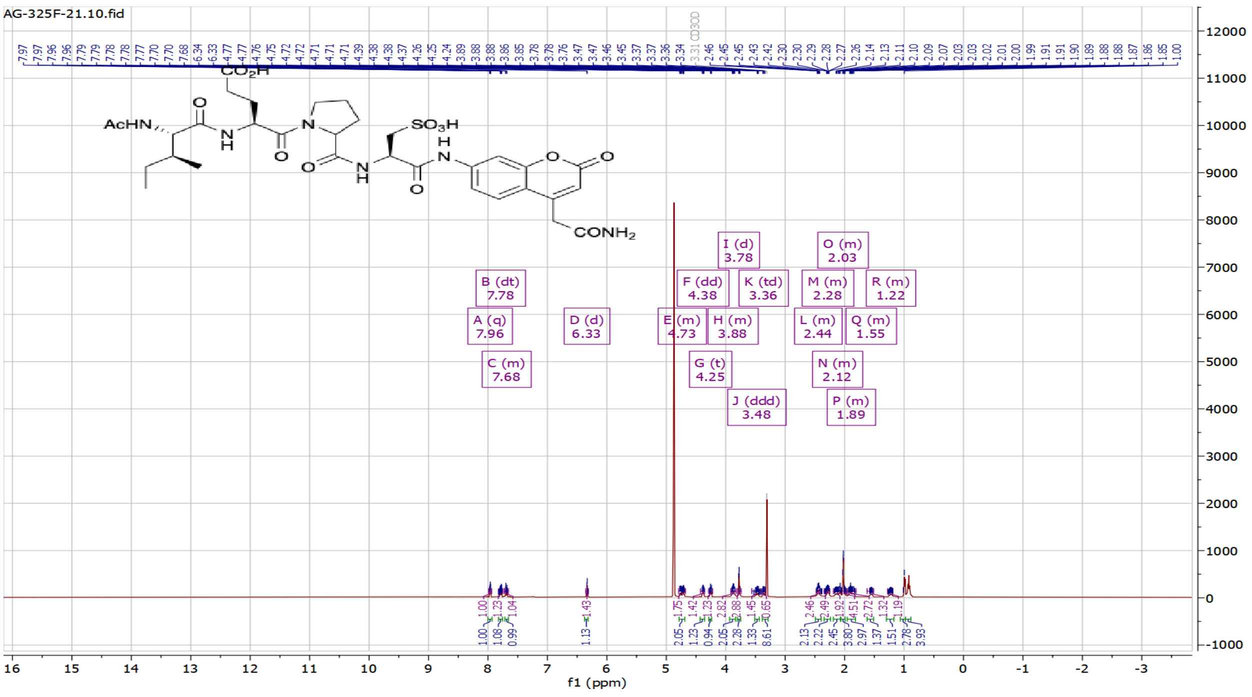

[1]H of Compound **I-1**

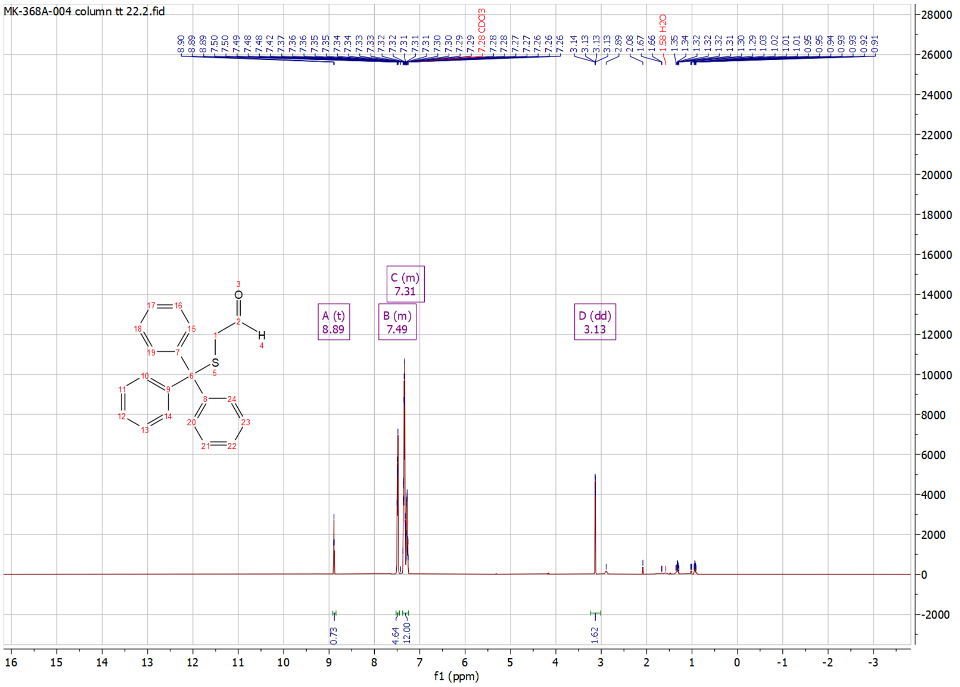


[1]H NMR of Compound **I-2**

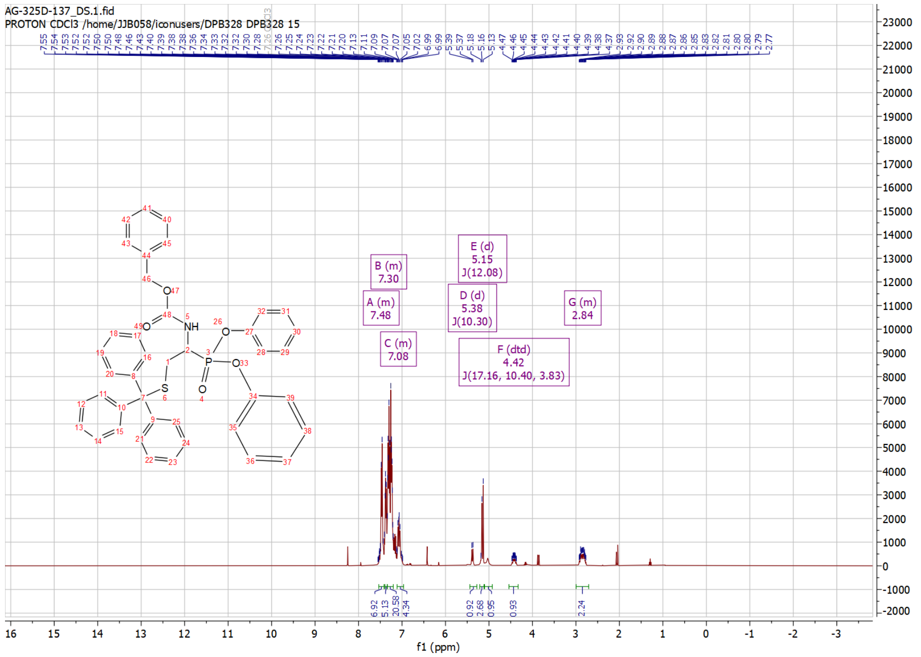

$^1$H NMR of Compound **I-3**

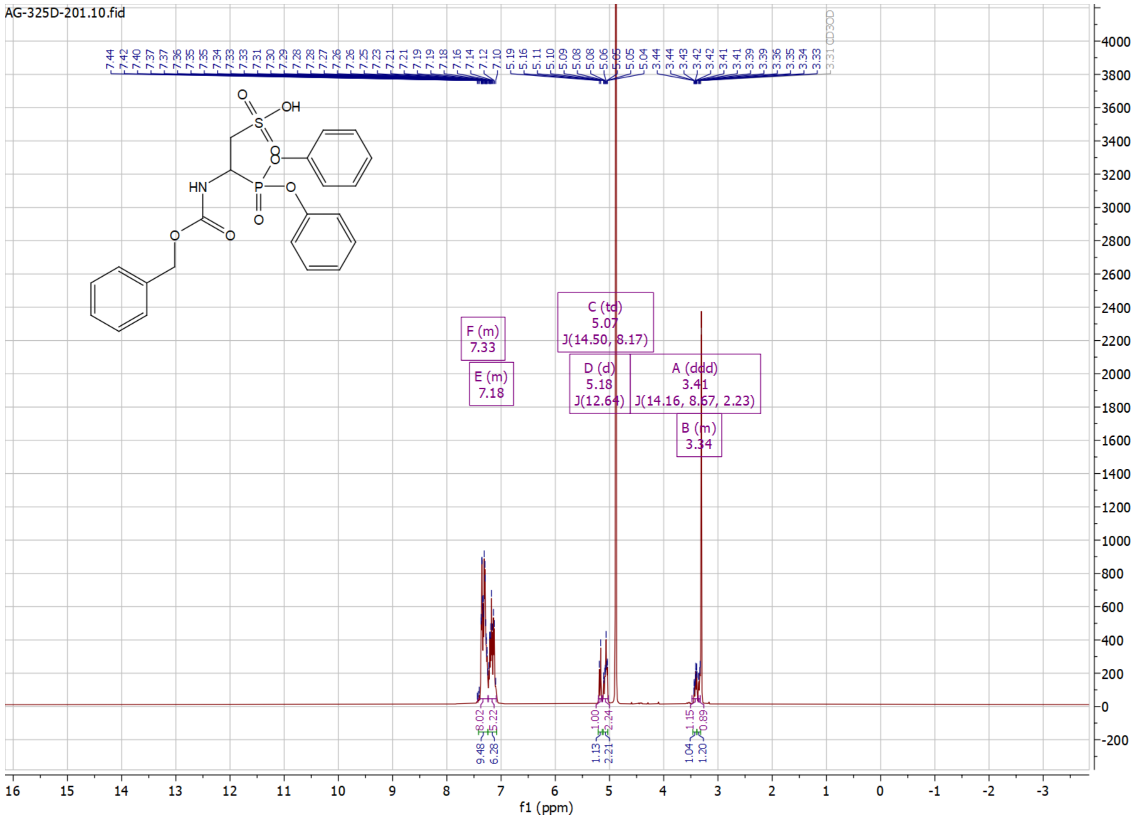


$^1$H NMR of Compound **I-4**

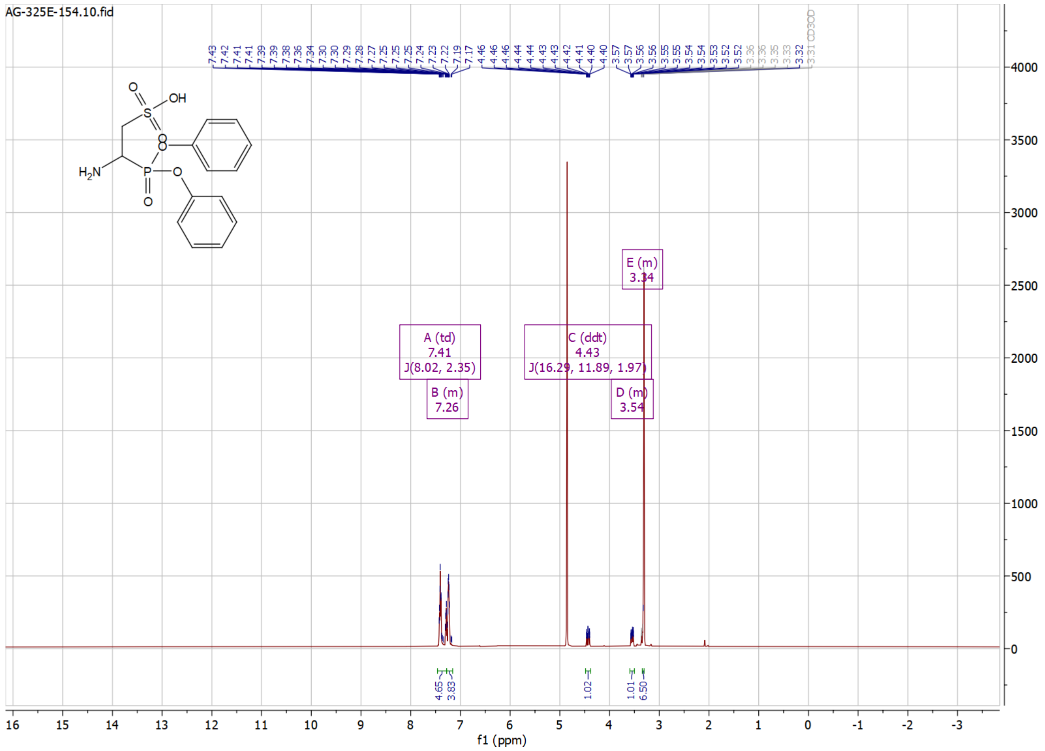

#### $^{1}$H NMR of Compound **6**

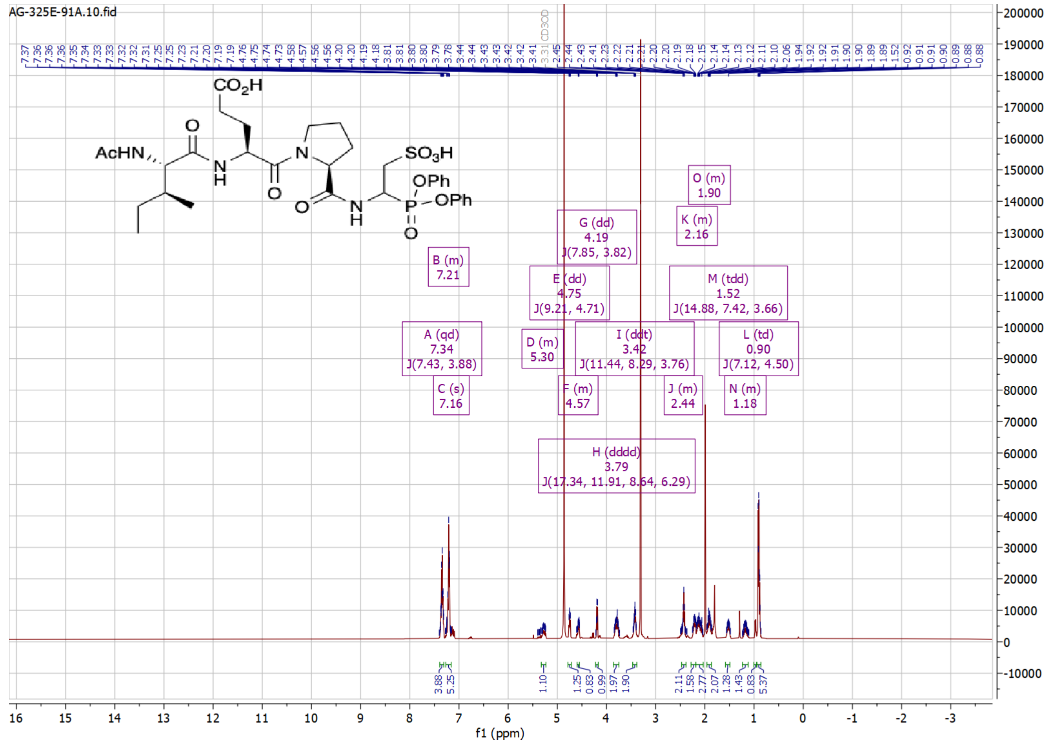


#### $^{1}$H NMR of Compound **I-5**

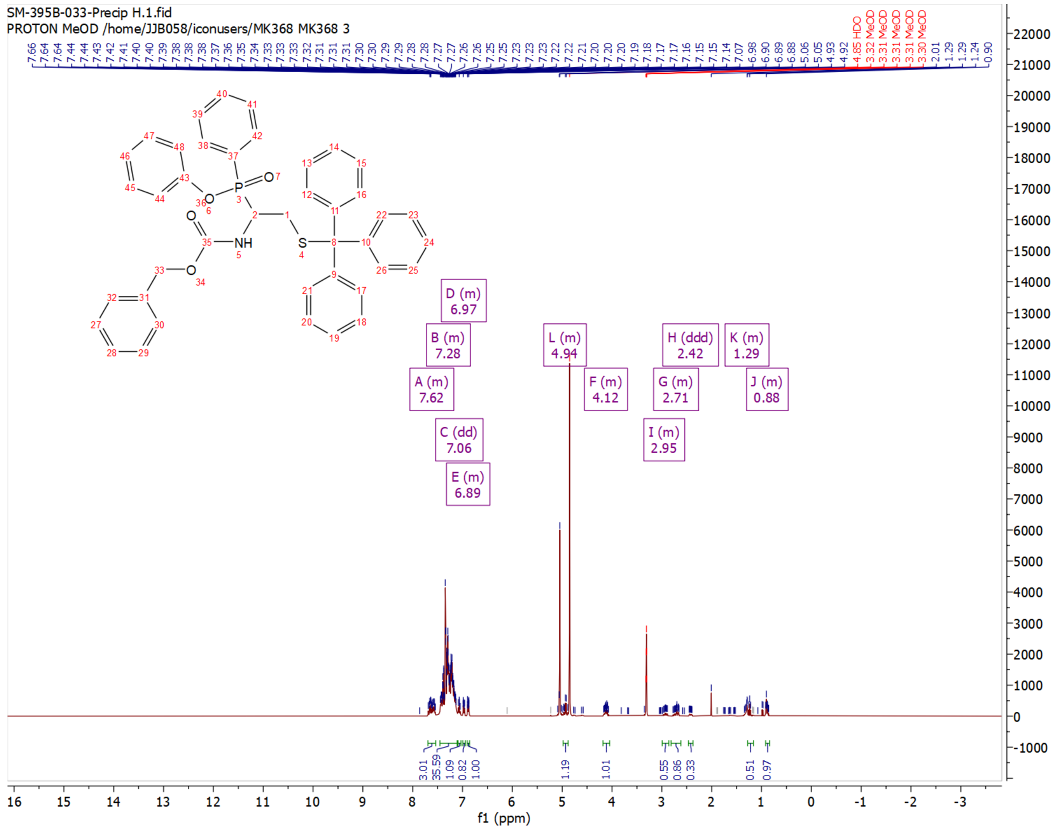

$^{1}$H NMR of Compound **I-6**

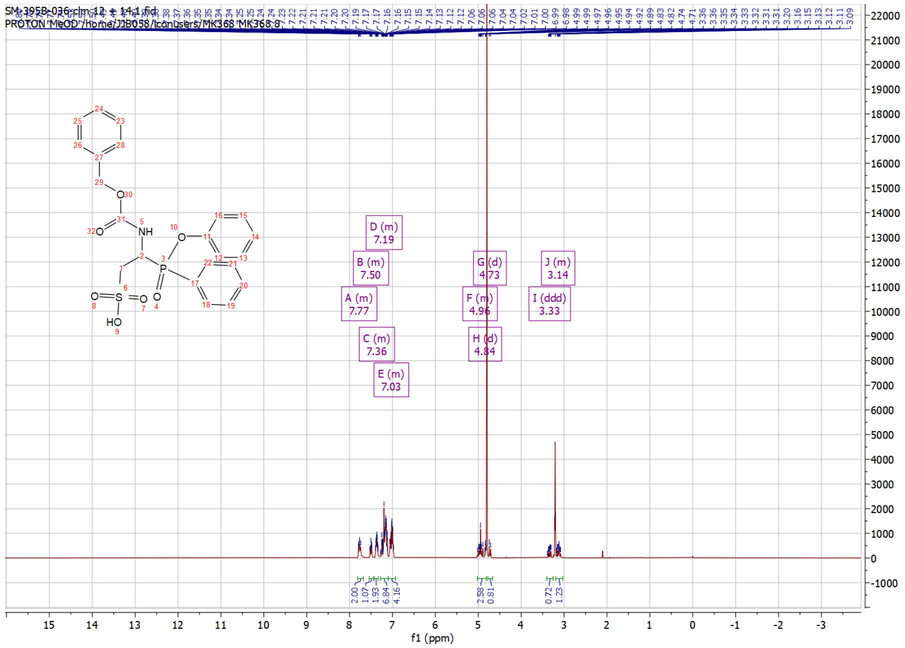


$^{1}$H NMR of Compound **I-7**

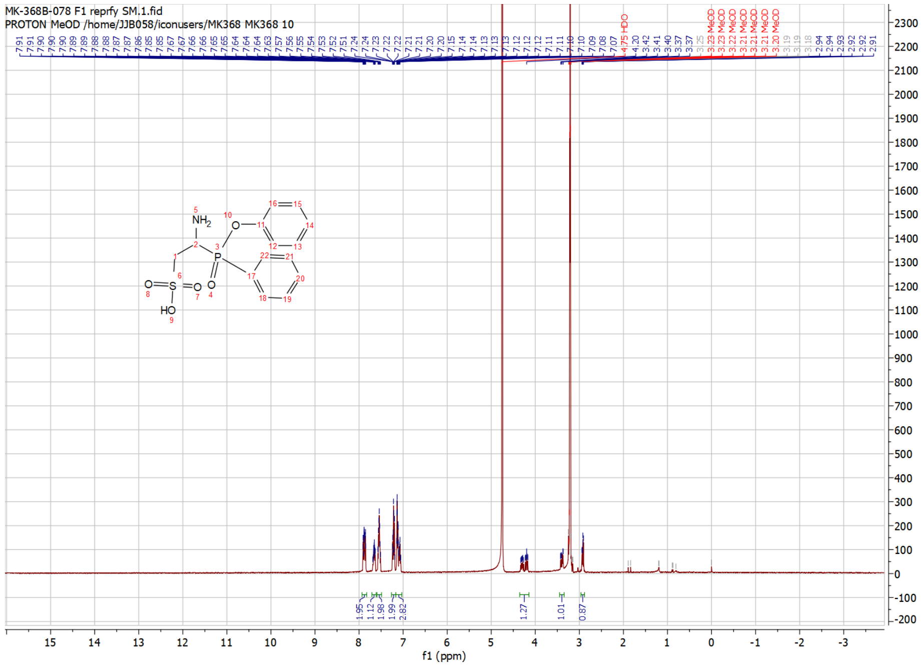

$^{1}$H NMR of Compound **7**

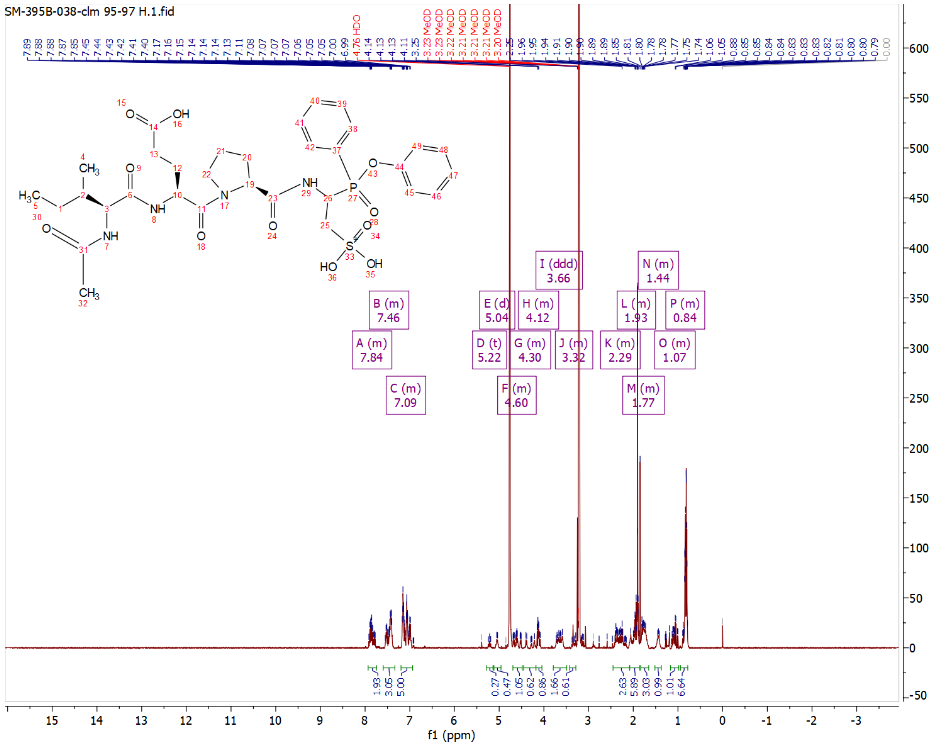

$^{1}H$ NMR of Compound **I-10**

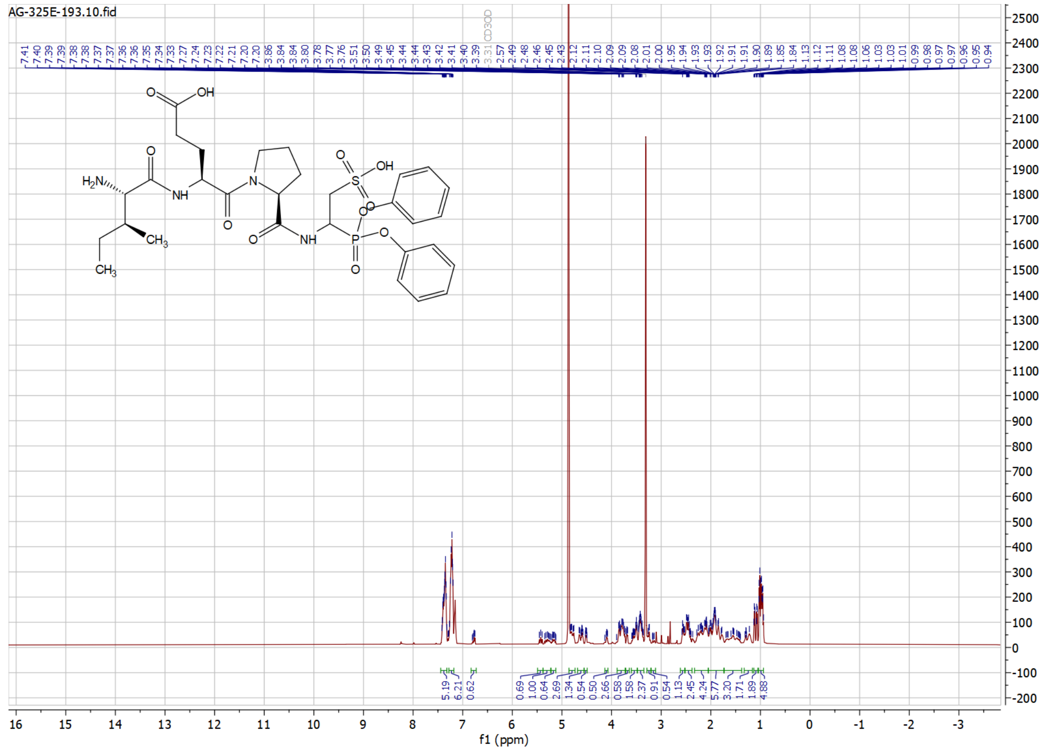

$^{1}H$ NMR of Compound **I-12**

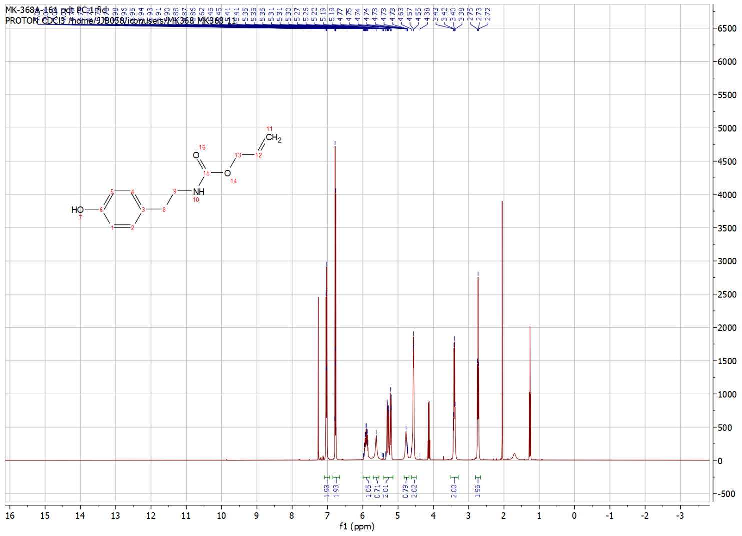

[1]H NMR of Compound **I-14**

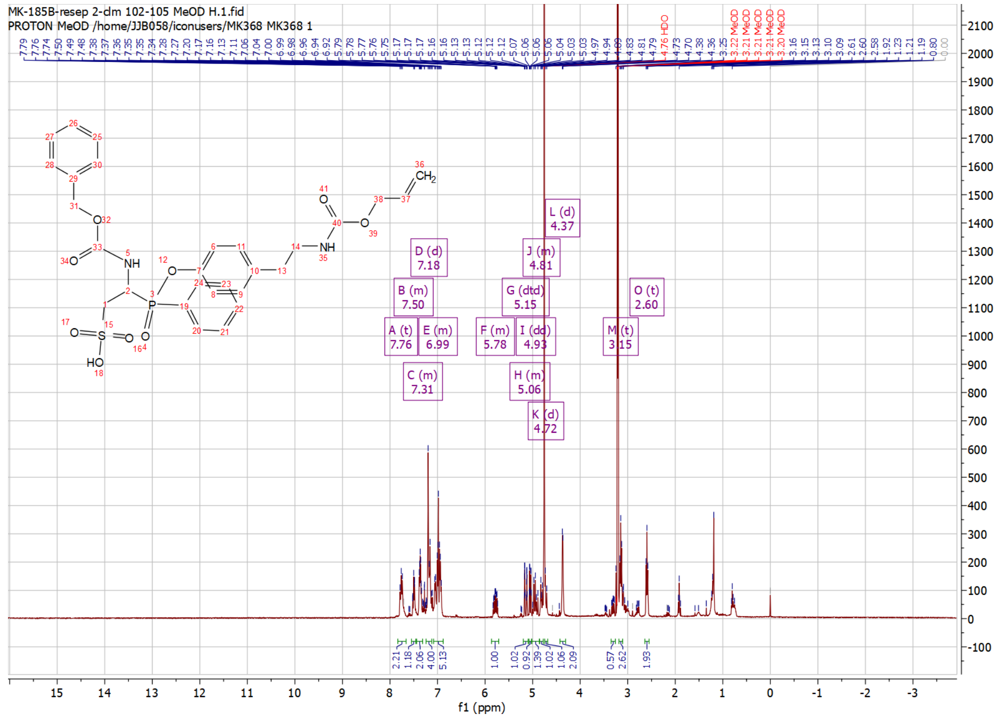

[1]H NMR of Compound **I-15**

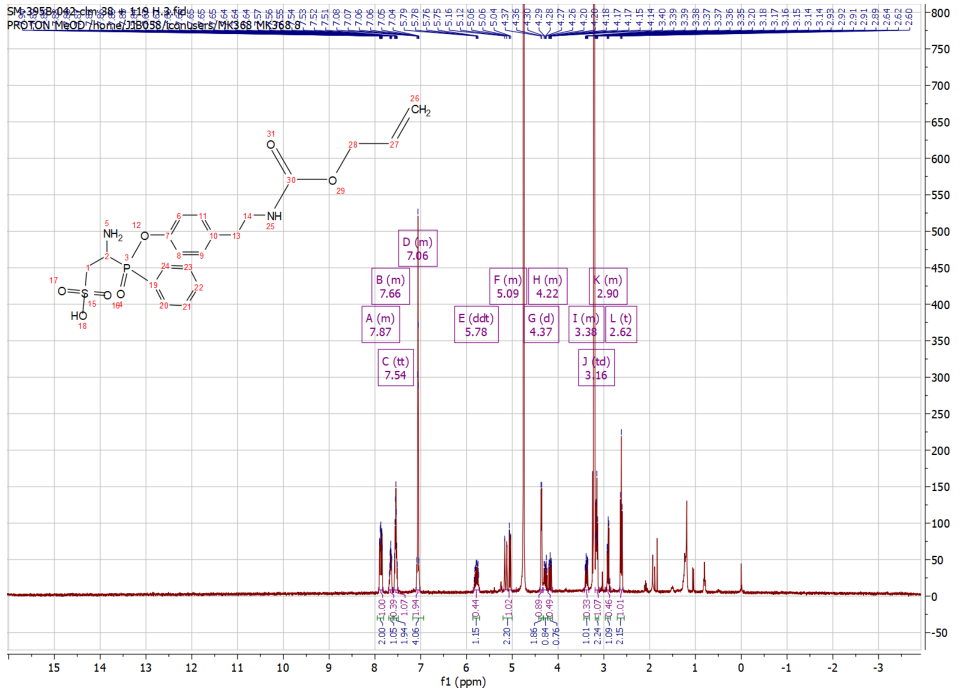

$^{1}$H NMR of Compound **I-18**

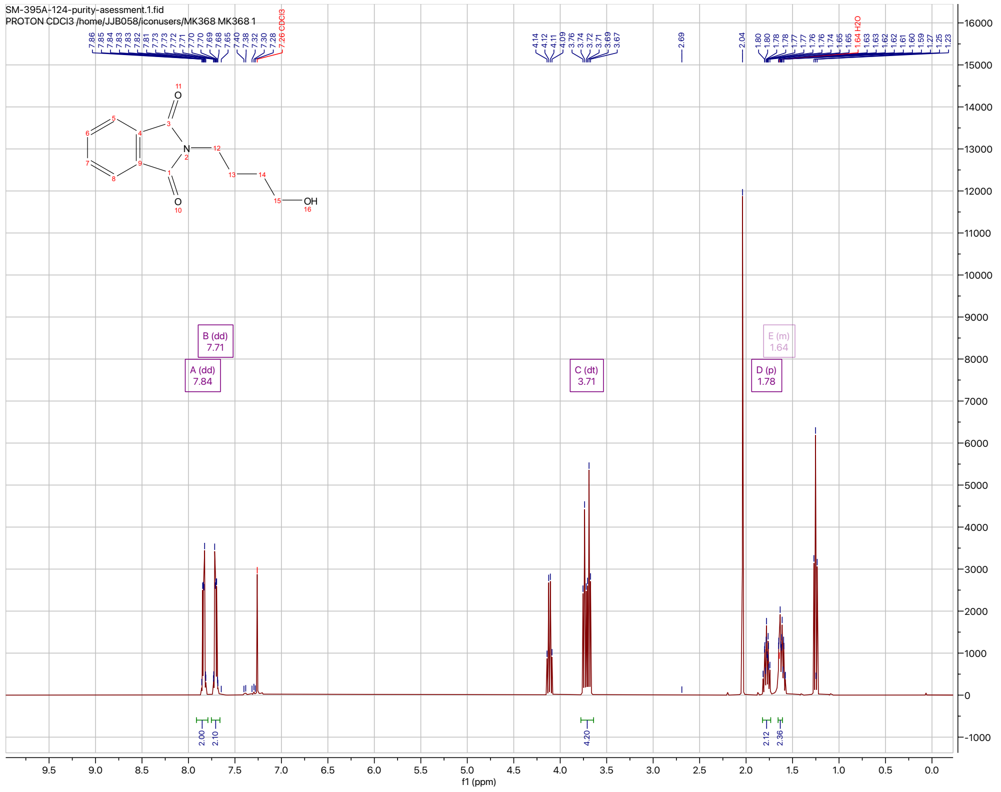

$^{1}$H NMR of Compound **I-19**

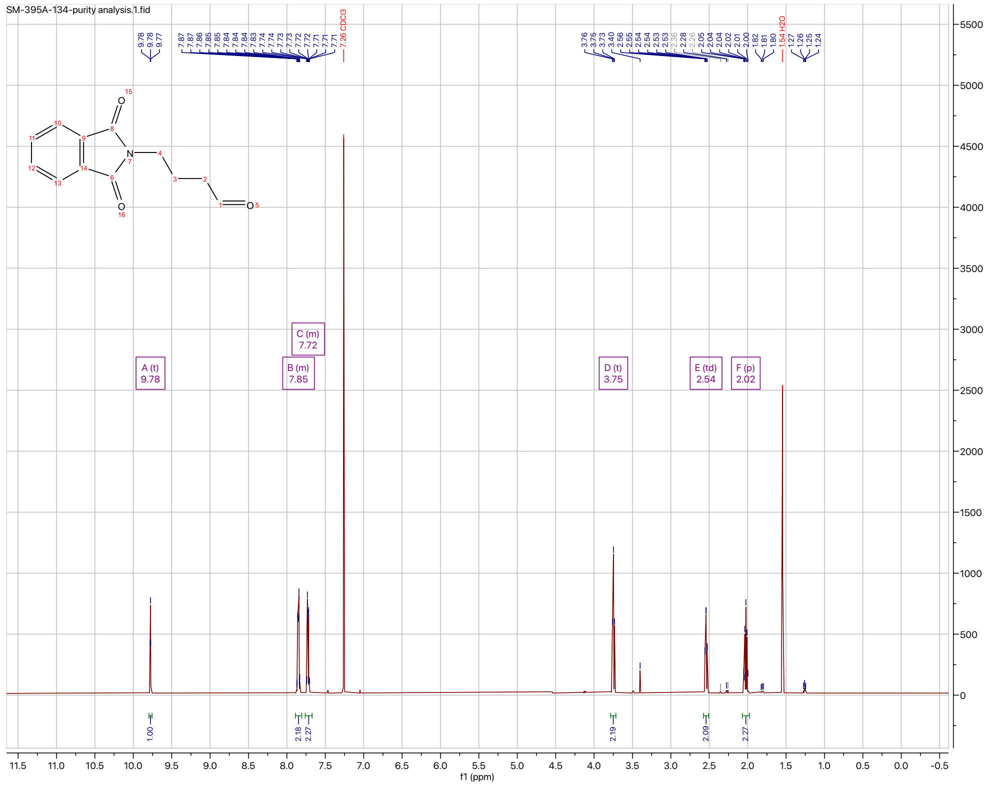

[1]H NMR of Compound **I-20**

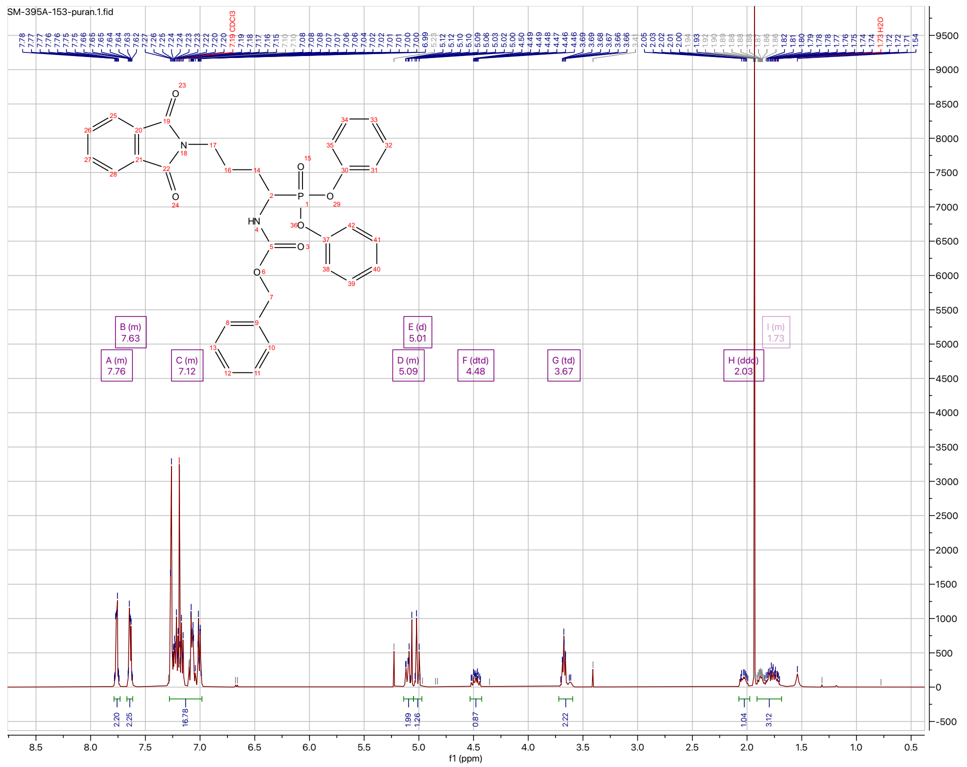

[1]H NMR of Compound **I-26**

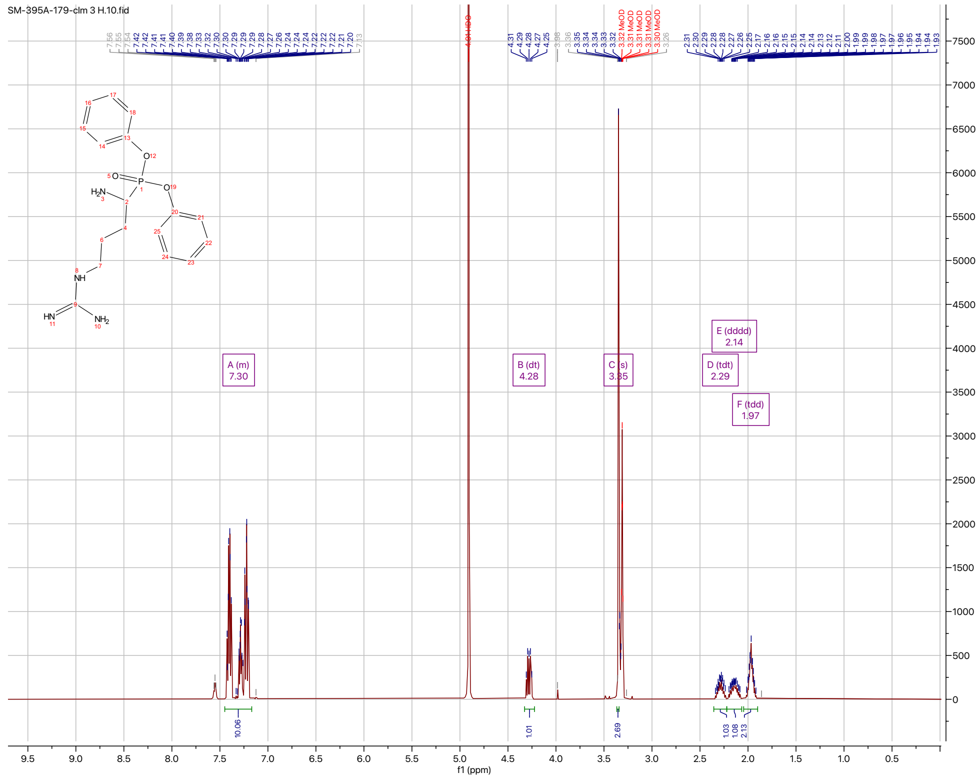

¹H NMR of Compound **I-27**

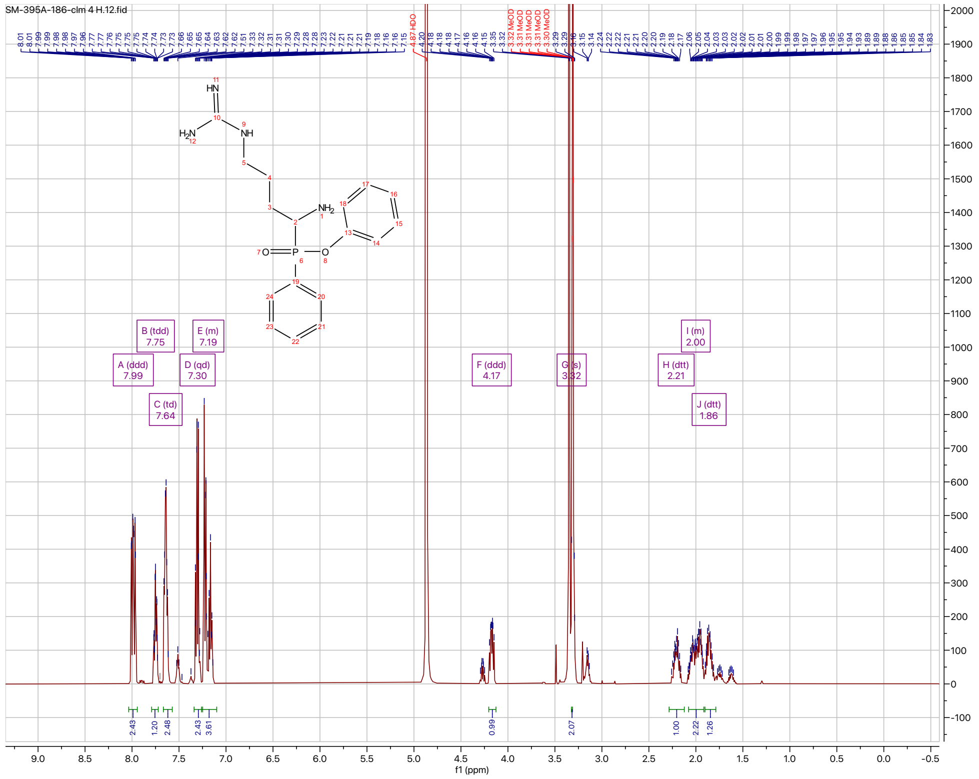

¹H NMR of Compound **I-33**

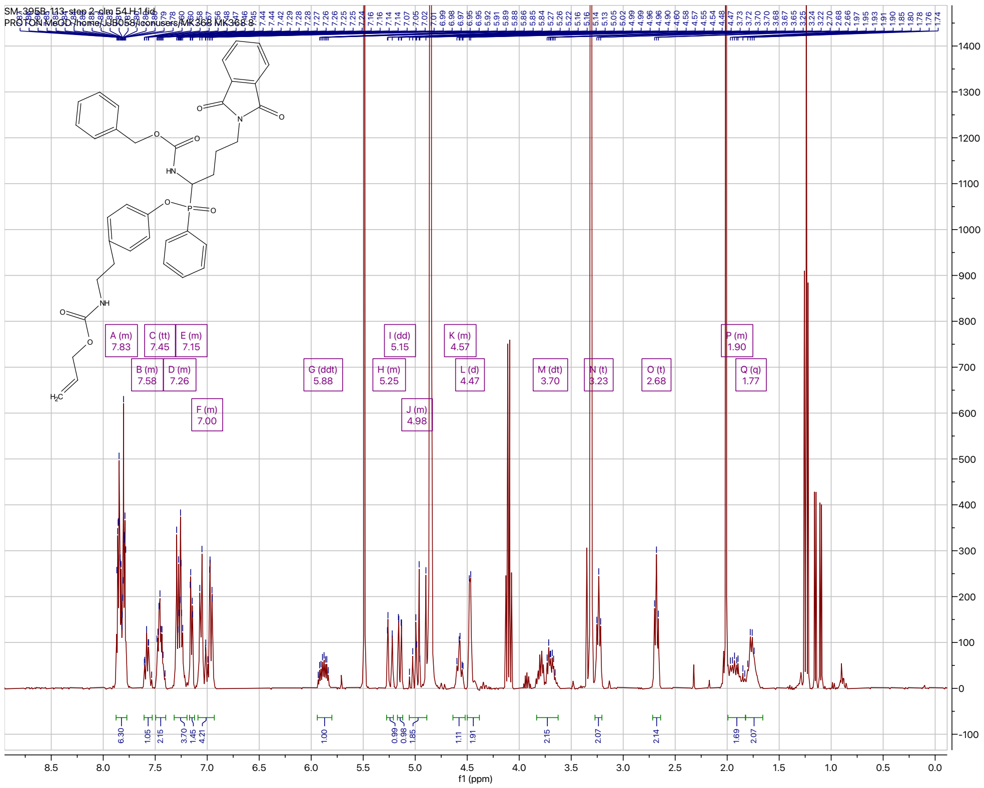

$^{1}$H NMR of Compound **I-34**

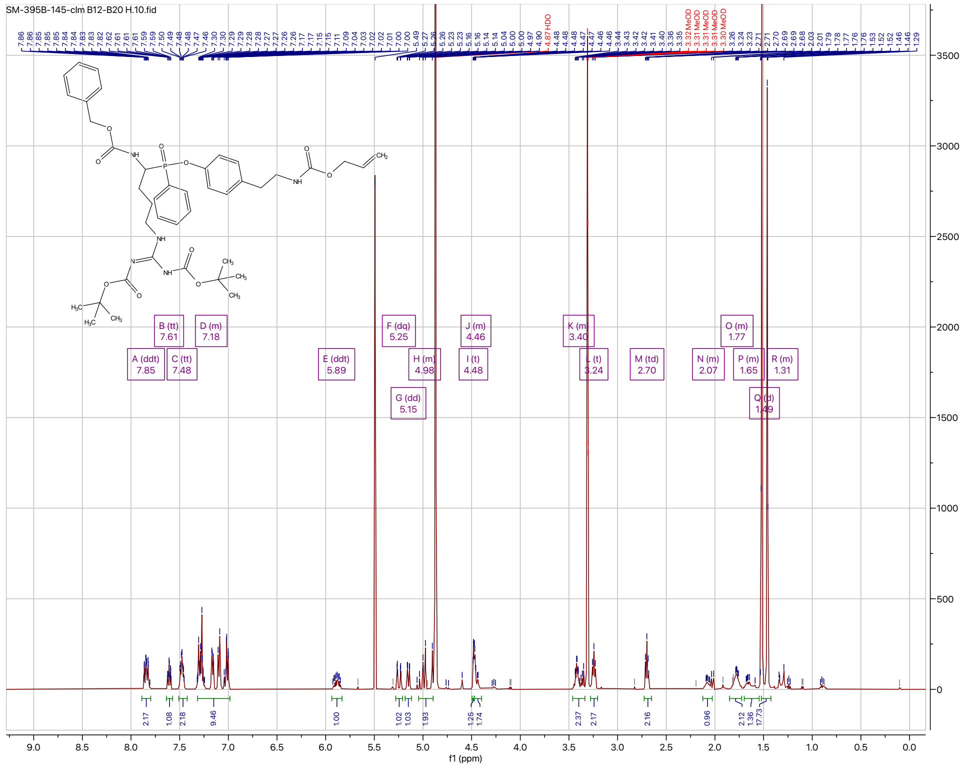

$^{1}$H NMR of Compound **I-35**

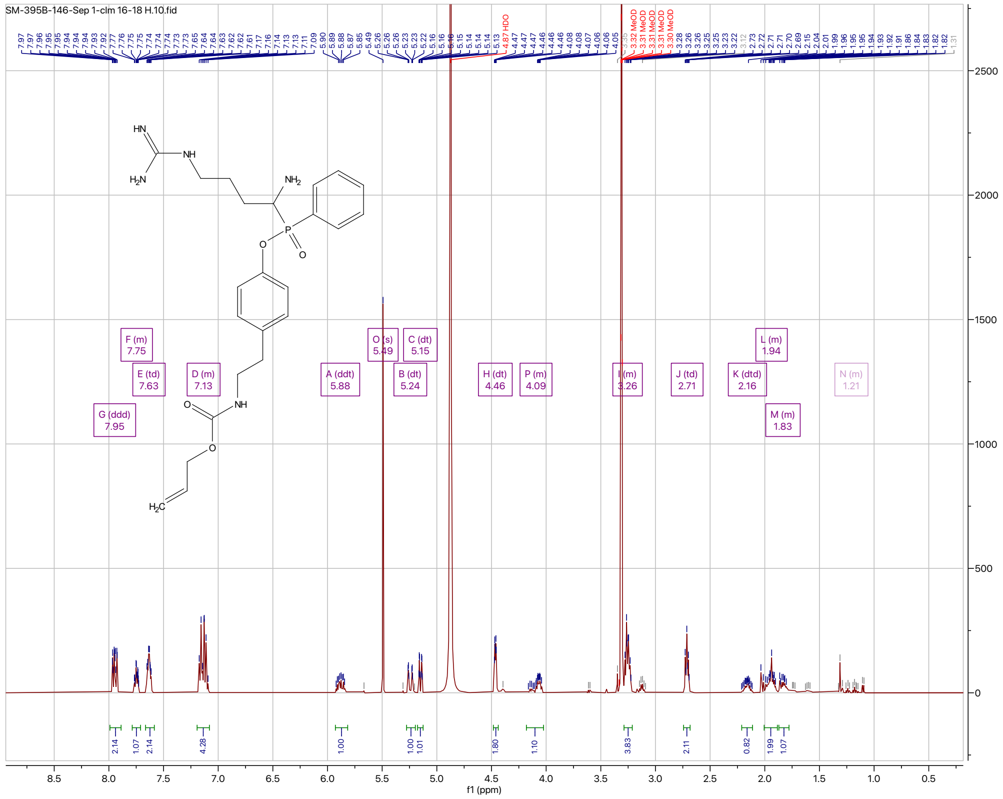

#### HRMS Data

HRMS of Compound **1**

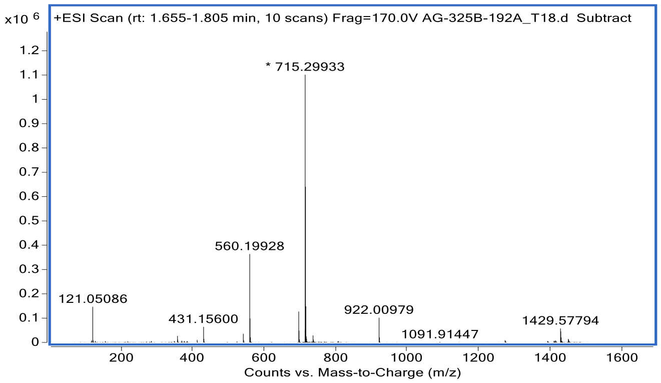


HRMS of Compound **2**

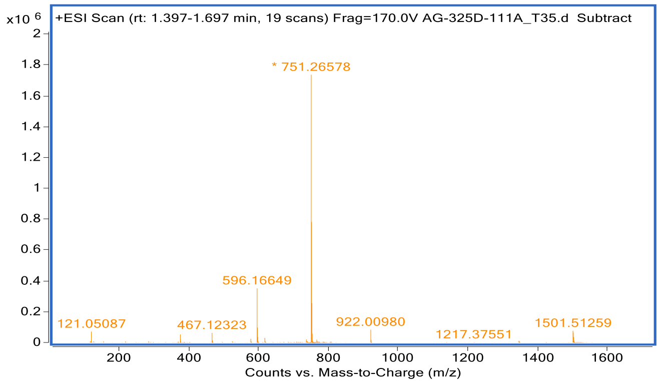


HRMS of Compound **3**

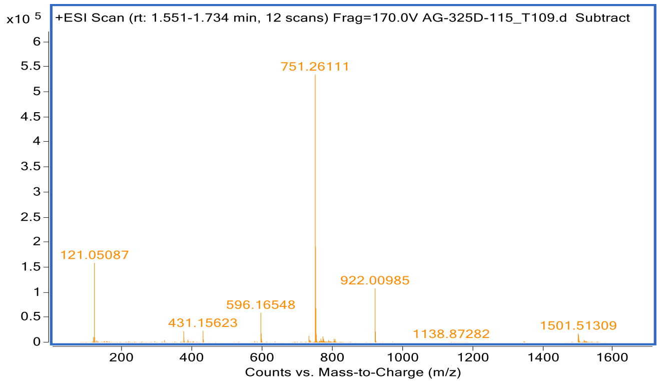

HRMS of Compound **4**

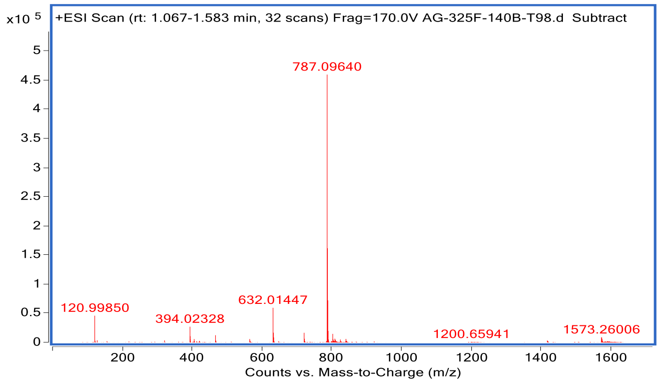


HRMS of Compound **5**

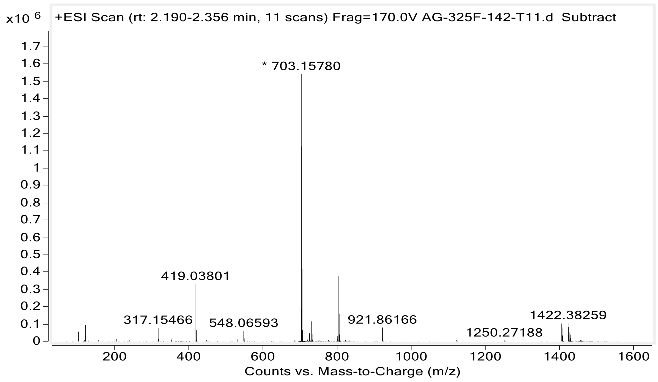


HRMS of Compound **6**

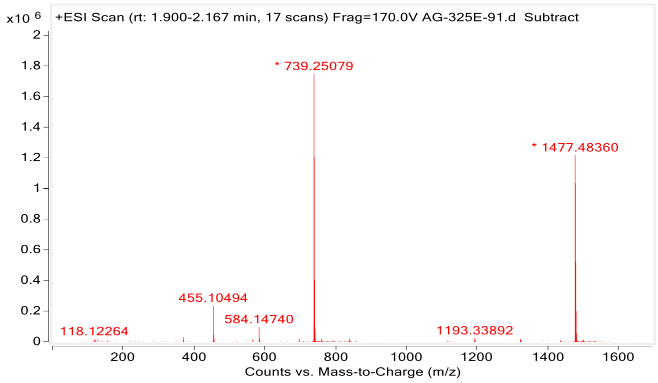

HRMS of Compound **7**

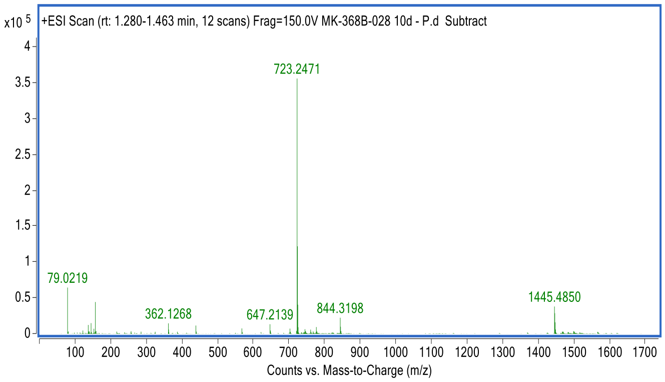


HRMS of **ABP 8**

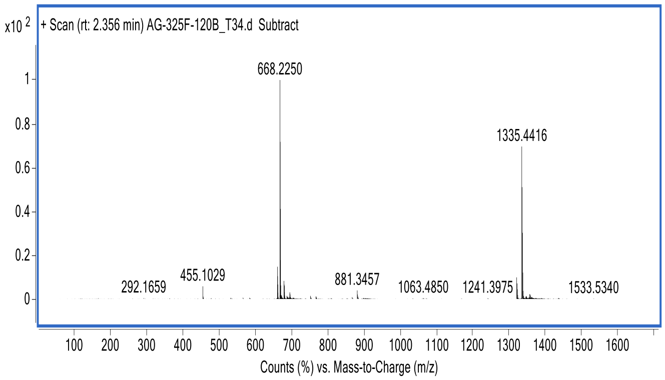


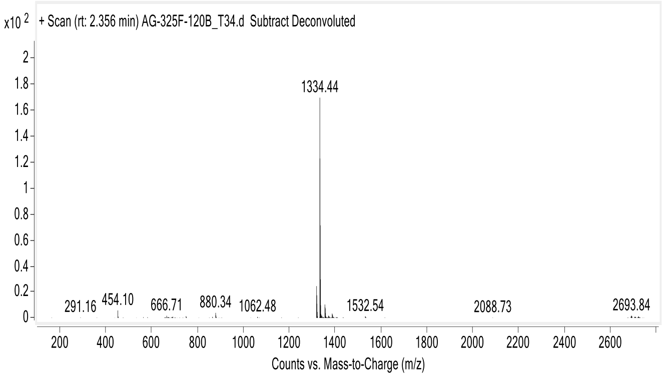

HRMS of **ABP 9**

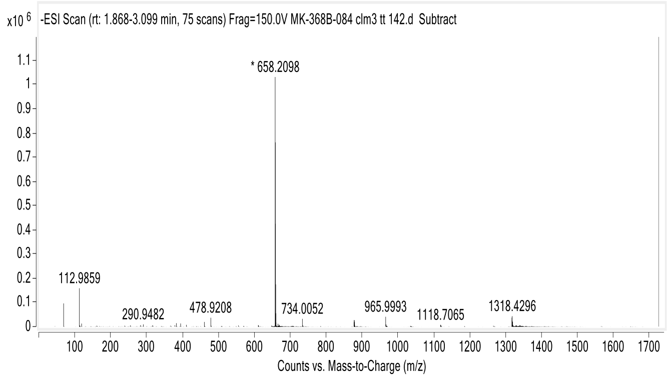
x10 6
-ESI Scan (rt: 1.868-3.099 min, 75 scans) Frag=150.0V MK-368B-084 clm3 tt 142.d Subtract
* 658.2098
112.9859
290.9482
478.9208
734.0052
965.9993
1118.7065
1318.4296
Counts vs. Mass-to-Charge (m/z)

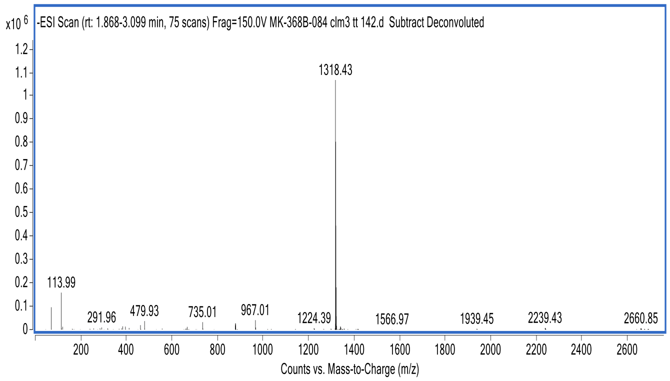
x10 6
-ESI Scan (rt: 1.868-3.099 min, 75 scans) Frag=150.0V MK-368B-084 clm3 tt 142.d Subtract Deconvoluted
1318.43
113.99
291.96
479.93
735.01
967.01
1224.39
1566.97
1939.45
2239.43
2660.85
Counts vs. Mass-to-Charge (m/z)

HRMS of **qABP 10**

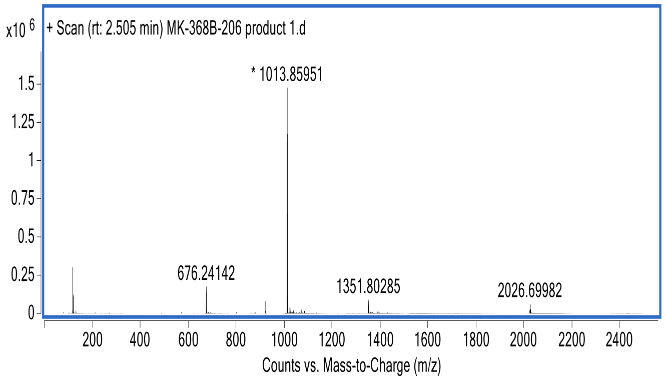


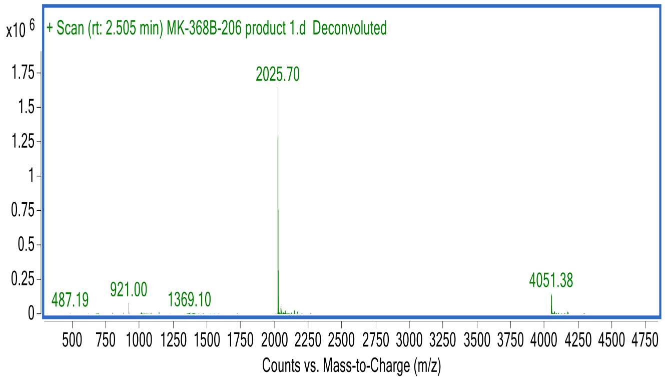


HRMS of Compound **11**

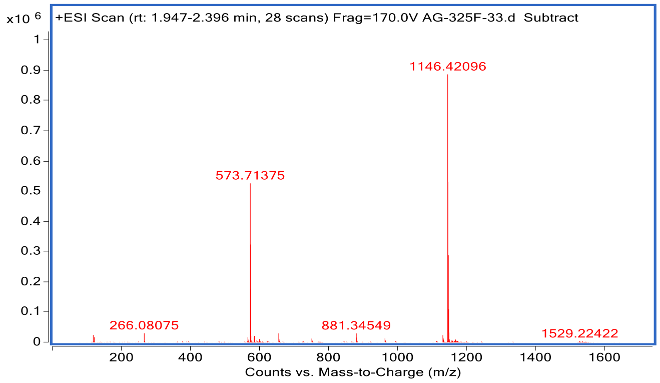

HRMS of Compound **12**

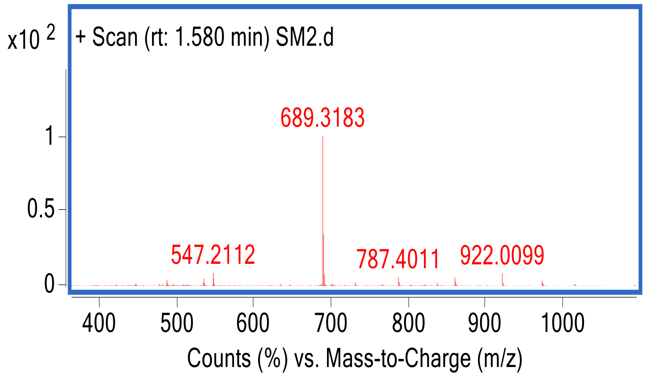


HRMS of Compound **13**

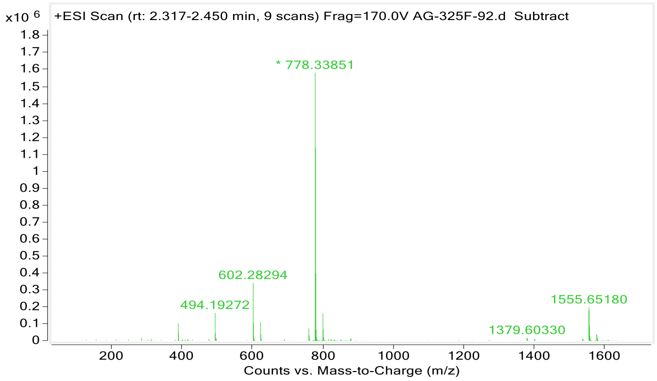

HRMS of **ABP 14**

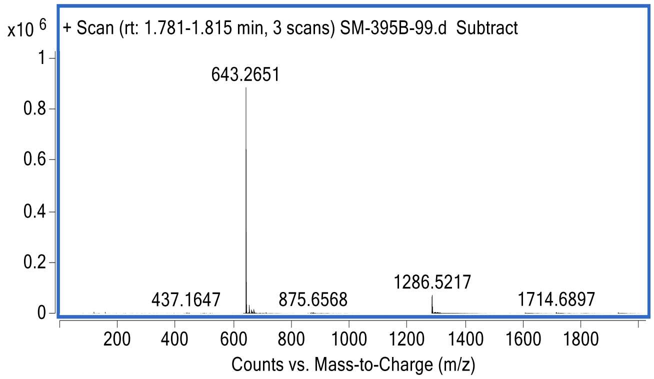


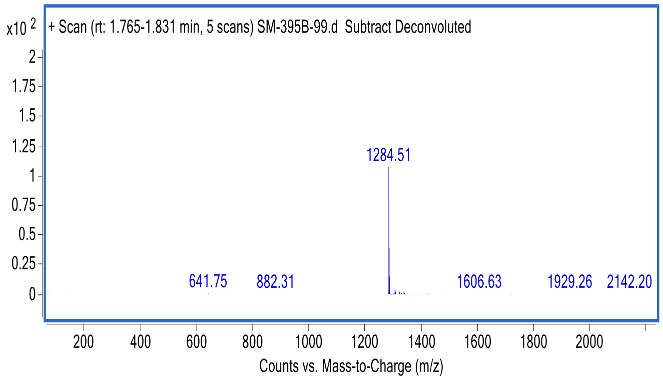

HRMS of **ABP 15**

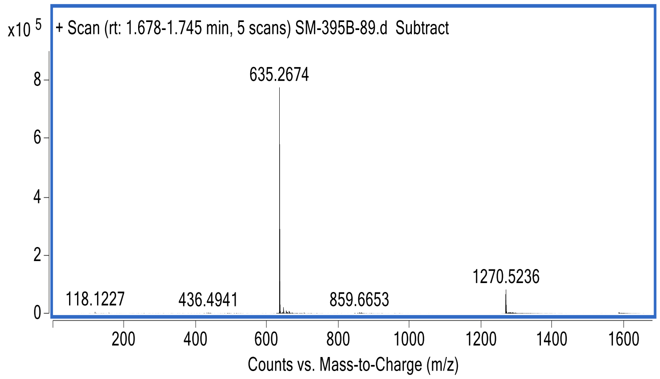


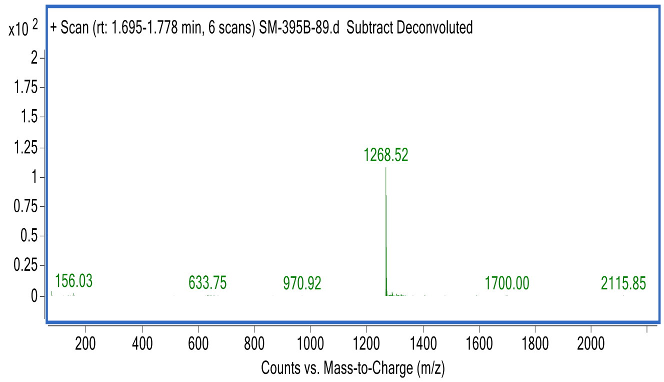

HRMS of **qABP 16**

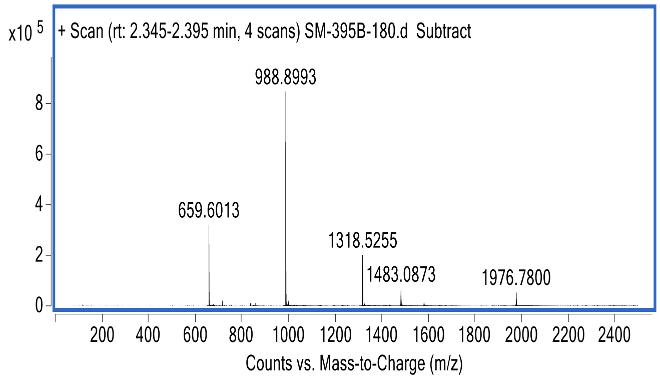


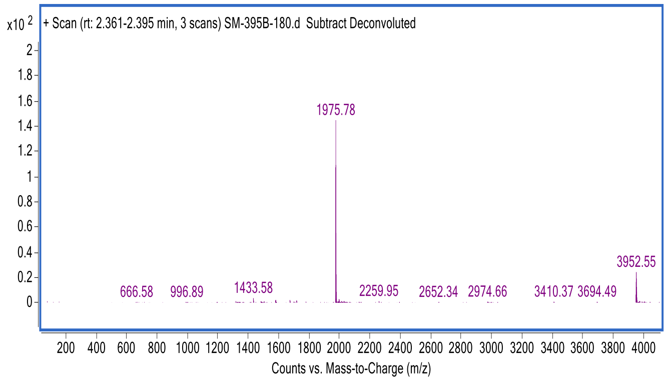

HRMS of **qABP 17**

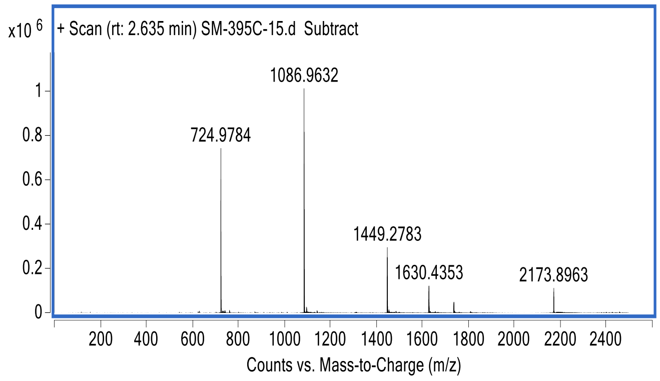


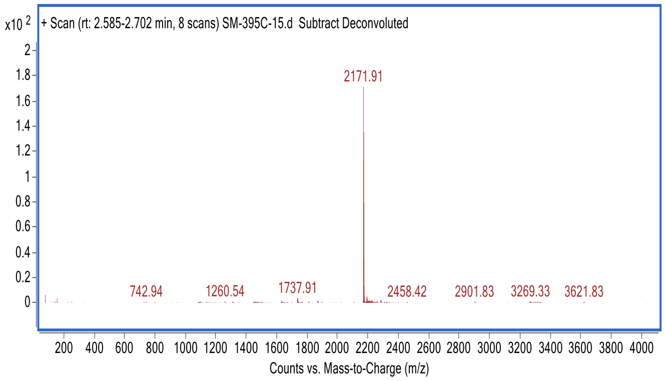
